# A unified model of aneuploid karyotype dynamics

**DOI:** 10.64898/2026.02.20.707056

**Authors:** Mathieu Hénault, Lisa M. Wood, Lydia R. Heasley

## Abstract

Aneuploidies—whole-chromosome copy number imbalances arising from nondisjunction—underlie numerous congenital and somatic disorders, but unlike many other disease-causing variants, they can readily revert back to euploidy through subsequent errors of the same type. The extent to which this inherent plasticity impacts the stability and persistence of aneuploid karyotypes in populations remains poorly understood, a gap in knowledge that continues to limit our understanding of aneuploidy-driven disease incidence, penetrance, and progression. To assess how reversion shapes aneuploid population dynamics, we developed a budding yeast system to systematically measure the rates at which aneuploidies arise and revert, and quantify the relative fitness differences between these karyotypic states. We integrated these data into a computational framework encompassing the broad physiological range of aneuploid karyotype dynamics captured in our experiments. The resulting models reveal that canonical reversion (*i.e.*, subsequent nondisjunction) occurs rarely, conferring a negligible effect on the population dynamics of most chromosomal aneuploidies. However, our models also identified that the reversion dynamics of some chromosomes—those that revert at extremely high rates—were more consistent with an alternative mechanism in which nondisjunction and reversion are directly coupled. Whole-genome sequencing and live-cell microscopy demonstrates that this coupled mechanism is facilitated by unresolved intermolecular linkages that disrupt chromosome segregation, leading to chromosome breakage and recombination-mediated repair over subsequent cell divisions. Collectively, this work establishes a unified model of aneuploid population genetics and expands our perspective of the diverse, and chromosome-specific mutational mechanisms shaping genome architecture.

## Introduction

The functional relevance of any novel genetic variation depends not only on the phenotypic effect it induces, but on its capacity to persist in the population. For most variants, persistence is primarily determined by the rates at which they arise (μ) and their relative fitness (ω). Yet the variants most likely to exert the strongest phenotypic effects—whole-chromosome copy number variations (*i.e.*, aneuploidies)—are additionally constrained by a third factor: reversion. Arising through mitotic or meiotic nondisjunction events, aneuploidies can impart profound physiological changes to cells by simultaneously altering the dosage of hundreds to thousands of genes. Aneuploidies underlie numerous pathological states and adaptive processes, ranging from constitutional diseases like Down syndrome (Bull 2020), to tumorigenesis (Weaver and Cleveland 2008; Turajlic et al. 2019), to the emergence of antimicrobial drug resistance (Selmecki et al. 2009). But these aneuploid states are inherently unstable in dividing populations; each subsequent mitosis carries a risk of nondisjunction, providing aneuploid cells with opportunities to revert to euploidy or acquire new aneuploid karyotypes (Kohanovski et al. 2024; Yona et al. 2012; Tan et al. 2013; Cromie et al. 2017; Hose et al. 2025; Gilchrist and Stelkens 2019; Gerstein and Berman 2015). As such, irrespective of any advantageous effects conferred by an aneuploid state, these phenotypes are ultimately only as stable as the underlying aneuploid genotype.

Aneuploidy reversion has been documented in numerous experimental systems (Tan et al. 2013; Heasley and Argueso 2022; Hose et al. 2025; Reid et al. 2008) and is generally described by a two-step sequential mutational framework predicated on the assumptions that a cell must first become aneuploid (step 1, A) before reverting (step 2, R), and that both aneuploid formation and reversion occur stochastically. Within this framework, models differ in their assertions about the likelihood of reversion. Since A and R both constitute nondisjunction of the same chromosome, the probability of R occurring could be expected to be equivalent to that of A (*i.e.,* μ^A^ = μ^R^), referred to here as the equal-likelihood model. By contrast, if A alters genome stability (Sheltzer et al. 2011; Heasley et al. 2020), the probability of subsequent nondisjunction events including R may be increased or decreased (*i.e.,* μ^A^ ≠ μ^R^), referred to here as the unequal-likelihood model. Yet, which of these models most accurately captures observed dynamics of reversion, and the extent to which this relationship shapes population-level karyotypic diversity remains poorly understood.

Empirically disentangling the effects of μ^A^, μ^R^, and ω on the observed frequency of an aneuploid karyotype has proven particularly challenging, since each determinant influences estimation of the others. For example, previous studies reporting the rates of aneuploidy in diploid yeast did not account for the effects of ω and μ^R^ on the aneuploid cell counts observed, potentially confounding the derived estimates of μ^A^ (Heasley et al. 2020; Heasley and Argueso 2022; Sharp et al. 2018; Kumaran et al. 2013). Likewise, since the relative fitness (ω) of aneuploid cells is often lower than euploids, estimates of aneuploidy-associated fitness effects may be substantially distorted if the emergence of revertants (μ^R^) is not considered (Hose et al. 2020; Dutcher et al. 2024; Zhu et al. 2016, 2012; Rojas et al. 2024; Torres et al. 2007; Pompei and Cosentino Lagomarsino 2023). Most recently, μ^R^ for three chromosomal aneuploidies in yeast was estimated with a system that accounted for fitness differences between the aneuploid and revertant states, but precluded parallel estimation of μ^A^ for the aneuploidies tested (Hose et al. 2025).

As it stands, a unified model incorporating the inherent ephemerality of aneuploid karyotypes into the complex interplay of factors governing their persistence in populations has not been established. Thus, our ability to anticipate the occurrence, stability, and functional implications of these important mutations remains fundamentally limited. To address this gap in knowledge, we systematically investigated aneuploidy reversion with an experimental system capable of integrating all major determinants of aneuploid frequency (μ^A^, ω, μ^R^). This approach allowed us to construct a comprehensive, quantitative model of aneuploid population dynamics and in doing so to identify and characterize a previously unidentified, non-canonical mutational mechanism that directly links the nondisjunction and rapid reversion of specific chromosomal aneuploidies.

## Results

### A genetic system to investigate aneuploidy dynamics in populations

Although chromosome loss is typically lethal in haploids, it can be tolerated in diploids (Heasley et al. 2020; Heasley and Argueso 2022; Peter et al. 2018; Sampaio et al. 2020). The resulting aneuploid state—called monosomy—can produce diverse effects on cellular physiology and fitness (Gerstein and Berman 2015; Weaver and Cleveland 2008; Sheltzer and Amon 2011; Gilchrist and Stelkens 2019; Heasley et al. 2020; Heasley and Argueso 2022), and once established, can persist for many generations. However, monosomic cells can also revert to euploidy by acquiring a second copy of the remaining homologous chromosome, generating a genotype called uniparental disomy (UPD) (Heasley and Argueso 2022; Reid et al. 2008; Engel 1993). Although UPDs restore a balanced chromosomal copy number, they result in whole-chromosome homozygosis and are, themselves, associated with numerous recessive genetic disorders (Engel 1993; Andersen and Petes 2012; Reid et al. 2008; Heasley et al. 2020). Given the important physiological implications of both monosomy and UPD, we focused our systematic analysis of aneuploidy frequency to this karyotypic pair using a tractable budding yeast (*Saccharomyces cerevisiae*) system.

To facilitate selective recovery of cells harboring monosomy or revertant UPD of each of the 16 yeast chromosomes, heterozygous diploid yeast strains were engineered as follows: for each pair of homologous chromosomes, two *CAN1* gene cassettes were inserted at loci on each arm of one homolog and a violacein pigment operon (Chuang et al. 2018) was inserted into the other homolog at a position allelic to one of the *CAN1* cassettes (Fig. 1A). Loss of *CAN1* expression confers resistance to the toxic arginine analog canavanine (CAN^R^), and this double-cassette architecture enriches for recovery of CAN^R^ clones arising from loss of the *CAN1*-marked chromosome while reducing those that result from copy-neutral loss of heterozygosity (LOH), segmental deletion, or point mutation (Heasley et al. 2020; Sampaio et al. 2020). When populations of these parental strains were pregrown in nonselective conditions and then plated onto canavanine-containing media, spontaneously arising CAN^R^ colonies developed and resolved into two major phenotypic classes—light purple (parental-like) or dark purple—reflecting the copy number difference of the pigment operon between monosomic and revertant UPD clones. In most backgrounds, dark CAN^R^ colonies were also larger, a phenotype consistent with euploidy revertants having higher predicted ω than monosomes (Fig. 1B) (Hose et al. 2025).

**Figure 1.**
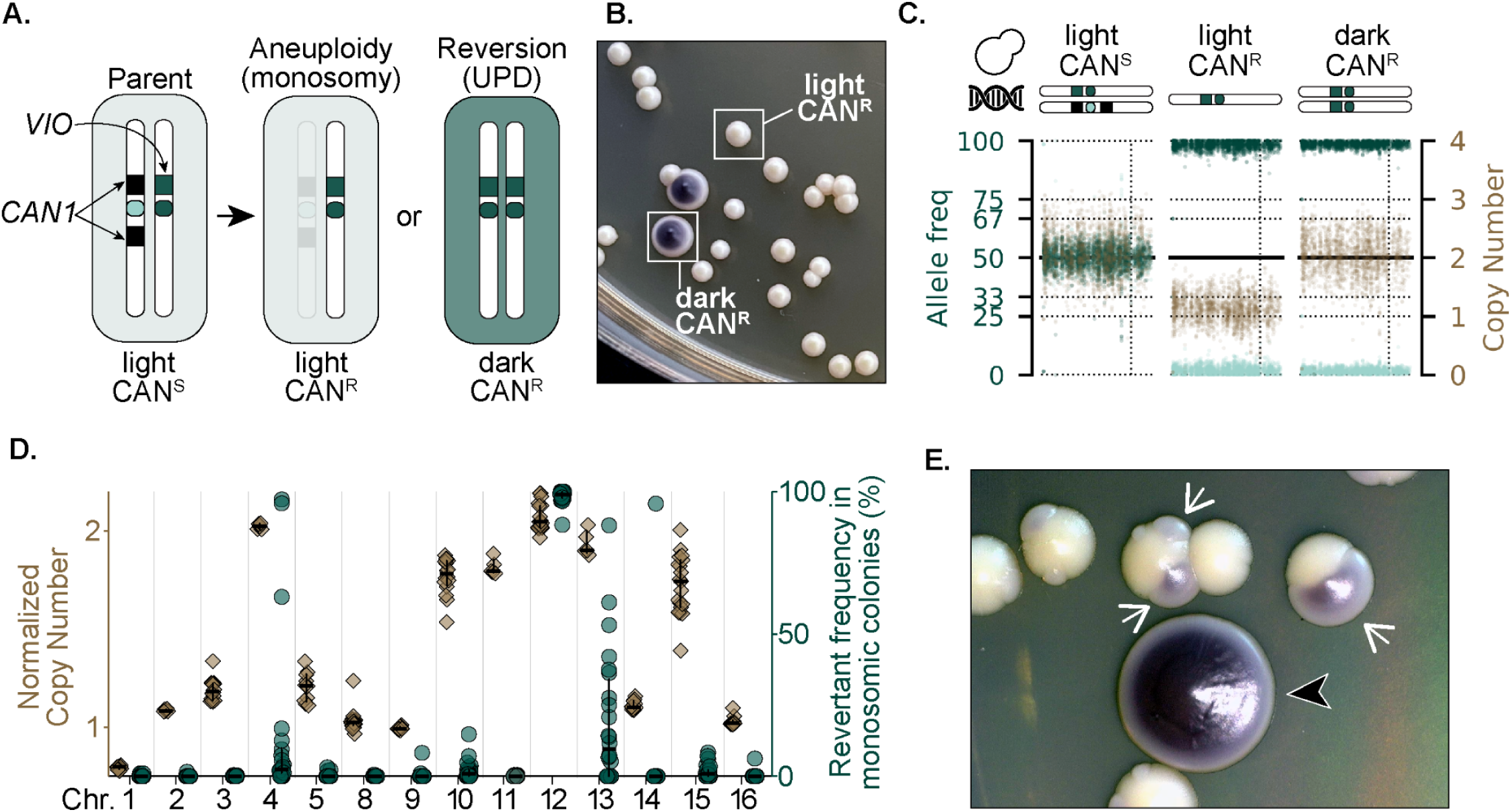
A new genetic reporter system reveals variations in reversion frequency across chromosomes. **A.** A schematic describing the genotypic and phenotypic outcomes reported by the *CAN1*-VIO selectable yeast system. **B.** A representative image of light and dark CAN^R^ colonies derived from the Chr10 strain. **C.** WGS data signatures associated with the parental (diploid) strain, light CAN^R^ (monosomic) colonies, and dark CAN^R^ (revertant UPD) colonies. Each dot denotes a heterozygous allelic position; colors denote normalized depth of coverage (brown), and the allele frequencies of the *CAN1*-marked chromosome (light green) and *VIO*-marked chromosome (dark green). **D.** Per-chromosome quantifications of the normalized copy number estimates inferred from WGS analysis of light CAN^R^ clones (brown diamonds) and the revertant frequency derived from the monosome replating tests (green circles). **E.** A representative image of the sectoring phenotype displayed by the CAN^R^ derivatives of the chromosome strains that display higher levels of revertant mosaicism, in this case, Chr10. Black arrowhead denotes a revertant colony identified at day 2 of the test, white arrowheads denote sectors that emerged within light colonies later in the experimental timecourse.

Whole-genome sequencing (WGS) of independent light and dark CANᴿ clones confirmed the accuracy of this selectable system (Fig. S1, Supplemental Table 2). Light CANᴿ clones predominantly showed a monosomic signature of the predicted chromosome—evidenced by whole-chromosome homozygosity and reduced read coverage—and dark CANᴿ clones primarily exhibited a UPD signature—evidenced by whole-chromosome homozygosity and diploid-level coverage (Fig. 1C). This analysis also revealed recurrent, chromosome-specific, reversion patterns: whereas the affected chromosome in monosomic clones of Chr1, Chr2, Chr3, Chr5, Chr8, Chr9, Chr14, and Chr16 usually had the expected normalized copy number near 1, all monosomic clones of Chr4, Chr10, Chr11, Chr12, Chr13, and Chr15 had a normalized copy number nearer to 2 (Fig. 1D, brown diamonds). This suggested that, despite displaying a light color phenotype, the clones of this latter group were mosaic, with sizable revertant subpopulations already present at the time of sequencing. We validated the variable revertant mosaicism inferred from WGS by directly measuring the frequency of revertant cells in monosomic populations. Light CAN^R^ colonies isolated from independent canavanine plates were resuspended and diluted to single-cell density on rich media plates. After 2 days, dark revertant colonies were counted; total population size was quantified after 2 additional days, to accommodate the slower growth of monosomes (Fig. 1E). WGS confirmed that the dark colonies recovered from these monosome-replating tests harbored the predicted revertant UPD genotype. In agreement with the WGS analysis, revertants were absent or very rare in populations of Chr1, Chr2, Chr3, Chr5, Chr8, Chr9, Chr10, Chr14, Chr15, and Chr16, but prevalent in populations of Chr4, Chr12, and Chr13 (Fig. 1D, green circles). In both assessments of revertant frequency, Chr12 was a striking outlier; every sequenced light CAN^R^ clone showed a penetrant signature of UPD, and every replated population was almost entirely fixed by revertants (average revertant frequency, 99%). At the experimental endpoint of these replating tests, we noted variations in colony appearance further reflecting the diverse reversion dynamics of different chromosomes: though dark colonies quantified after 2 days were large and fully purple (Fig. 1E, black arrowhead), many smaller monosomic colonies developed dark-purple sectors, indicating that reversions had occurred post-plating (Fig. 1E, white arrowheads). This sectored morphology was most pronounced in CAN^R^ colonies from the Chr4, Chr10, Chr12, Chr13, and Chr15 strains.

### A per-chromosome assessment of the fitness costs associated with monosomy

The frequency of revertants in a monosome-derived population is determined by both the rate of reversion (μ^R^) and the relative growth rates (ω) of monosomic and revertant cells. For example, the high frequency of revertants in Chr4 and Chr12 monosomic populations could reflect a high rate of reversion or the high relative fitness of revertants compared to their slower-growing monosomic predecessors. Conversely, the low frequency of revertants in Chr2 monosomic populations could result from a low rate of reversion or the similar fitness of monosomic and revertant cells. To distinguish between these possibilities on a per-chromosome basis, we measured ω of monosomic and euploid cells using two orthogonal liquid growth assays. In the first assay, light CAN^R^ populations were inoculated in replicate microwell rich media cultures and growth kinetics were monitored over ∼3-4 divisions by optical density (OD). Because revertant subpopulations would distort estimates of ω, this approach was restricted to monosomes that consistently exhibited undetectable revertant frequencies by both WGS and monosome replating tests (Fig. 2A, preselected monosomes). For the second assay, we engineered a new suite of strains that enabled measurement of ω immediately after monosomic cell formation. Hemizygous insertions of the inducible *GAL1* promoter (*GAL1p*) were introduced adjacent to one centromere of each respective homologous pair (*GAL1p-CEN)*(Fig. 2B) (Hill and Bloom 1987; Reid et al. 2008). This cassette also encoded a *URA3* gene, rendering cells prototrophic for uracil (URA+) but sensitive to the drug 5-FOA (FOA^S^) (Boeke et al. 1984). In dextrose-containing media, *GAL1*p is repressed, allowing the centromere to function normally to facilitate accurate spindle-mediated disjunction of the sister chromosome pair in mitosis. Induction of *GAL1p* in galactose-containing media disrupts centromere activity via constitutive transcription, specifically impairing kinetochore assembly and spindle-mediated disjunction of the *URA3-GAL1p*-*CEN* chromatid pair (Tanaka et al. 2007). Consequently, the mother cell retains both copies, becoming a trisomic URA+/FOA^S^ cell and producing a monosomic URA-/FOA^R^ bud cell; the latter cell type can be enriched by selection in 5-FOA containing-media (Fig. 2B). To eliminate pre-existing FOA^R^ cells that could confound fitness measurements, populations were pregrown in media lacking uracil prior to induction with galactose. Induced cultures were then transferred into microwell cultures in dextrose- and 5-FOA-containing media to counterselect against the trisomic population and any residual uninduced disomic cells. The growth kinetics of the FOA^R^ subpopulation generated was monitored by OD. For both assays, OD measurements were used to determine the average maximum slope of the logarithmic growth phase for each monosomic and a euploid control, and ω was defined as the quotient of these values, respectively.

**Figure 2.**
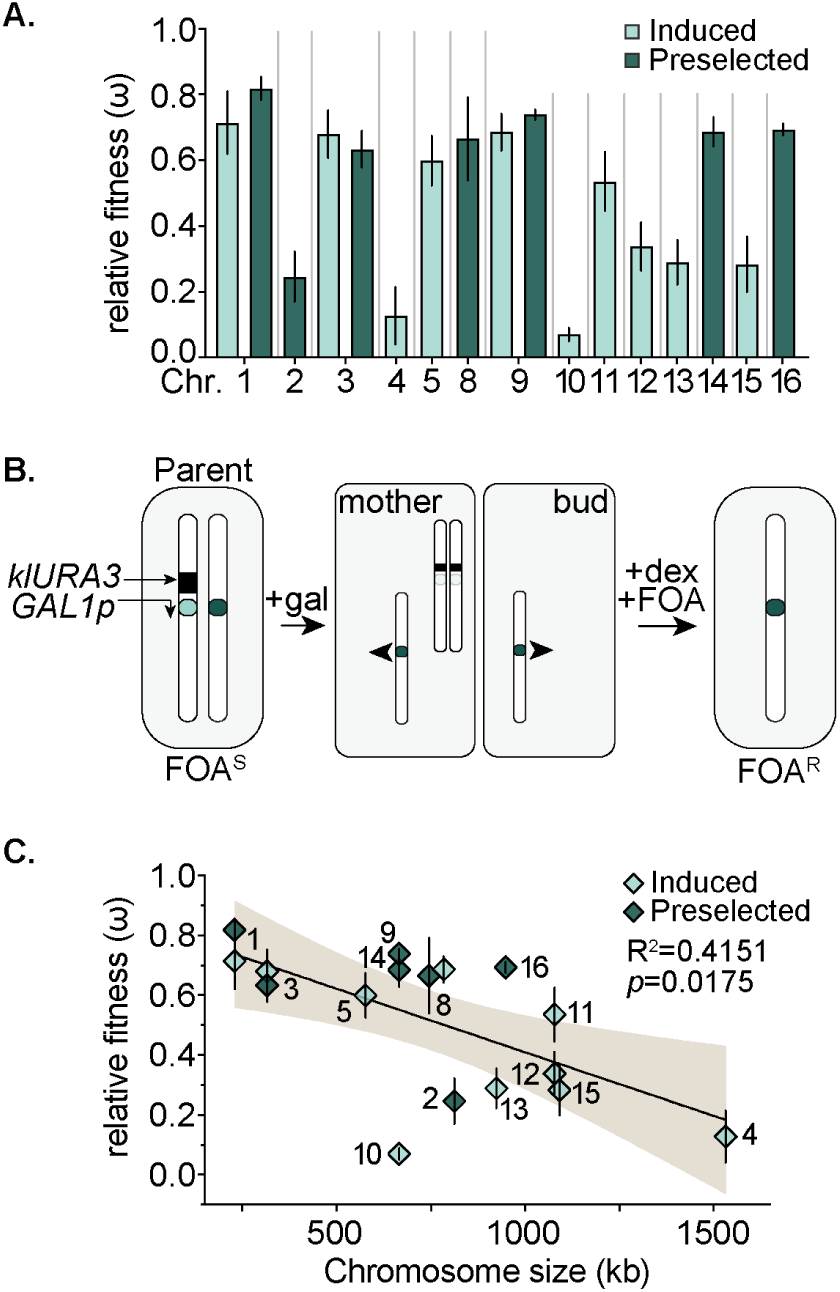
Fitness costs associated with monosomy vary markedly across chromosomes. **A.** The ω values derived from the induced (light green) and preselected (dark green) growth assays for each chromosome **B.** A schematic describing the *URA3-GAL1p-CEN* monosome generation and enrichment approach. **C.** The deduced ω values measured from the induced (light green) and preselected (dark green) growth assays for each chromosome plotted according to chromosome size. Brown curve depicts the 95% confidence intervals produced from the simple linear regression of ω vs. chromosome size.

In both assays, monosomy universally reduced relative fitness, though the costs incurred depended greatly on the chromosome affected (Fig. 2A). These findings provide additional support for the widely accepted model that aneuploidies disrupt essential biosynthetic and metabolic pathways through gene dosage imbalances (Rojas et al. 2024; Hose et al. 2020; Torres et al. 2007; Selmecki et al. 2009; Dutcher et al. 2024; Oromendia et al. 2012; Pompei and Cosentino Lagomarsino 2023; Pavelka et al. 2010). Consistent with the work of others, we identified a statistically significant correlation between ω and chromosome size (Fig. 2C)(R^2^=0.42, p=0.0175). We did not investigate the biological basis of this relationship in the present study, since this has been reported elsewhere (Rojas et al. 2024). These results provide insight into the reversion patterns observed in our earlier experiments by establishing two central principles: 1) fitness influences aneuploid frequency, and 2) reversion probability differs across chromosomes. For example, monosomy of Chr2 confers a severe fitness defect (ω=0.246) yet replated monosomic clones yielded almost no revertants (Fig. 1D), indicating that μ^R^ is very low. In contrast, monosomes of Chr4, Chr12, and Chr13—despite displaying similarly severe fitness costs—produced high frequencies of revertants, indicating that they revert at markedly higher rates.

### Establishing an integrated framework of aneuploid karyotype population dynamics

We proceeded to estimate per-chromosome rates of monosome formation (μ^A^) and reversion (μ^R^) using fluctuation assays (Luria and Delbrück 1943; Lang 2018). For each *CAN1*-*VIO* strain, replicate populations were seeded into microwells containing rich media and expanded to saturation. A fraction of each replicate was plated on canavanine-containing selective media to quantify CAN^R^ colonies, and a diluted fraction was plated on non-selective media to estimate total population size. Dark CAN^R^ revertants and light CAN^R^ monosomes were scored after 2 and 4 days, respectively. To estimate mutation rates from the extant colony counts while also accounting for relative fitness differences between monosomic and euploid cells (both parental or revertant), we used the maximum likelihood estimator (MLE) program mlemur (Łazowski 2023).

Light CAN^R^ colony counts were used to estimate μ^A^ (Fig 3A). The resulting ω-adjusted µ^A^ values varied >10-fold across chromosomes—ranging from 1.2010^-6^ cell^-1^ gen^-1^ (Chr7) to 1.3310^-5^ cell^-1^ gen^-1^ (Chr12)(Fig. 3A)—suggesting that yeast chromosomes are differentially subjected to processes which influence their individual rates of nondisjunction. Similar values were also obtained from orthologous fluctuation tests using populations grown on solid media (Tables S7 and S8). Although chromosome size has been correlated with nondisjunction frequency in humans (Worrall et al. 2018), no significant relationships between μ^A^ and chromosome size, genic-, or non-genic element content were identified (Fig. S2)(Rojas et al. 2024).

**Figure 3.**
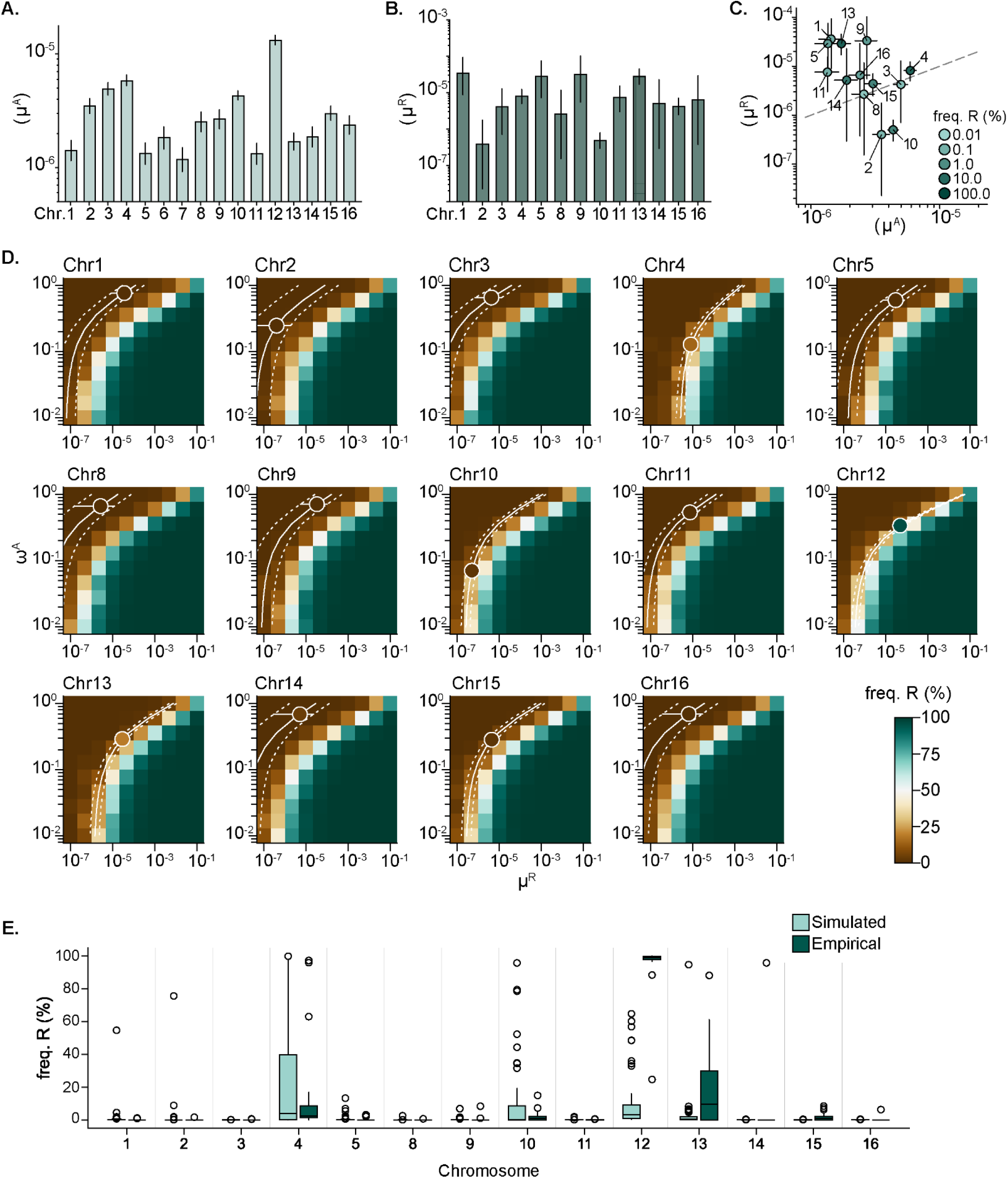
Rates of aneuploidy and reversion generally conform to an unequal likelihood model. **A.** Per-chromosome rates of monosomy (µ^A^) estimated from light CAN^R^ colony counts. **B.** Per-chromosome rates of reversion (µ^R^) estimated from the monosome replating test revertant count data. **C.** Comparison between µ^A^ and µ^R^. Horizontal and vertical bars indicate 95% confidence intervals of rate estimates. Dot color represents the average extant frequency of revertant cells in the replating tests from which µ^R^ values were computed. Dotted diagonal represents identity. **D.** Heatmaps showing the average extant frequency of revertant cells in 50 replicate simulated population expansions for each combination of input parameter values (μ^R^ and ω^A^). For each chromosome, simulated final population sizes were set to the average from the empirical replating tests. Outlined circles represent the empirical values for μ^R^ and ω^A^, and the circle fill color represents the empirically observed mean frequency of revertant cells. Horizontal lines denote the 95% confidence interval of the empirical μ^R^. Solid and dotted curves denote the estimated 95% confidence interval for μ^R^, respectively, fitted on empirical colony counts across the variable range of ω shown on the Y axis, representing the value computed for μ^R^ if there were an experimental error in the estimate of ω^A^. **E.** Comparison of the distributions of revertant frequencies across replicates of the simulated and empirical monosome replating tests. Individual circles denote outliers. Data are also presented as cumulative distribution curves in Fig. S5A.

Estimating μ^R^ required additional consideration. Since a cell must first become monosomic before reverting to form a dark CAN^R^ colony, the rate estimated from dark CANᴿ colony counts would typically be interpreted as the compound rate (μ^C^) of these two sequential mutations:

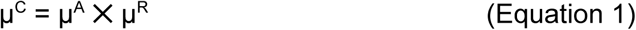

Additionally, because the populations subjected to the canavanine fluctuation tests were founded by diploid cells, dark CANᴿ revertant colonies could arise from two parallel mechanisms: reversion from monosomy or from trisomy (Fig. S3). Monosomic and trisomic karyotypic states form as reciprocal products of the same nondisjunction event; therefore, the rate of trisome formation is equivalent to μ^A^. However, neither the relative fitness of trisomes (ω^T^) nor their rates of reversion (μ^T^)—both of which would influence the frequency of extant dark CAN^R^ colonies—could be directly measured with this selectable-genetic system, precluding accurate estimation of μ^R^. To circumvent these issues, we instead estimated μ^R^ using the revertant colony counts from the monosome replating tests. Because the populations subjected to the replating tests started with monosomic founders, the dynamics of reversion could be directly assessed without having to account for the parallel dynamics of trisomy. The resultant μ^R^ values varied 89-fold between chromosomes (4.0210^-7^(Chr2) to 3.5810^-5^ (Chr1)) (Fig. 3B). For most chromosomes, reversion probability best fits an unequal likelihood model (μ^A^ ≠ μ^R^): Chr2 and Chr10 had lower μ^R^ than μ^A^ (8.75-fold and 8.63-fold respectively), while Chr1, Chr9, Chr11, Chr13, Chr14, and Chr16 had higher μ^R^ than μ^A^ (2.74-fold to 25.03-fold). Only Chr3, Chr4, Chr8, and Chr15 show similar μ^A^ and μ^R^ values more indicative of an equal likelihood model of reversion (Fig. 3C). In general, the consistently low values of μ^R^ indicate that chromosome loss does not substantially perturb genome stability in this organism, clarifying a central facet of aneuploid population dynamics: across the physiologically relevant range of μ^A^, μ^R^, and ω values measured in these experiments, reversion is a minor determinant of monosomic karyotype stability in populations.

Estimation of μ^R^ for Chr12 was confounded by the near-complete sweep of revertants in all replicate monosomic populations. Revertant fixation violates an integral requirement of fluctuation assay-derived rate estimation: mutants must be rare enough that independent mutant lineages remain distinguishable within each population. This high revertant frequency permits at least two alternative interpretations: reversion from monosomy could occur through a single mechanism operating at an exceptionally high rate, or reflect the combined output of multiple parallel mechanisms. To distinguish between these possibilities, we developed fflucsim, a forward simulator of population mutational dynamics. fflucsim performs *in silico* fluctuation tests parameterized by user-specified values for all relevant variables (namely μ^A^, μ^R^, μ^T^, ω^A^, ω^T^, and final population size) and explicitly tracks mutation events, mutant frequencies, and population genealogies. We validated that fflucsim accurately recapitulates standard mutational models by simulating population expansion across a broad range of mutation rates and relative fitness values, then benchmarking the known input rates against those estimated from output mutant counts using rSalvador (Zheng 2017)(Fig. S4).

We used fflucsim to investigate whether the frequency of revertants observed in the monosome replating tests reflected one or multiple mechanisms. If revertants had emerged via a single mechanism, then fflucsim models parameterized solely by ω and μ^R^ should be sufficient to reproduce the observed frequencies. If multiple reversion mechanisms had contributed to the emergence of revertants, even models permitting very high μ^R^ values would fail to recapitulate the observed revertant abundance. We simulated replicate monosomic population expansions across the parameter space defined by experimentally derived ω and μ^R^ values for each chromosome tested, then directly compared the simulated and observed revertant frequencies across the full range of input values. For most chromosomes (Chr1, Chr2, Chr3, Chr5, Chr8, Chr9, Chr10, Chr11, Chr14, Chr15, and Chr16), simulations parameterized with their empirically-derived μ^R^ and ω values produced revertant count distributions that closely matched the observed data; any deviations were fully attributable to minor uncertainties in ω or μ^R^ (Fig. 3D). In contrast, no combination of ω and μ^R^—even μ^R^ values close to 1, which would constitute an essentially deterministic event—was sufficient to recapitulate the high frequency of revertants observed in the replated populations of Chr12 monosomes. In addition, the observed revertant frequency distributions for Chr4, Chr10, and Chr13 also differed from the simulated results (Fig. 3E; Fig. S5), indicating that the reversion dynamics of Chr4, Chr10, Chr12, and Chr13 do not conform to the predictions of standard mutational models. Instead, they better support the interpretation that for these chromosomes, reversion occurs through multiple parallel mutational routes.

Before investigating the molecular basis of these predicted alternative reversion pathways, we re-examined the canavanine fluctuation test results to evaluate the contribution of monosomic cell-derived reversion to the observed frequencies of dark CAN^R^ colonies. If dark CAN^R^ colonies arise solely from the reversion of monosomic and trisomic cells, then the compound rate of formation (μ^C^) would be the sum of rates describing both two-step mutational processes (Fig. S3):

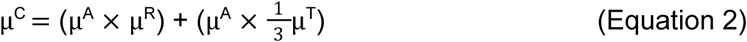

With chromosome-specific μ^A^ and μ^R^ values determined, we inferred the per-chromosome compound rates of monosomic cell-derived reversion (μ^A+R^ = μ^A^ μ^R^) and compared the resulting values to the rates estimated directly from the dark CAN^R^ colony counts (μ^C^) (Fig. 4A). The resulting μ^A+R^ values were ∼2-5 orders of magnitude lower than μ^C^, indicating that reversion from monosomy had contributed minimally to the occurrence of dark CAN^R^ colonies in the tested populations. This result suggested that the majority of observed revertants derived from trisomic cells, reflecting either their increased relative fitness (ω^T^) or higher reversion rates (μ^T^). Higher ω^T^ values would result in greater expansion of trisomic subpopulations, thereby increasing the opportunity for reversion events to occur. Alternatively, if trisomic karyotypes are more genetically unstable, reversion could occur at higher rates than in monosomic cells (μ^T^ ≫ μ^R^). To explore these possibilities quantitatively, we again used fflucsim, this time simulating the expansion of populations founded by diploid ancestors. These simulations incorporated 5 parameters: empirically determined values of ω^A^, µ^A^ and µ^R^, and a broad range of hypothetical values of ω^T^ and μ^T^. We compared the frequency of revertant CAN^R^ cells produced by these simulations to the actual values observed in the CAN^R^ fluctuation tests (Fig. 4B). These simulations show that for most chromosomes, μ^T^ values in the range of 10^-3^-10^-1^ cell^-1^ gen^-1^ would be necessary to produce frequencies of dark CAN^R^ cells compatible with our experimental results (Fig. 4B, white cells). However, the limited experimental data available indicate that trisomic cells are less fit than euploids (ω^T^ < 1) (Pompei and Cosentino Lagomarsino 2023), meaning that μ^T^ values approaching 1 would often be required to reconcile our observations with model outputs. To our knowledge, such values of μ^T^ would be orders of magnitude higher than any aneuploidy rate yet reported and incompatible with previous observations demonstrating the prevalence and long-term stability of trisomic karyotypes in wild populations of yeast (Heasley and Argueso 2022; Sharp et al. 2023; Peter et al. 2018; Zhu et al. 2016). Instead, these results provide additional support for the existence of one or more alternative mechanisms that generate revertant cells. Crucially, both our experimental data and forward simulations indicate that these mechanisms 1) need not proceed through a stable aneuploid intermediate (monosomic or trisomic) and 2) may operate at effective rates or with mutational dynamics which render them poorly described by the constant-rate assumptions underlying standard mutational models.

**Figure 4.**
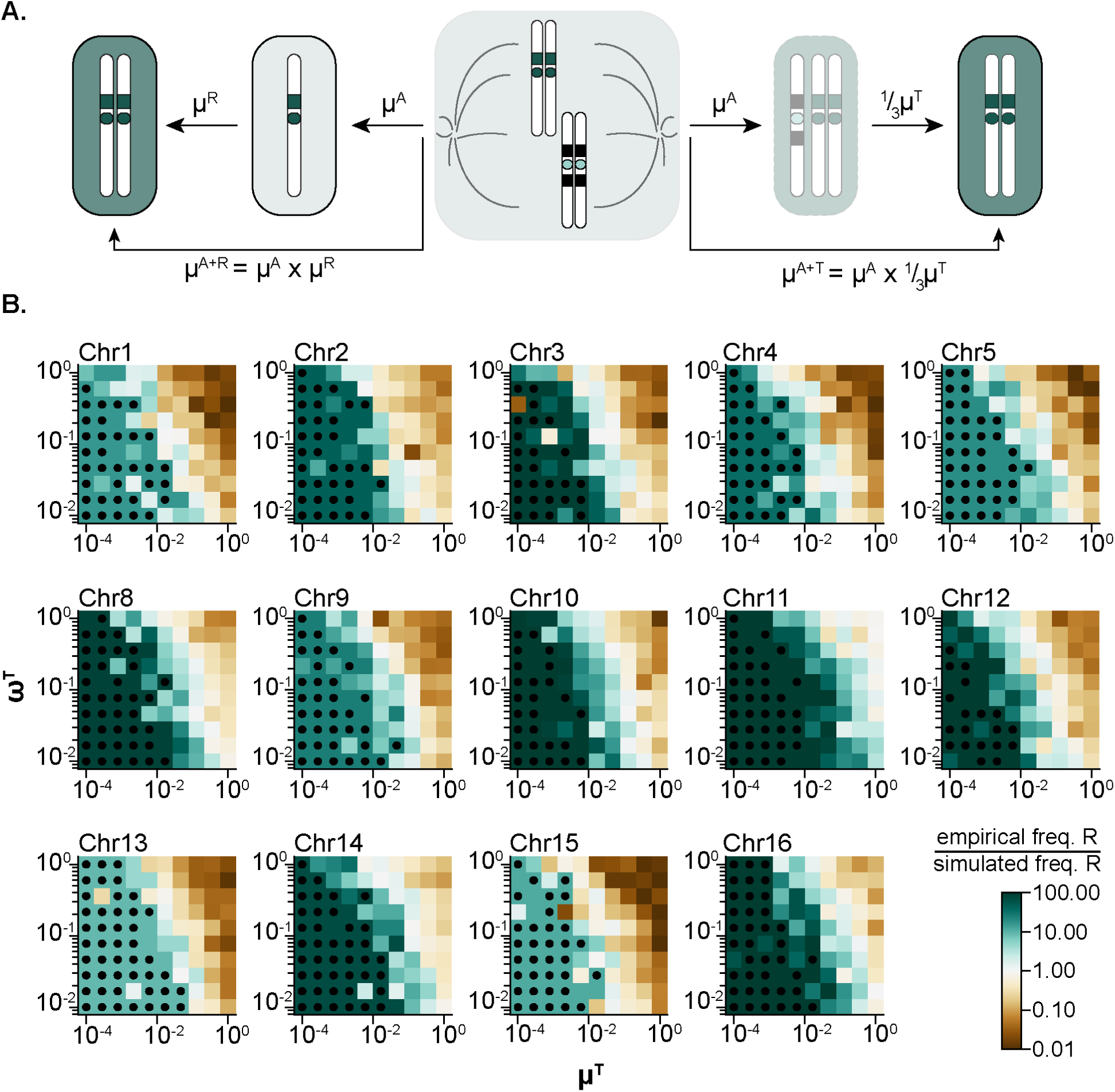
Empirical revertant frequencies are not fully described by reversion from intermediate aneuploid states. **A.** Trisomic cells are produced as reciprocal products of the same aneuploidy event. Monosomic cells can revert to a disomic karyotype at rate μ^R^, while trisomic cells revert to a disomic karyotype at rate μ^T^. **B.** Heatmaps show the fits of the revertant CAN^R^ cell frequencies derived from the experimental and simulated fluctuation assays. Cell color corresponds to the ratio of the mean revertant CAN^R^ cell frequency in 50 replicate simulated populations and the empirical frequency for each combination of input parameter values (μ^T^ and ω^T^); see heatmap legend at bottom left. The empirical estimates were specified for the parameters: μ^A^, μ^R^ and ω^A^. For each chromosome, simulated final population sizes were set to the average from the CAN^R^ fluctuation assays. Dots mark parameter combinations for which no viable revertant cell was produced across all 50 simulated replicates, and were substituted by a lower bound frequency corresponding to a single viable revertant cell in a single replicate.

### Alternative mutational mechanisms facilitate generation of revertant karyotypes

To better understand the alternative reversion mechanisms implicated by our rate analyses, we performed a focused molecular characterization of reversion events guided by observations from the WGS analysis of light and dark CANᴿ clones. Unlike most light CAN^R^ colonies—in which only the copy number of the predicted chromosome was affected—each clone from the Chr6 and Chr7 strains carried 1-13 additional unselected karyotypic alterations, many of which were mosaic within the sequenced population (Fig. 5A, Supplemental Table 2). This mutational signature is typical of chromosomal instability (CIN), a mutator phenotype defined by the constitutive, high frequency accumulation of aneuploidies and other structural variations (Zhu et al. 2012; Funk et al. 2016). Indeed, aneuploidies of Chr6 are known to induce CIN because copy number imbalances of Chr6 resident genes *ACT1* (actin) and *TUB2* (beta-tubulin) disrupt spindle assembly and positioning, increasing nondisjunction rates (Zhu et al. 2012; Anders et al. 2009; Espinet et al. 1995; Deutschbauer et al. 2005; Sheltzer et al. 2011). In contrast, the dark CANᴿ clones from the same cultures had acquired only the predicted UPD, with none of the additional karyotypic alterations harbored by their matched light counterparts (Fig. 5B). Given the high aneuploid burden and ongoing instability present in the light clones, it is improbable that they were the direct predecessors of the dark clones, particularly within the generational limits of our experiments. Rather, these genotypic patterns imply that revertant UPDs can also arise by mechanisms that bypass a stable aneuploid intermediate state. One proposed mechanism, called reciprocal UPD, posits that intermolecular DNA- or protein-based linkages persisting between homologs in mitosis could facilitate their simultaneous nondisjunction, producing daughter cells each carrying a UPD of one homolog (Andersen and Petes 2012). Yet, direct evidence demonstrating that intermolecular linkages are sufficient to drive cooperative missegregation of multiple chromosomes has remained limited.

**Figure 5.**
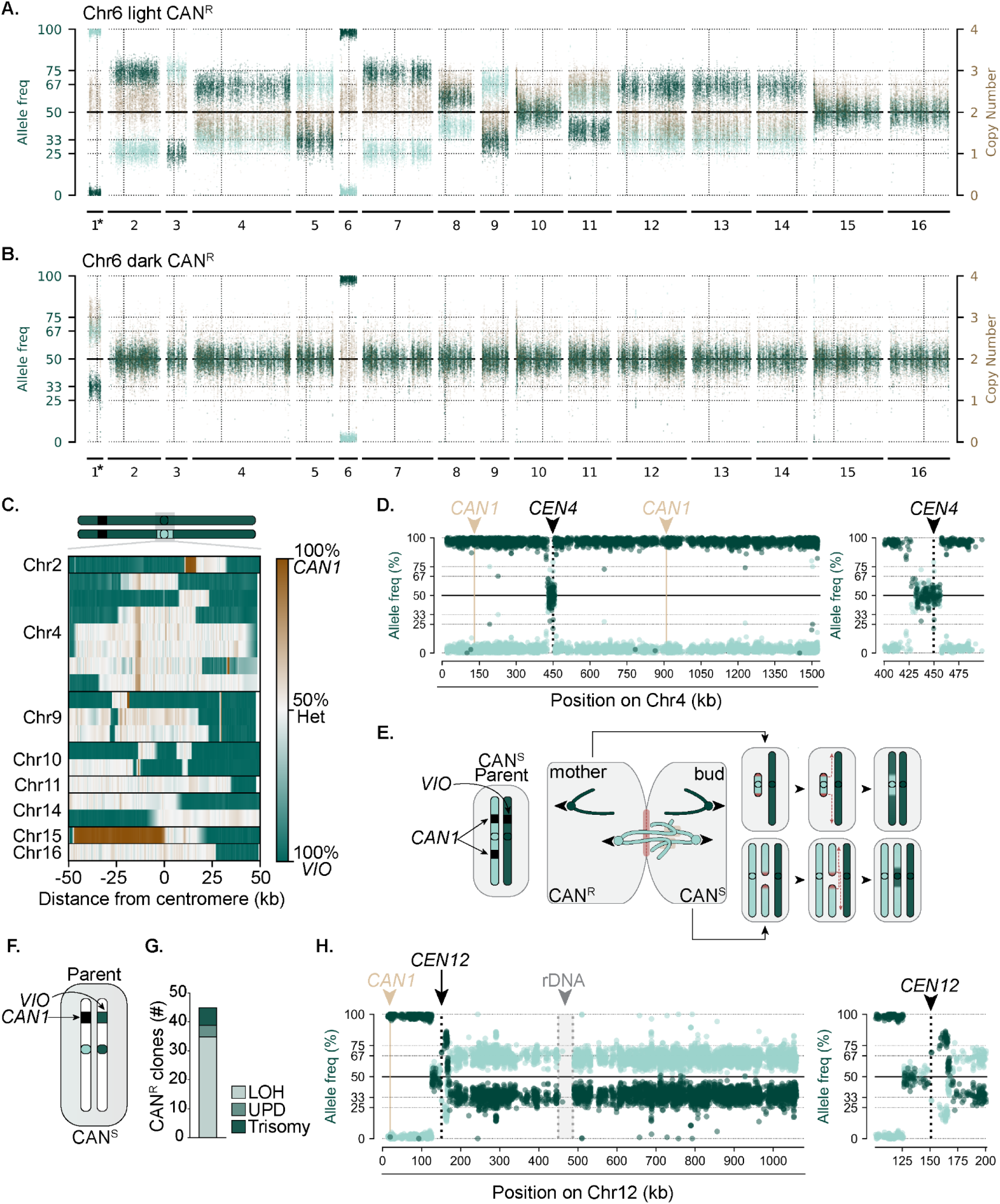
UPD karyotypes develop independent of a stable aneuploid intermediate state. **A. and B.** Representative WGS mappings of light and dark CAN^R^ clones derived from the Chr6 *CAN1*-VIO strain. Each dot denotes a heterozygous allelic position; colors denote normalized depth of coverage (brown), and the allele frequencies of the *CAN1*-marked chromosome (light green) and *VIO*-marked chromosome (dark green). Vertical dashed lines delineate centromeric positions. Trisomy of Chr1, denoted with the asterisk, is a preexisting aneuploidy present in the parental Chr6. **C.** A heatmap of allele frequency at each parental heterozygous position to illustrate the pericentromere-biased breakpoint positions observed in recombinant dark CAN^R^ clones. **D.** A representative Chr4-specific WGS mapping of a dark CAN^R^ colony recovered from the Chr4 *CAN1-*VIO strain displaying the characteristic pericentromeric breakpoints and heterozygosity. Each dot denotes a heterozygous position; color denotes the *CAN1*-marked chromosome (light green) and VIO-marked chromosome (dark green). **E.** A schematic illustrating the predicted mechanism and outcomes of a sister-sister intermolecular linkage mediated missegregation event. **F.** A schematic of the single *CAN1*-VIO cassette Chr12 strain. **G.** Quantification of the karyotypic spectra harbored by CAN^R^ clones derived from the single-cassette strain. **H.** Chr12-specific whole-chromosome or centromere-proximal WGS mappings of a recovered trisomic CAN^R^ colony.

Another karyotypic pattern captured in the WGS analysis provided insight into the plausibility of such a model: while most dark CAN^R^ clones harbored a typical UPD, ∼10% harbored a disomic genotype defined by a short heterozygous tract bounded by pericentromeric breakpoints (Fig. 5C). Dark CAN^R^ clones from the Chr4 strain exhibited a pronounced bias for this genotype (8/10 sequenced clones) (Fig. 5D), a finding that was particularly surprising given that the *CAN1* markers on Chr4 were separated by more than 750kb and positioned at least 300kb from the centromere in either direction. In principle, such molecules could form via independent, sequential recombination events on each arm, but this mechanism would not explain the pericentromeric breakpoint enrichment we observed. Similarly, although centromeric gene conversion—in which breaks in one homolog’s centromere are repaired using sequence from the other (Kozmin et al. 2024)—can generate recombinant molecules, the resulting breakpoint patterns are distinct from those observed in dark CANᴿ clones. Instead, their breakpoint bias more closely resembles the mutational signature generated by the break–fusion–bridge cycles of dicentric chromosomes (Lopez et al. 2015; McClintock 1941; Guérin and Marcand 2022). Previous studies have shown that when dicentric chromosomes are engaged by microtubules emanating from opposite spindle poles in mitosis, they are broken during cytokinesis, typically at positions near one of the centromeres (Lopez et al. 2015; Song et al. 2013; Cook et al. 2024). Although neither WGS nor the visible phenotypes of these recombinant clones indicated the presence of dicentric molecules undergoing iterative cycles of breakage and repair, the similarity in breakpoint distribution suggested the existence of alternative molecular scenarios in which two centromeres would behave as though physically linked.

We reasoned that sister chromatids, homologs, or even nonhomologous chromosomes connected by persistent intermolecular linkages during mitosis could mimic the missegregation dynamics of a dicentric and result in the formation of the recombinant molecules recovered in the CAN^R^ clones (Guérin and Marcand 2022; Lopez et al. 2015; Chan et al. 2007; Finardi et al. 2020; Quevedo et al. 2012; Uhlmann et al. 1999; Holm et al. 1985, 1989). One variation of this model—in which such linkages persist between sister chromatids—is illustrated in Fig. 5E. If intermolecularly linked sisters segregate to opposite spindle poles during mitosis, the persistent linkages between them will form DNA bridges during anaphase. Left unresolved, this bridged DNA will be severed during cytokinesis (Chan et al. 2018, 2007; Lopez et al. 2015; Quevedo et al. 2012). The resulting daughter cells will inherit complementary fragments of the linked pair: one retaining only the centromeric fragment, the other retaining the distal arm fragments together with the intact sister chromatid (Fig. 5E). In the next cell cycle, homology-templated repair of these fragments would generate recombinant chromosomes (Quevedo et al. 2012). Repair of the centromere-containing fragment using the homolog as the template would yield a homologous pair that retains heterozygosity near the centromere but is homozygous along the arms—precisely the signature displayed by the recombinant CAN^R^ clones (Fig. 5C-D). Oppositely, repair of the arm fragments using either the sister chromatid or the homolog as a template would result in trisomy or tetrasomy, with at least one molecule potentially containing sequences derived from both homologs.

Under this model, and constrained by the positioning of the dual *CAN1* cassettes in our system, clones that inherited the centromeric fragment were more likely to be recovered by CAN^R^ selection than the reciprocal trisomic or tetrasomic products, since these would likely retain at least one *CAN1* cassette and fail to form colonies on canavanine-containing media. Indeed, in the >500 CAN^R^ clones sequenced in this study, only a few recombinant trisomes displayed the allelic pattern predicted in Fig. 5C. Thus, to more rigorously test this model, we engineered a new strain in which the homologs of Chr12 harbored either a single *CAN1* or pigment cassette ∼131kb from the centromere and 19kb from the left telomere (Fig. 5F). This positioning increased the likelihood of recovering recombinant trisomic products, although most dark CAN^R^ clones were expected to harbor simple copy-neutral LOH tracts that eliminated the *CAN1* cassette and duplicated the pigment operon. Indeed, 77.7% (35/45) of dark CAN^R^ clones recovered harbored short terminal tracts of LOH near the chromosome end (Fig. 5G). However, the other 22.2% harbored aneuploidies of Chr12—both UPDs and trisomy. Consistent with the predictions of the intermolecular linkage model, every trisomic clone contained at least one recombinant molecule with pericentromeric breakpoints (Fig. 5H, Supplemental Figure 6). In addition, many displayed complex recombination patterns including LOH across all three molecules or intermediate allele frequencies reflecting mosaicism within the sequenced populations. Both suggest that generation of the recombinant trisomy may have involved multiple broken molecules or repair templates, or the production of unstable recombinant products (Vasan et al. 2014; Tsaponina and Haber 2014).

The recombinant patterns exhibited by the disomic and trisomic CANᴿ clones are consistent with a model in which intermolecular linkages between molecules can coordinate their missegregation, leading to breakage and subsequent repair by homologous recombination. Yet, these results did not formally exclude alternative mechanisms such as sequential recombination events or centromeric gene conversion. Nor could these CAN^R^ systems directly test a central assumption of the model: that bipolar segregation of linked molecules in mitosis is necessary for the generation of the recombinant products. We used the *URA3-GAL1p-CEN* strains to test this assumption. In yeast, forces generated by the elongating anaphase spindle are required to drive chromosome translocation through the bud neck constriction into the bud cell (Yeh et al. 2000; Li et al. 1993; Palmer et al. 1992). Conditional centromere inactivation—via *GAL1p* induction—prevents kintochore-mediated coupling of these segregation forces to the specific pair of sister molecules harboring the *URA3-GAL1p-CEN* cassette, resulting in their retention in the mother cell. In this controlled context, only monosomic bud cells arising from divisions in which the *URA3-GAL1p-CEN* sister pair remained unsegregated in the mother cell would be predicted to manifest a FOA^R^ phenotype (Fig. 2A). Alternatively, recovery of FOA^R^ cells harboring a recombinant product of the inactivated chromosome would indicate that its formation occurs independently of direct spindle-mediated segregation, but coincident with bipolar segregation of the remaining chromosomes. We assessed these possibilities by plating galactose induced populations of *URA3-GAL1p-CEN* strains to solid media containing 5-FOA, allowing FOA^R^ colonies to develop, and isolating individual clones for characterization by WGS (Fig. 6A).

**Figure 6.**
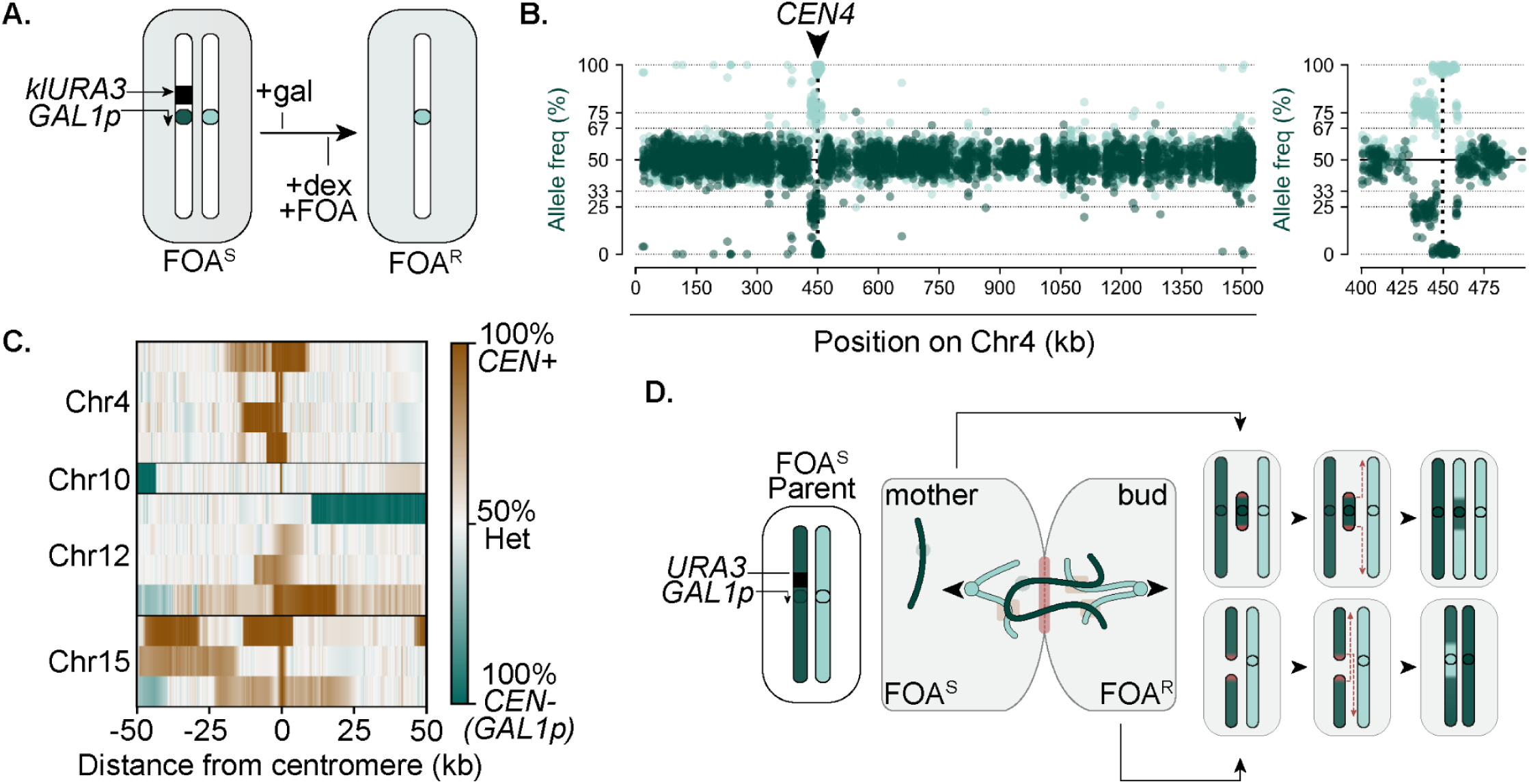
Recombinant chromosome generation coincides with chromosome segregation. **A.** Schematic of the centromere inactivation assay. **B.** A representative Chr4-specific WGS mapping of a FOA^R^ colony recovered from the *URA3-GAL1p-CEN4* strain that displays pericentromeric breakpoints. **C.** A heatmap of allele frequency at each parental heterozygous position illustrating the pericentromere-biased breakpoint positions observed in FOA^R^ clones. **D.** A schematic illustrating the predicted mechanism and outcomes of an intermolecular linkage event that generates an FOA^R^ daughter cell bearing the recombinant genotypes shown in B. and C.

Consistent with our previous results (Fig. 1D), most sequenced FOA^R^ clones exhibited stable monosomic or revertant mosaic signatures. Yet, 19% displayed a recombinant disomic signature typified by centromere-proximal breakpoints, with Chr4 again overrepresented among this group (4/10 sequenced clones)(Fig. 6B, C). This recombinant genotype resembled the one displayed by the CANᴿ clones, with one key difference: whereas CAN^R^ recombinant chromosome pairs retained heterozygosity only near the centromere, FOA^R^ recombinant pairs were homozygous in the pericentromeric region but retained heterozygosity along the length of the chromosome arms (Fig. 6B, C).

Such products could only arise if the arm segments of an inactivated chromatid were segregated into the bud cell—a process that, to our knowledge, cannot occur in the absence of spindle-generated force. Rather, this result implies that bipolar spindle-generated forces were transmitted to the inactivated chromatid by an indirect mechanism, the most parsimonious being via linkage to actively segregating chromosomes (Fig. 6D). As illustrated in Fig. 6D, a possible outcome of this type of event would be the breakage and partial inheritance of the inactivated chromatid. The daughter cell inheriting the arm fragments would become FOA^R^ if those fragments were repaired using the homolog as a template, generating a pair of chromosomes precisely displaying the recombinant signature observed in the FOA^R^ clones. This result refines a central principle of the intermolecular linkage model: while mitotic chromosome segregation is necessary for recombinant molecule formation, the direct segregation of linked partners is not. Intermolecular linkages can sufficiently transfer the forces necessary to establish a chromosomal architecture that will result in breakage and asymmetric inheritance of the linked molecules.

22% of sequenced FOA^R^ Chr12 clones were typified not by pericentromeric breakpoints, but by a breakpoint within the ribosomal DNA (rDNA) array on the right arm instead (Fig. S7). Due to the homotypic and repetitive architecture of the rDNA array, we were unable to define the precise location of these breakpoints. Like the Chr12 trisomic clones (Fig. 5F-G), these clones usually displayed mosaic coverage and allele frequency, indicating the existence of multiple distinct molecules of Chr12. Such patterns are consistent with a model in which intermolecular linkages within the rDNA lead to chromosome breakage during mitosis (Quevedo et al. 2012), and that repair of these fragments may occur through a process yielding diverse molecular outcomes. Notably, in addition to the selected chromosomal aneuploidy, 19% of FOA^R^ clones derived from the other *URA3-GAL1p-CEN* strains also harbored terminal tracts of LOH originating in the rDNA and extending to the right telomere. The coincidence of this independent mutation suggests that the rDNA may be a hotspot for linkage formation (Fig. S7).

Although the *URA3-GAL1p-CEN* system generates a mitotic scenario that would rarely occur in wild-type cells—because an unattached chromatid pair typically activates a mitotic checkpoint-mediated cell cycle arrest(Musacchio and Salmon 2007; Musacchio 2015)—it constituted a valuable tool to further probe the validity of the intermolecular linkage model. Next, we used time-lapse fluorescence microscopy to visualize the segregation dynamics of *CEN4* throughout mitosis in dextrose- and galactose-containing media (Tanaka et al. 2005; Pearson et al. 2001). Into the *URA3-GAL1p-CEN4* strain, a multicopy *tetO* array was introduced adjacent to the *GAL1p-CEN4* cassette, along with fluorescently-tagged alleles of the *tetO*-binding protein (TetR-GFP) and alpha tubulin (mRuby-Tub1). Proliferating populations were imaged over time and the *CEN4* disjunction dynamics in mitosis were assessed in each condition. As expected, 100% of observed *CEN4* disjunction events (59/59) occurred normally in dextrose-containing media: the duplicated *tetO* loci localized between the spindle poles in prometaphase and, at anaphase onset, remained closely associated with the spindle poles as they separated into the mother and bud cells (Fig. 7A, Supplemental Movie 7A). In galactose-containing media, we expected to see two disjunction phenotypes: 1) normal disjunction in cells that had not sufficiently inactivated *CEN4 (Palme et al. 2021)*, and 2) inactivation-mediated nondisjunction, in which both *tetO* loci dissociated from the spindle throughout mitosis and remained unsegregated within the mother cell (Fig. 7B, Supplemental Movie 7B). Indeed, we observed these behaviors in 28.3% (17/60) and 55% (33/60) of dividing cells, respectively. However, we also observed alternative segregation events affecting *tetO* pairs that were clearly dissociated from the spindle at the time of anaphase onset (Fig. 7C-D, Supplemental Movies 7C-D). In 6.6% of dividing cells, both *tetO* loci segregated into the bud cell (Fig. 7C, Alternative Type 1). Unlike normal disjunction—in which the centromeres remain closely juxtaposed to the poles as the spindle elongates—the *tetO* loci in these cells showed no directed poleward movement at anaphase onset. Only late in anaphase, when the spindle reached its maximum length, did the two *tetO* loci abruptly translocated into the bud cell (Fig. 7C).

**Figure 7.**
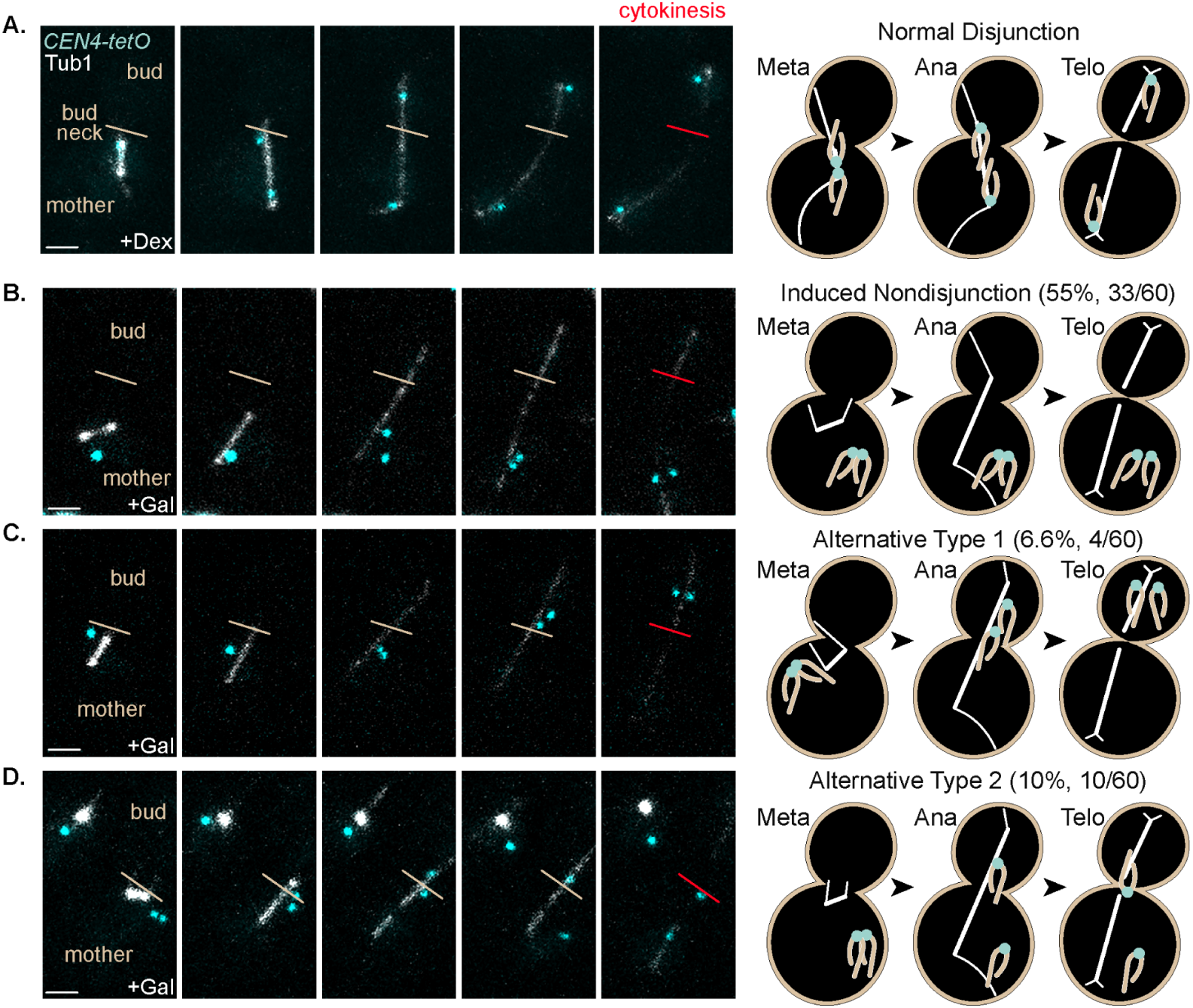
Chromosome missegregation occurs independently of direct spindle attachments. A.-D. Representative time-lapse image series (left) and interpretive schematic and quantitation (right) of *URA3-GAL1p-CEN4-tetO* cells expressing TetR-GFP and Tub1-mRuby2: **A.** a cell in dextrose-containing media progressing normally through mitosis, **B.-D.** cells in galactose-containing media displaying (**B.**) the expected nondisjunction of the inactivated chromosome pair encoding, (**C.**) Alternate Type 1 missegregation in which both inactivated chromosomes segregate into the bud cell (**D.**), or Alternate Type 2 missegregation in which the inactivated molecules migrate into the budneck region prior to cytokinesis. The red budneck delineation denotes the completion of mother-bud cell separation by cytokinesis as assessed by brightfield imaging. Scale bars, 2μM.

In approximately 10% of cells, one or both *tetO* loci migrated to the bud neck region before moving either into the bud cell or back into the mother cell (Fig. 7D, Alternative Type 2). Two distinct behaviors were observed at this stage. In some cases, bud neck localization was transient and culminated in rapid movement of a *tetO* locus into either compartment at the time of cytokinesis, resembling the dynamics of dicentric chromosome severing (Lopez et al. 2015). In other cases, *tetO* loci remained at the bud neck for extended periods and drifted slowly back into the cell interior only after cytokinesis had occurred (Fig. 7D). *TetO* pairs displaying these alternative segregation patterns lacked any detectable colocalized Nuf2, an outer kinetochore protein required for microtubule binding (Fig. S8), again demonstrating that their bud-cell directed movements occurred independently of direct spindle attachment but concurrent with chromosome segregation.

To determine whether intermolecular linkages with actively segregating chromosomes facilitated these alternative *tetO* segregation behaviors, we repeated this experiment using a strain expressing a fluorescently tagged histone (Htb2-TdTomato) and analyzed the chromatin dynamics in Alternative Type 2 cells. Consistent with the intermolecular linkage model, 100% displayed chromatinized DNA bridges spanning the dividing nuclei and encompassing at least one *tetO* locus for variable durations—this was never observed in cells exhibiting normal segregation dynamics (Fig. 8A). Transient, rapid movement of *tetO* loci into the bud neck coincided with bridge resolution (Fig. 8B, Supplemental Movie 8B), indicating that these bridges consisted of chromatin linking at least one unattached *tetO*-bearing chromatid to a chromosome that had segregated into the bud cell. As is shown in Fig. 8B, only after the abrupt, late-stage translocation of one *tetO* locus through the bud neck does the chromatin bridge which had spanned the dividing nuclei for the duration of anaphase finally resolve (Fig. 8B, black arrowhead, 18’). Prolonged localization of *tetO* loci at the bud neck reflected a distinct chromatin architecture and mode of bridge resolution, which occurred only at cytokinesis through the apparent severing of chromatin proximal to the bud neck. In 38% of cases, this process generated a chromatin fragment containing at least one tetO locus; this fragment was never reincorporated into the primary nuclear compartment of the inheriting daughter cell but instead remained cytoplasmically separated throughout the subsequent cell cycle. In the example shown in Fig. 8C, both *tetO* loci are encompassed within the chromatin bridge, although their relative positions with respect to the bud neck differ. In this instance, the orientation of the *tetO* loci and the Htb2 signal suggests that the unattached *tetO-*bearing chromatids may be linked both to one another and to chromosomes that have segregated into each cell. Following cytokinesis, the *tetO* locus positioned closest to the bud neck resolves into a new, chromatinized compartment in the mother cell, leaving the bud cell devoid of this *CEN4*-adjacent DNA fragment (Fig. 8C, Supplemental Movie 8C). Together, these observations demonstrate that intermolecular linkages promote centromere-proximal breakage and loss of centromere-bearing DNA, leading to daughter cells with incomplete or damaged chromosomes. These findings directly support our proposed mechanism of recombinant CAN^R^ and FOA^R^ clone formation (Fig. 8D).

**Figure 8.**
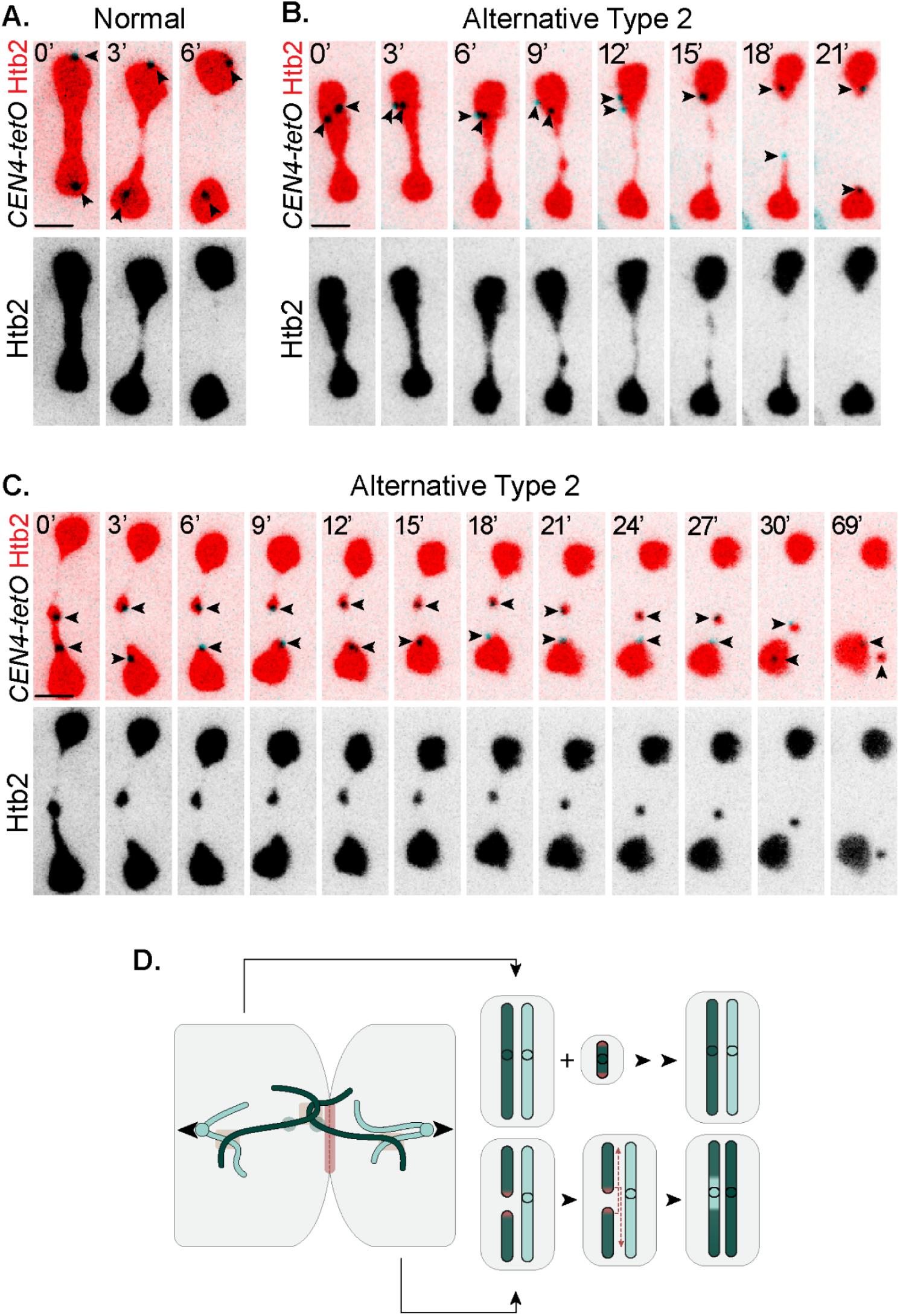
Chromatin bridges and chromosome fragmentation coincide with alternative missegregation events. **A.-C.** Representative time-lapse image series of *URA3-GAL1p-CEN4-tetO* cells expressing TetR-GFP and Htb2-tdTomato that exhibit **A.** normal segregation of the *CEN4-tetO* locus and resolution of the dividing nuclei, **B.**, alternative *CEN4-tetO* segregation and persistence of a long-lived chromatin bridge that resolves only after the rapid translocation of one *tetO* locus into the bud cell, and **C.**, alternative *CEN4-tetO* segregation and persistence of a chromatin bridge that resolves to form a *tetO*-containing chromosome fragment. Time-stamps (in minutes) are relative to the first frame of the displayed series. Arrowheads denote the two *tetO* loci. **D.** A schematic illustrating the observed chromatin dynamics and predicted molecular outcomes of the alternative missegregation even displayed in C. Scale bars, 2μM

## Discussion

The inherent reversibility of aneuploid karyotypes has long been recognized, yet the impacts on population-level dynamics have remained difficult to discern. Here we empirically quantified the occurrence, associated fitness effects, and reversibility of monosomic karyotypes. We developed new *in silico* mutational models with which to integrate these data, yielding a unified framework of the determinants of aneuploid karyotype diversity and stability in populations. Because we evaluated monosomies of multiple different chromosomes, this framework encompasses the physiologically relevant range of rates and fitness coefficients associated with aneuploidy in *S. cerevisiae*, enabling us to articulate general principles of aneuploidy dynamics and to resolve additional mutational mechanisms that impact the basal stability and inheritance of chromosomes.

A primary goal of this study was to define how reversion constrains aneuploid karyotype stability. We find that the dynamics of canonical 2-step reversion are generally explained by an unequal likelihood mutational model: a cell loses a chromosome and then, via a subsequent random nondisjunction event, reverts to a disomic state. But this mechanism occurs rarely (<10^-4^ per cell per generation), indicating that 2-step reversion contributes minimally to the stability of aneuploid subpopulations under conditions where populations expand from euploid precursors in permissive, nearly static environments. In such scenarios, the relative fitness of aneuploid cells is the primary determinant of karyotype frequencies, suggesting that selection, rather than reversion, dominates the maintenance and diversity of aneuploid lineages. The fitness costs and mutation rates derived in the growth conditions used in this study are likely to differ across stressful or fluctuating environments. Consequently, and in line with a growing body of work, the extent to which natural selection shapes karyotypic diversity within populations is strongly dependent on the environment (Merlo et al. 2006). However, reversion may become a more influential determinant of karyotype stability in populations that are already highly karyotypically diverse (Bakker et al. 2016; Gao et al. 2016; Casasent et al. 2018; Bollen et al. 2021; Starostik et al. 2020), or in those undergoing major shifts in effective size or structure—such as bottlenecks, founder events, or selective sweeps. Indeed, reversion had a markedly greater impact on the karyotypic diversification of populations derived from aneuploid founders than from euploids. Moving forward, the influence of specific environmental or population characteristics, and how they modulate the primary determinants of aneuploid karyotype stability, can be systematically interrogated using the experimental systems and computational models presented in this work.

A second objective of this study was to investigate aneuploidy dynamics across a broad chromosomal context. We find substantial variation in nondisjunction incidence across the different yeast chromosomes. Discerning the underlying basis for this variance warrants further study, as doing so is certain to clarify central principles of genome stability and refine current paradigms of aneuploidy-driven disease risk. Perhaps variations in centromeric sequences, which are highly variant and fast-evolving across Eukarya (Altemose et al. 2022; Salinas-Luypaert et al. 2025; Henikoff and Henikoff 2012; Krassovsky et al. 2012; Bensasson 2011; Kozmin et al. 2024; Helsen et al. 2025), may underlie chromosome-specific differences in kinetochore assembly and disjunction fidelity. Alternatively, the spatial organization of chromosomes within the nucleus may impact the likelihood of nondisjunction (Duan et al. 2010; Costantino et al. 2020; Kim et al. 2017). Our work presents an additional possibility: for certain chromosomes—particularly those exhibiting the highest rates of loss—nondisjunction also frequently occurs through a mechanism defined by intermolecular linkage-mediated missegregation, chromosome breakage, and homologous repair. We hypothesize that the propensity to form such linkages—a result of chromosome size, the presence of hyper-recombinant loci, the enrichment of cohesin and condensin complexes, etc.—is an important, yet understudied contributor to the observed diversity in nondisjunction rate.

The genetic systems used in this study allowed detection of but a subset of outcomes predicted by our models of linkage-mediated missegregation, suggesting that we have substantially underestimated the true incidence of this mutational mechanism in populations. For example, recombinant products formed when linked molecules are severed in repetitive regions—such as the rDNA (Quevedo et al. 2012), subtelomeric arrays, telomeric repeats, or dispersed repeats—may not resolve with detectable breakpoints or heterozygous tracts displayed by the recombinant clones characterized in this study. Instead, these events would manifest as typical UPDs, making them indiscernible from the revertant karyotype formed through the canonical 2-step reversion pathway. This possibility is particularly salient when interpreting the reversion dynamics of Chr12, which displayed an exceptionally high reversion rate and penetrant revertant WGS signature consistent with a typical UPD. Although the incidence of recombinant CAN^R^ or FOA^R^ clones was low, many such clones displayed signatures of rDNA breakage and repair. Genetic and chemical perturbations affecting rDNA transcriptional regulation, condensation, and decatenation have been shown to promote the formation of intermolecular rDNA linkages and chromosome severing in cytokinesis (D’Amours et al. 2004; Quevedo et al. 2012; Machín et al. 2006). Our work prompts the hypothesis that even in unperturbed wild type cells, persistent linkages existing within the ∼1Mb rDNA array facilitate high-frequency chromosome nondisjunction events capable of resolving into a simple UPD karyotype through a complex mechanism of templated repair. Our future studies will seek to define the positional and architectural features of such molecular linkages within the rDNA and genome-wide, to determine why some chromosomes are subject to this mode of genomic instability more than others, and to discern the molecular mechanisms by which repair of severed chromosomes occurs subsequent to nondisjunction.

## Data availability

Sequencing reads are available at NCBI under accession number PRJNA1414607. Custom scripts for data analysis and visualization are available at GitHub (https://github.com/heasleyl/A_unified_model_of_aneuploid_karyotype_dynamics). The source code of fflucsim and scripts for running the simulations presented in this study available at GitHub (https://github.com/mhenault1/fflucsim).

## Acknowledgements

We thank those who provided generous support and feedback for this work: J. Moore, R. Rothstein, J. Boeke and J. Temple for sharing strains, plasmid constructs, and equipment; V. Fogg for useful feedback on this work throughout its development; H. Vincent and J. Lucas Argueso for their input during the early design and testing of the genetic systems used in this study (partly supported through grant R35GM119788 to JLA); A. Nguyen, C.G. Pearson, and J. Lucas Arugeso for their feedback on this manuscript; Finally we are grateful for funds from the National Institutes of Health (GM134193; LRH), the Natural Sciences and Engineering Research Council of Canada (Postdoctoral Fellowship, MH) and the Fonds de Recherche du Québec – Santé (Postdoctoral Fellowship, MH).

## Author contributions

Conceptualization: LRH, MH, and LMW; Methodology: LRH, MH, and LMW; Investigation: LRH, MH, and LMW; Formal Analysis: LRH, MH, and LMW; Data Curation: LRH, MH, and LMW; Visualization: LRH and MH; Software: MH and LRH; Validation: LRH and MH; Writing: LRH, MH, and LMW; Supervision: LRH; Project Administration: LRH; Funding Acquisition: LRH and MH.

## Methods

### Strains and Media

Strains and plasmids used in this study are listed in Supplementary Table 1. Strain construction was performed using standard transformation, crossing, and sporulation procedures. Directed insertions of selectable cassettes and reporter constructs were validated using PCR and WGS. Depending on the experiment, yeast cells were grown in one of the following media: rich nonselective media (10g/L yeast extract, 20 g/L Peptone, 20 g/L bacteriological agar, and 20g/L glucose (2.0%)), synthetic-defined complete media (20g/L glucose or 20g/L galactose, 5g/L ammonium sulfate, 1.7g/L yeast nitrogen base without amino acids, 1.4g/L complete drop-out mix, 20g/L bacteriological agar), canavanine-containing media (20g/L glucose, 5g/L ammonium sulfate, 1.7g/L yeast nitrogen base without amino acids, 1.4g/L arginine dropout mix, 20g/L bacteriological agar, 0.06g/L canavanine sulfate), uracil drop-out media (20g/L glucose or 20g/L raffinose, 5g/L ammonium sulfate, 1.7g/L yeast nitrogen base without amino acids, 1.4g/L uracil dropout mix), 5-FOA containing media (20g/L glucose, 5g/L ammonium sulfate, 1.7g/L yeast nitrogen base without amino acids, 1.4g/L complete drop-out mix, 20g/L bacteriological agar, g/L 5-Fluoroorotic Acid).

### Canavanine Fluctuation Tests

*Liquid media pregrowth:* For each *CAN1*-marked strain, replicate ∼500-cell populations were seeded into microwells containing YPD and expanded for 48hrs (median population size = 7.3×10^6^, median number of generations = 13.8). *Solid media pregrowth:* Each *CAN1*-marked strain was streaked onto YPD plates to single cell density and independent colonies were allowed to develop for 48hrs before isolating each and resuspending in 200uL of sterile water. *Plating:* A fraction (20-40%; average 3-7×10^6^ cells) of each replicate was plated on canavanine-containing media, and a diluted fraction plated on YPD. Plates were incubated at 30°C for 96 hrs; both dark CAN^R^ and YPD-plated colonies were counted after 48hrs, the light CAN^R^ colonies at 96hrs. *Data availability:* Raw colony counts, frequency calculations, and inferred population sizes are presented in Supplemental Table S7. For strains LRH587, LRH585, Chr684, and LRH588, we were unable to successfully integrate the complete VIO cassette, even after multiple attempts. Therefore, revertant phenotypes were deduced on the basis of colony size at 48hrs. WGS demonstrated that revertant genotype identification by colony size was accurate.

### Monosome replating tests

#### Plating

Small (48-72hr) light CAN^R^ colonies isolated from independent canavanine plates were resuspended and diluted to single-cell density on synthetic defined complete media plates and incubated at 30°C for 48hrs. After 48hrs, dark revertant colonies were quantified; after 96hrs, light monosomic colonies were quantified. *Data availability:* Raw colony counts, frequency calculations, and inferred population sizes are presented in Supplemental Table S5. *Visualization:* The frequency of revertant clones in each replicate monosomic population assessed are presented in Fig. 1D in a plot created in Graphpad Prism 10.

### Rate estimation using mlemur

#### Rate estimation

We used mlemur v0.9.6.5 (Łazowski 2023) to estimate the mutation rates for all the experimental fluctuation test data. The batch mode in the graphical user interface was used to input selective and non-selective colony counts, culture volumes, dilution factors and relative mutant fitness. *Data availability:* The colony count input and rate outputs for the monosome replating assay and canavanine fluctuation tests are presented in Supplemental Tables S6 and S8, respectively. *Visualization:* The rates of reversion estimated from the canavanine fluctuation tests and monosome replating tests are presented in Fig. 3A and B with plots created in Graphpad Prism. The comparative plot in Fig. 3C was made with a custom plotting script (see Data Availability).

### Relative fitness quantification

#### Preselected monosome assay

For monosomes of Chr1, Chr2, Chr3, Chr8, Chr9, Chr14, and Chr16, 3 independent 72hr grown light CAN^R^ clones were isolated and inoculated in 1mL of liquid rich YPD media. A diploid control strain constructed in the same hybrid genetic background was also inoculated in triplicate and all cultures were grown overnight at room temperature to saturation. Saturated cultures were then diluted 20-fold into fresh YPD and dispensed in duplicate into microwell dishes. *Induced monosome assay:* The pGAL1-CEN strains were grown overnight 1mL cultures of liquid uracil drop-out media containing 2% raffinose. These cultures were then diluted 3-fold into synthetic defined complete media containing 2% galactose and grown for 2.5hrs to induce centromere inactivation. At least 5 replicate 100-fold dilutions were seeded into 384-well microplates (Corning cat. 3680; Corning, New York) containing synthetic defined complete media containing 2% glucose and 1mg/L 5-FOA, in a total volume of 80 μL per well. *Optical density measurements:* Growth kinetics were tracked over 36 hrs by measuring the optical density (OD600) of each microwell every 20 min in a Cytation3 (BioTek) plate reader heated to 30°C without shaking. *Growth curve analysis:* For each replicate growth curve, the maximum growth rate was estimated using custom Python v3.10 (Van Rossum and Drake 2009) scripts. Linear regressions of OD600 values against time were fitted for overlapping sliding windows of 20 timepoints (6.67 hrs) using scipy v1.14 (Virtanen et al. 2020), and the maximum slope value was used. *Determination of fitness coefficients:* The average maximum slope values calculated from each replicate monosomic population of a given strain or isolate was divided by the average maximum slope value calculated from the replicate growth curves of the appropriate diploid control strain. *Data availability:* Raw slope values and fitness coefficients are presented in Supplemental Table S4. *Visualization:* The plot of inferred relative fitness coefficients and simple linear regression analysis presented in Fig. 2B was generated using Graphpad Prism 10.

### Whole Genome Sequencing (WGS) library preparation

The information for all clones sequenced in this study is deposited in Supplementary Table S9. WGS of selected clones was performed as described previously (Heasley et al. 2020; Heasley and Argueso 2022). Briefly, candidate colonies were isolated using sterile toothpicks, repatched to rich media plates, and expanded for one day. The entire patch of cells was then used to isolate genomic DNA using the Yeastar Genomic Kit from Zymo Research (cat. D2002; Irvine, CA). DNA quantity and quality was assessed using a Qubit fluorometer (Thermo Fisher Scientific). Pooled, barcoded libraries of 96-384 individual genomes were generated using a Seqwell plexWell-384 kit. The final barcoded libraries were sequenced at Novogene Corp. using an Illumina Novaseq sequencer.

### WGS data analysis

#### Read processing and trimming

Reads were trimmed using fastp v0.23.4 (Chen et al. 2018) with default options. *Read mapping:* Reads were mapped to the *S. cerevisiae* reference genome vR64 (http://sgd-archive.yeastgenome.org/sequence/S288C_reference/genome_releases/) using bwa-mem2 v2.3 (Vasimuddin et al. 2019) with default options. Secondary alignments were filtered out using samtools v1.21 (Li et al. 2009). Picard v3.1.1 (http://broadinstitute.github.io/picard) was used to mark and remove duplicates (MarkDuplicates) and add read groups to the BAM files (AddOrReplaceReadGroups). *Variant detection:* Variants were called using bcftools v1.21 (Li 2011), first generating pileups (mpileup) and subsequently calling the variants (call) with options -m -v. Biallelic single nucleotide variants were kept for the genome analyses. *Data availability:* Per chromosome assessment of the total clones sequenced and the percentage of which harbor defined karyotypic variations is presented in Supplemental Table S2. Raw sequencing data are deposited at NCBI under Bioproject PRJNA1414607.*Visualization:* Figures 1C, 5A, 5B, 5G, 5H, 6B, 6C, and 6E were made with custom scripts deposited on this study’s Github page (Data Availability)

### Monosome copy number analysis

#### Genomic analysis

The WGS data derived from each of the light CAN^R^ clones which harbored only a monosome of the marked chromosome were remapped to a repeat masked reference assembly. The use of a masked reference eliminated the possibility that mismapping to repetitive regions of the genome (*i.e.*, the rDNA array, subtelomeric repeats, mobile elements) would distort coverage analyses and allowed us to more precisely calculate the normalized copy number of the marked chromosome relative to the rest of the genome. The average depth of coverage of the respective marked chromosome was divided by the mean coverage of the other 15 chromosomes and normalized to relate a copy number value. *Data Availability:* Per clone normalized copy number values are presented in Supplemental Table S3. *Visualization:* The deduced copy numbers of each monosome analyzed are presented in Fig. 1D in a plot created in Graphpad Prism 10.

### Forward simulation of fluctuation assays

We developed fflucsim, a forward simulation tool to generate in silico populations expanding from a single diploid cell ancestor. fflucsim is a Python library that includes objects and functions to generate cells and expand them into large populations under adjustable parameters dictating the population dynamics that generate monosomic karyotypes and the mutational events immediately following that revert to disomic karyotypes. Briefly, founder cells are initiated with user-specified values for aneuploidy rates and relative fitness of aneuploid karyotypes. Cells divide synchronously (Sarkar 1991) using relative fitness and mutation rates as probabilities of karyotype change and generation of a daughter cell, respectively, until a user-specified target population size is reached. Cells implement two distinct homologs, one carrying a counter-selectable marker, which enables the selection of surviving cells at the end of the population expansion (mimicking the *CAN1* cassettes in our experimental fluctuation assays). Importantly, fflucsim keeps full records of the parameters, mutation events and full genealogy in cell and population objects, making it a uniquely powerful framework to track the dynamics of complex mutations with alternative paths to back-mutation. The source code and the Snakemake v9.15 (Köster and Rahmann 2012) scripts used for all simulations presented in this study are available at GitHub (see Data accessibility section).

### Microscopy

#### Growth conditions

Cells were pregrown to log phase in synthetic-defined complete media containing 2% raffinose, before dilution into complete media containing either 2% glucose or 2% galactose. After 90 min of growth, cells were prepared for imaging. *Slide preparation:* Cells were concentrated, spotted on a 1.5% agarose pad containing the same respective media, covered with a coverslip and sealed with paraffin. *Imaging procedures:* Images were collected on a Nikon Ti-E microscope equipped with a 1.45NA100 ×CFIPlan Apoobjective, piezo electric stage (Physik Instrumente; Auburn, MA), spinning disk confocal scanner unit (CSU10; Yokogawa), 488-nm and 561-nm lasers (Agilent Technologies; Santa Clara, CA), and an EMCCDcamera (iXonUltra 897;AndorTechnology, Belfast, UK) using NIS Elements software (Nikon). Z-series spanning 7-9 μm with 0.5 μm steps were acquired every 3min for 2 hours. *Data Availability*: Raw image files will be provided upon request. *Visualization*: Image files were imported as hyperstacks and processed using Fiji (Schindelin et al. 2012). Maximum projections of z-series for each channel were generated. Intensity and color adjustments, cropping, and merging was performed with tools in either Fiji or Adobe Photoshop.

---

**Figure S1.**
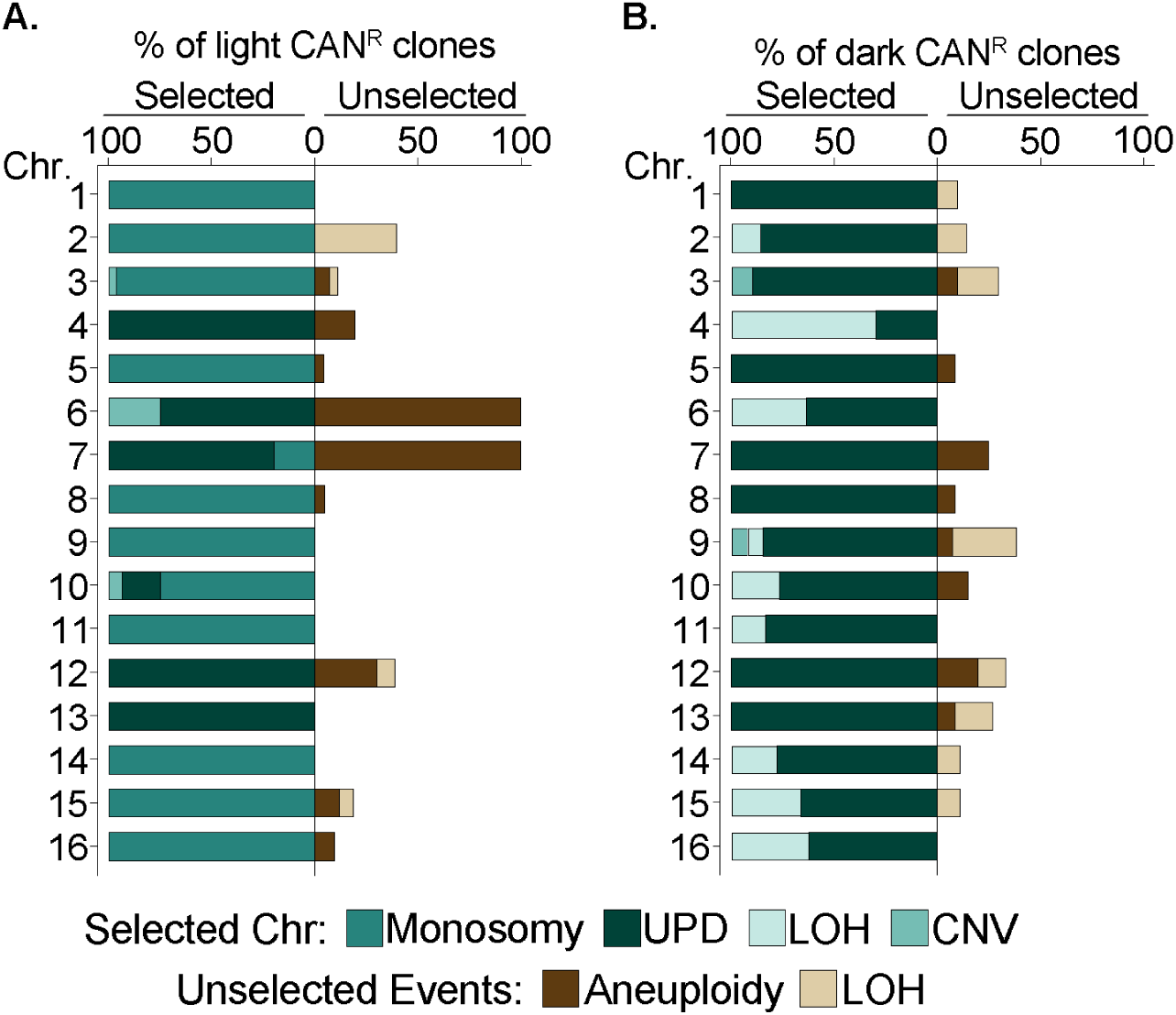
Karyotypic alterations associated with the light and dark CAN^R^ phenotype on a per-chromosome basis. A, **B.** Quantification of the percentage of light CAN^R^ (**A**) and dark CAN^R^ (**B**) clones that harbored the denoted karyotypic states affecting the predicted chromosome (Selected) as well as any additional unselected events (Unselected).

**Figure S2.**
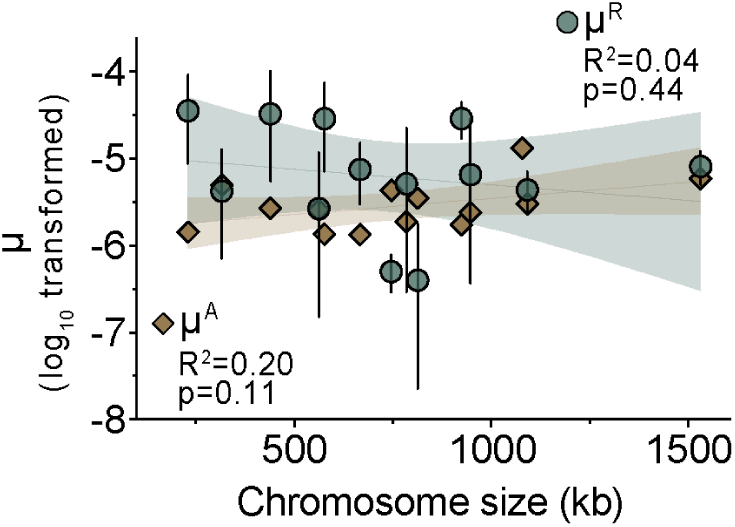
Mutation rates are not correlated with chromosome size. Log-transformed per-chromosome estimates of the rates of monosomy and reversion plotted by chromosome size. Vertical error bars denote the 95% confidence intervals associated with each per-chromsome estimate of μ^A^ or μ^R^. Brown and green areas depict the 95% confidence intervals produced from the simple linear regression of μ^A^ (brown) μ^R^ (green) or vs. chromosome size.

**Figure S3.**
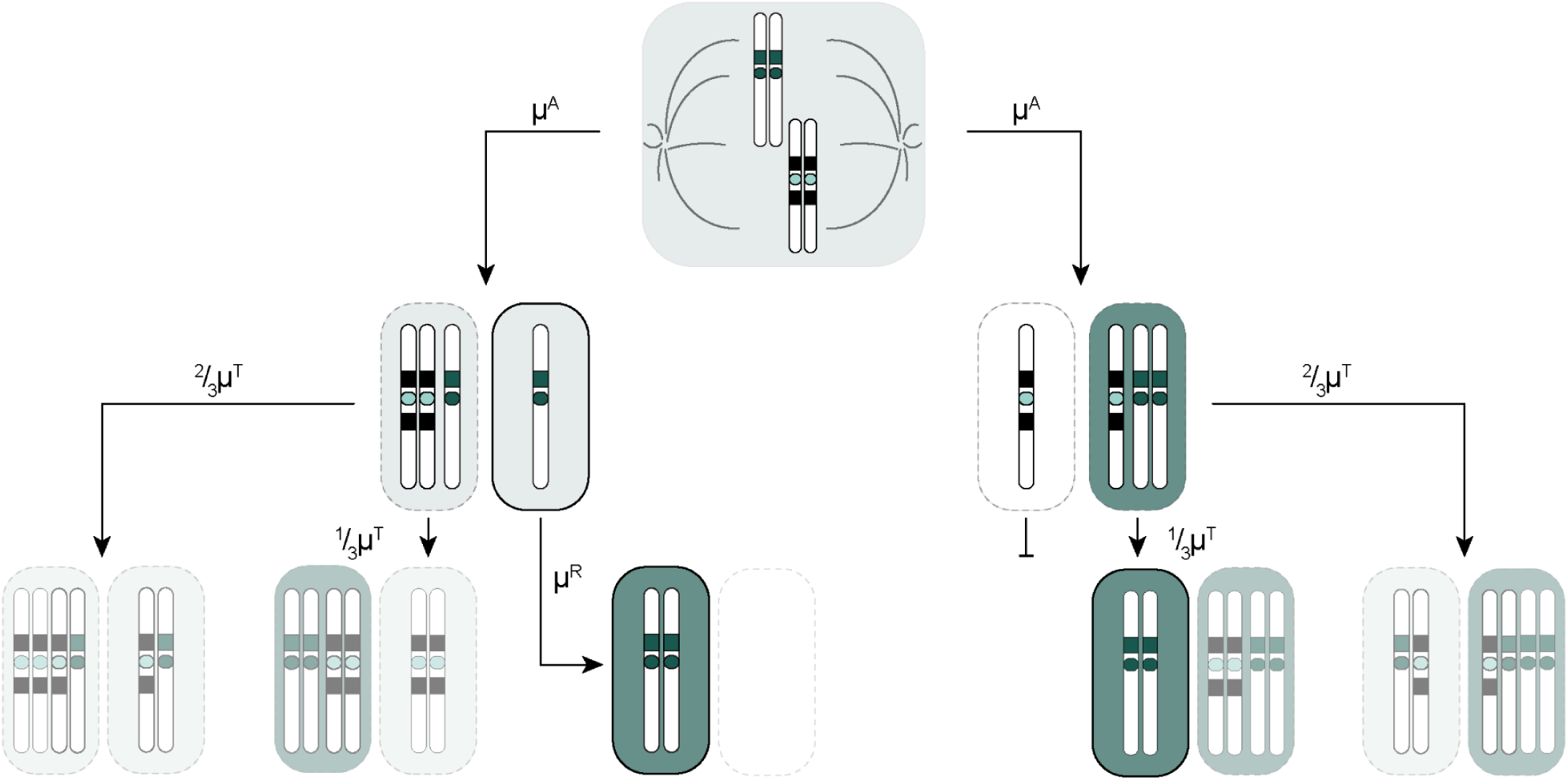
Parallel mutational routes can generate revertant UPDs. A schematic illustrating genotypes, phenotypes, and rates associated with nondisjunction events that result in the formation and subsequent reversion of monosomic and trisomic daughter cell pairs. Symbols are the same as in Fig. 1. Only the derivatives outlined with a solid line would be recoverable in the canavanine fluctuation tests.

**Figure S4.**
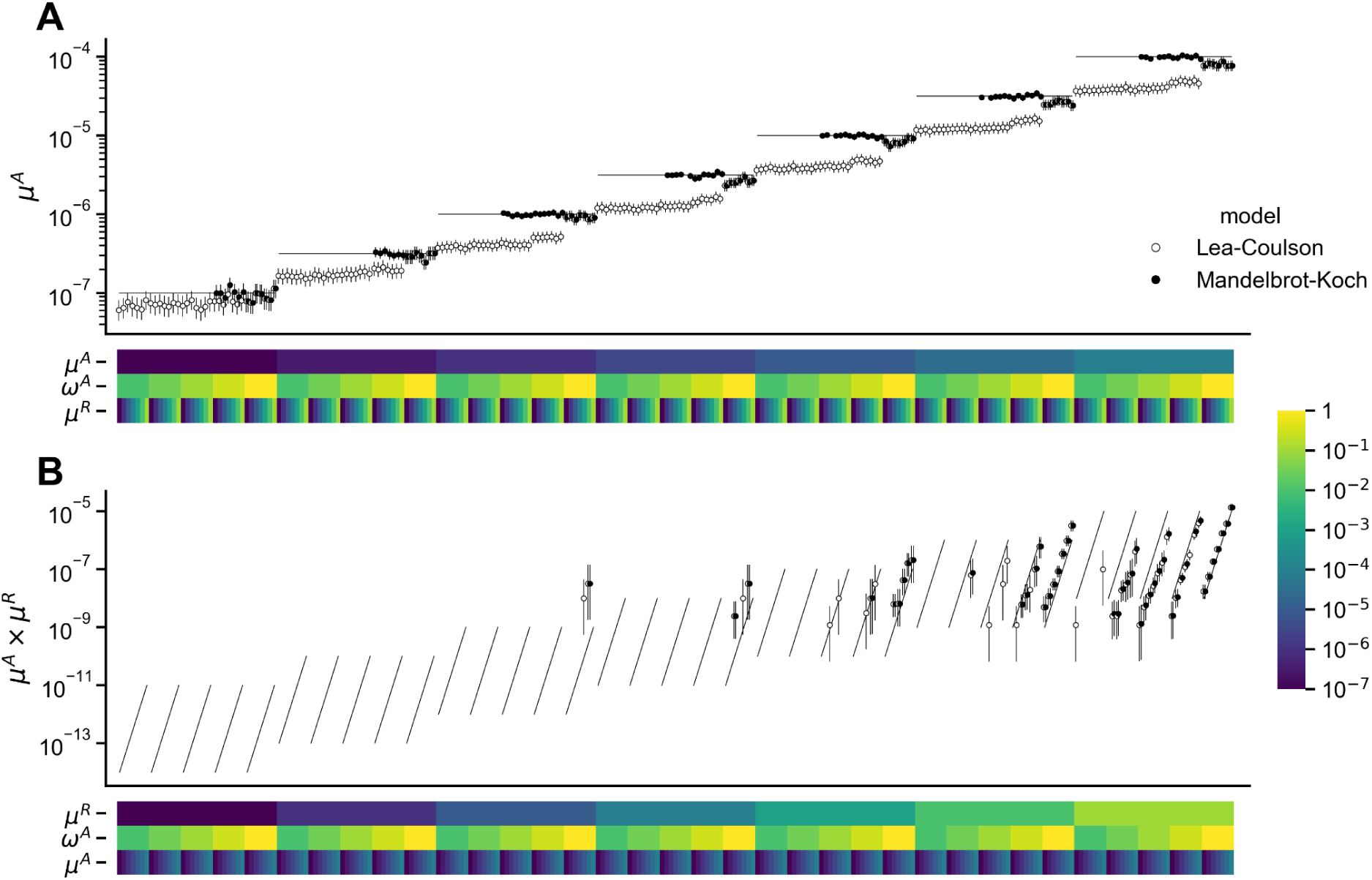
Validation of fflucsim using back-calculation of mutation rates from simulated mutant counts. Mutation rates (dots) and 95% confidence intervals (vertical lines) were estimated from simulated mutant counts using rSalvador (Zheng 2017). We performed 245 simulations across all combinations of seven μ^A^ values (10^-7^–10^-4^), seven μ^R^ values (10^-7^–10^-1^) and five ω^A^ values (10^-2^–10^0^). For these validation simulations, we ignored reversion that occurs via the trisomic counterparts of monosomy events, setting μ^T^=0 and ω^T^=1. 50 replicates for each parameter combination. Two models were fitted on simulated mutant counts, the standard Luria-Delbrück model (Lea-Coulson, open dots) and the model accounting for variable mutant fitness (Mandelbrot-Koch, closed dots). The rate values expected from the input parameters are shown as thin background lines. Input parameters are summarized as heatmaps below each plot. Monosome rates (μ^A^) were estimated from combined counts of monosomic and revertant mutants (**A**), and compound rates (μ^C^ = μ^A^ ⨉ μ^R^) were estimated from revertant mutant counts alone (**B**).

**Fig S5.**
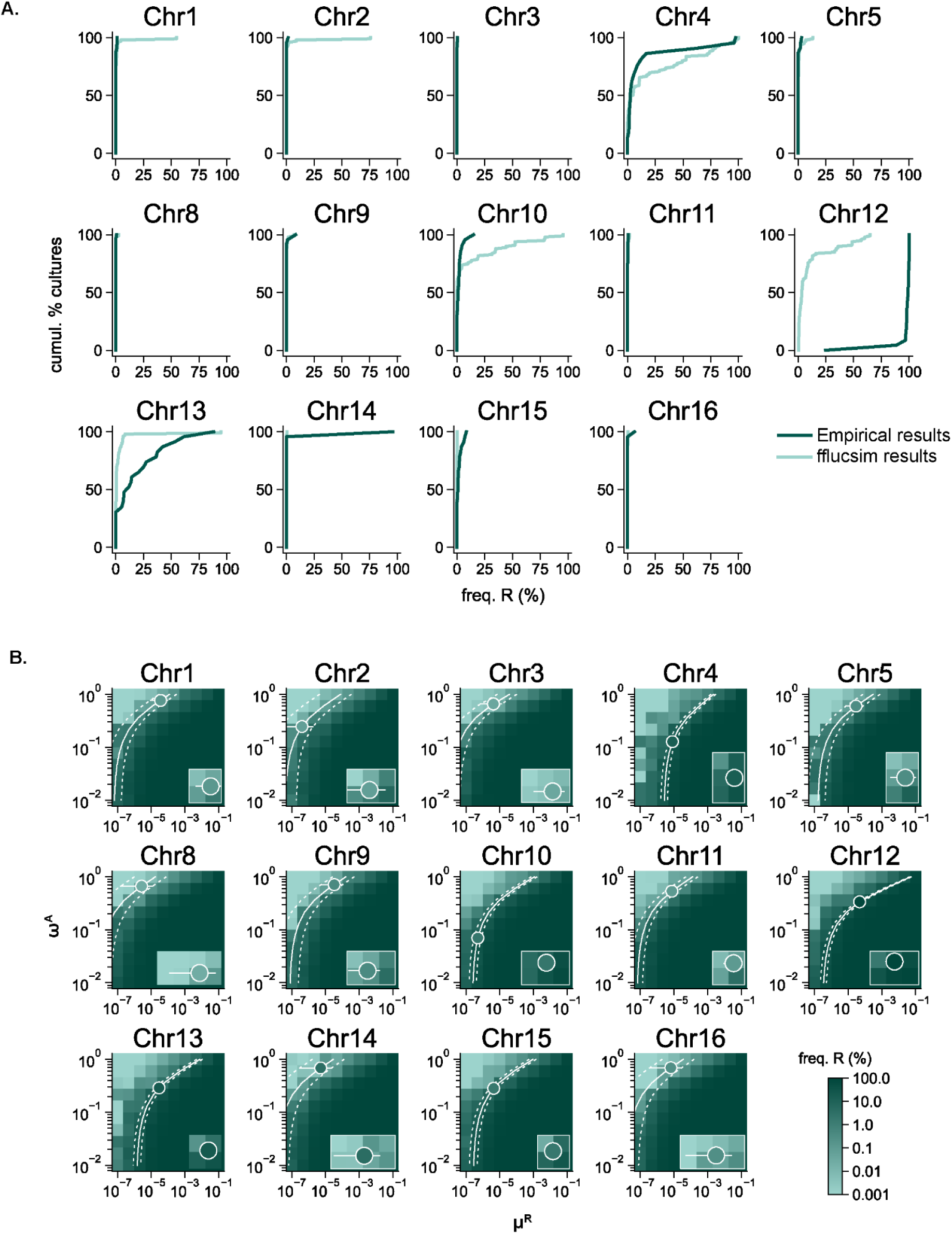
Empirical revertant frequencies deviate substantially from the expected value for a subset of chromosomes. **A.** Cumulative distributions of revertant cell (R) frequency for the monosome replating tests (dark green) and simulated populations generated by fflucsim (light green). The fflucsim distributions correspond to the 50 replicates of the single simulated assay parameterized with the pair of μ^R^ and ω^A^ values closest to the empirical estimates for the corresponding chromosome. **B.** Heatmaps showing the same data as Fig. 3D with revertant frequency mapped on a logarithmic scale. Insets highlight the set of input parameters closest to the empirical estimates.

**Figure S6.**
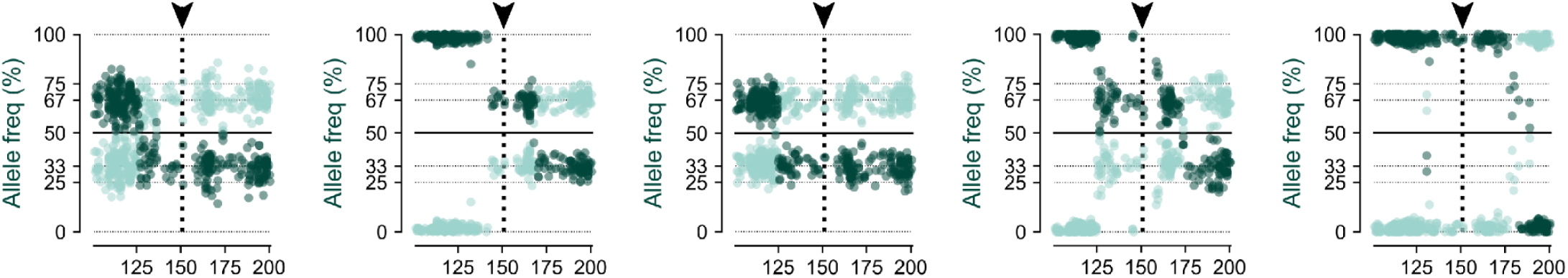
Pericentromeric recombination patterns suggest an intermolecular linkage-mediated mechanism of chromosome missegregation. Chr12-specific whole-chromosome or centromere-proximal WGS mappings of the recovered trisomic CAN^R^ colonies. Black arrowheads in denote the position of *CEN12*.

**Figure S7.**
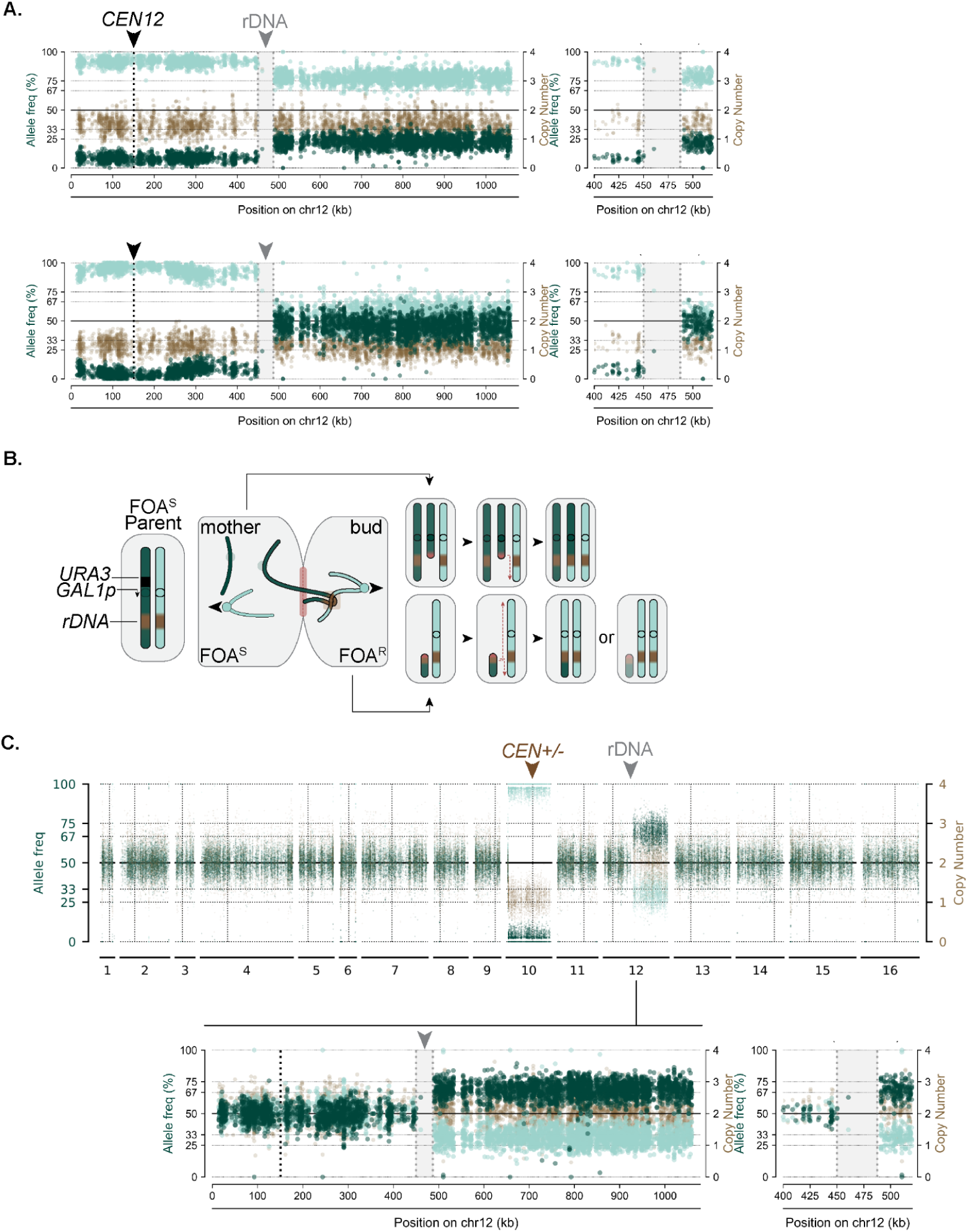
Breaks within the ribosomal DNA coincide with FOA^R^ karyotype formation. **A.** Representative Chr12-specific WGS mappings from recombinant FOA^R^ colonies derived from the *URA3-GAL1p-CEN12* strain displaying a breakpoint within the rDNA locus. **B.** A schematic illustrating the predicted mechanism and outcomes of a nondisjunction event resulting from intermolecular linkages arising in the rDNA. **C.** A representative WGS mapping and corresponding Chr12-specific WGS mapping of a FOA^R^ clone recovered from the *URA3-GAL1p-CEN10* strain. As shown in the top mapping, this clone is monosomic for Chr10 and also harbors a mosaic structural variation involving the right arm of Chr12 beginning at a breakpoint in the rDNA. The bottom mapping more clearly illustrates the rDNA breakpoint and mosaic allele frequency on Chr12.

**Figure S8.**
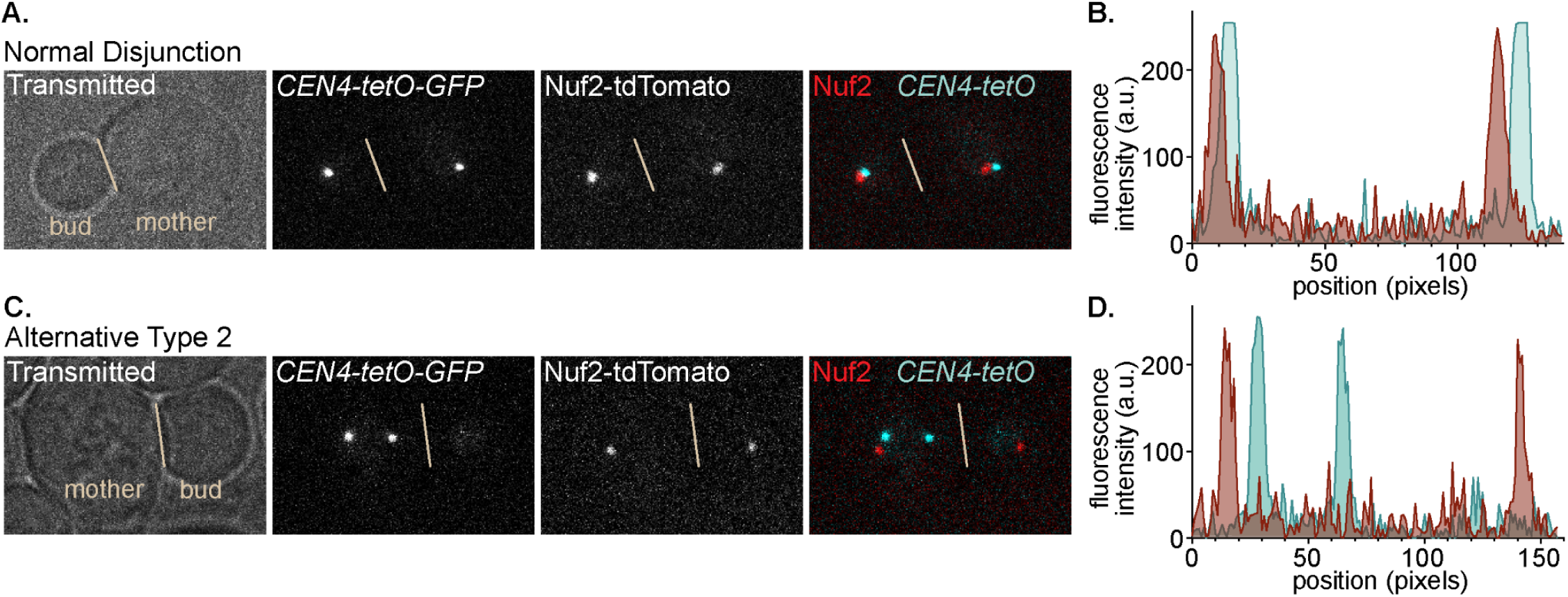
Alternative missegregation occurs in the absence of intact kinetochores. A. and. **C.** Representative images of a normal disjunction event (**A.**) and an alternative Type 2 missegregation event (**C.**) occurring in *URA3-GAL1p-CEN4-tetO* cells expressing TetR-GFP and Nuf2-tdTomato, a kinetochore protein required for spindle microtubule attachment. **B. and D.** Corresponding line scan analysis illustrating the degree of TetR and Nuf2 signal overlap in **A.** and **C.**; Line scan analysis was performed by determining the pixel intensity across the distance spanning all four fluorescent foci for both detection channels using Fiji.

**Supplemental Table 1.**
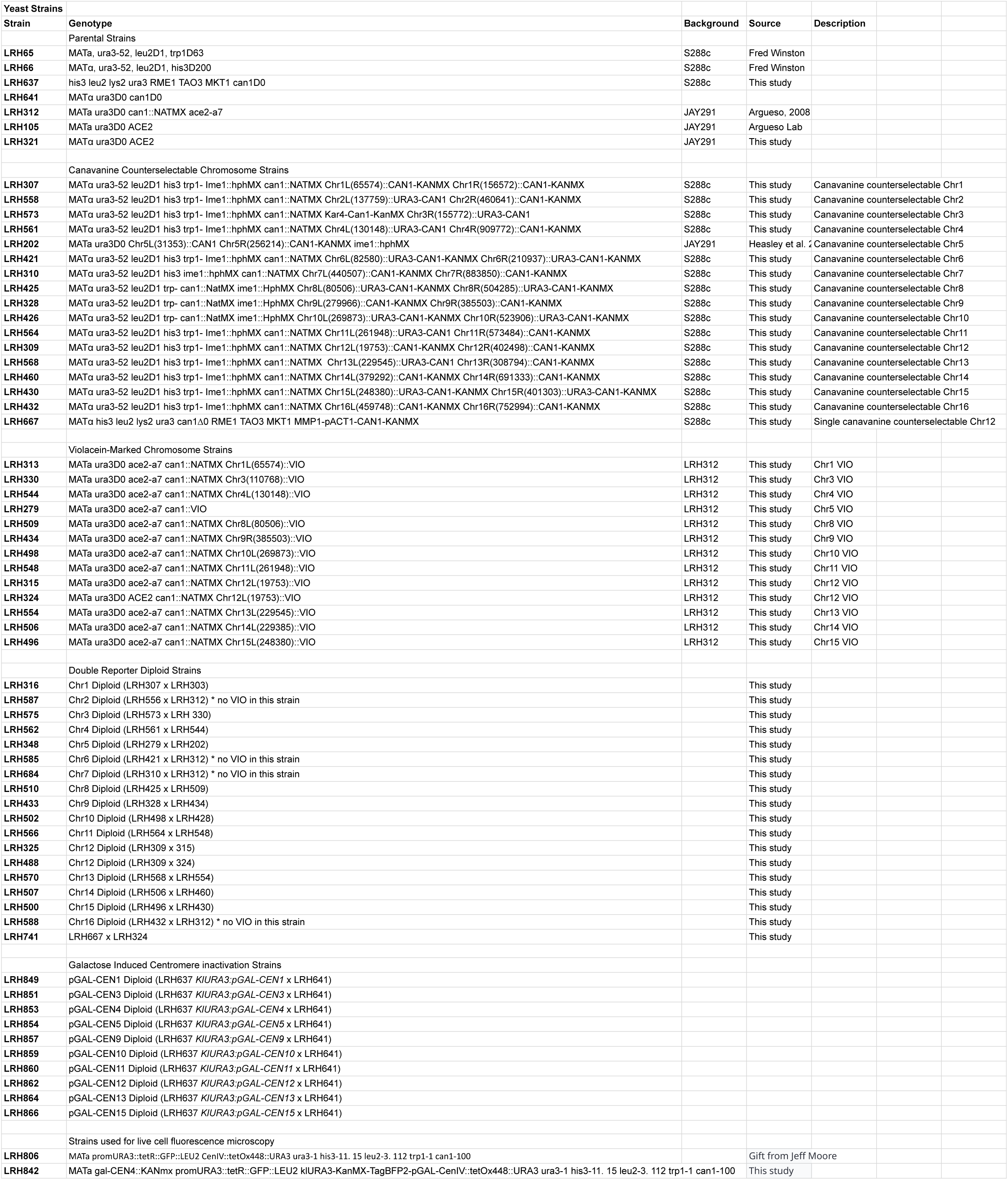
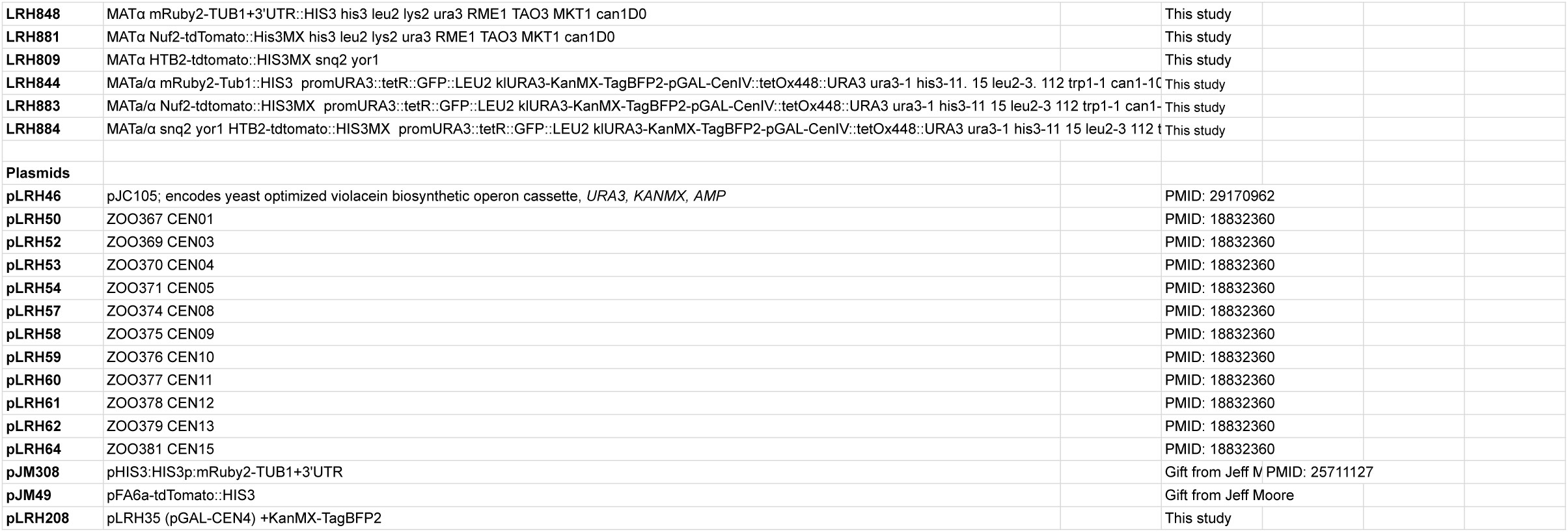
Yeast strains and plasmids used in this study.

**Supplemental Table 2.**
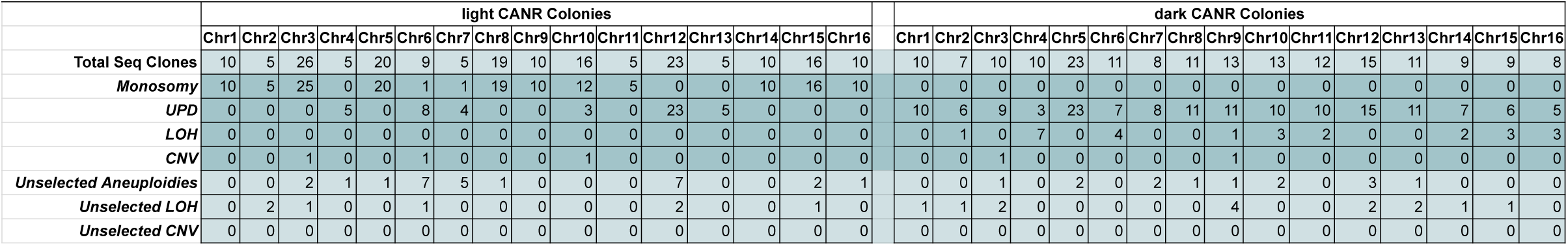
Karyotypic variant analysis of sequenced clones.

**Supplemental Table 3.** Per chromosome monosome coverage analysis.

|  | Chr1 | Chr2 | Chr3 | Chr4 | Chr5 | Chr8 | Chr9 | Chr10 | Chr11 | Chr12 | Chr13 | Chr14 | Chr15 | Chr16 |
| --- | --- | --- | --- | --- | --- | --- | --- | --- | --- | --- | --- | --- | --- | --- |
| Per clone normalized copy number | 0.7996355 | 1.091706 |  | 2.0098828 | 1.118964 | 1.032545 | 1.01113 | 1.537517 | 1.828786 | 2.053768 | 1.9704103 | 1.159009 | 1.631311 | 1.031625 |
|  | 0.807015 | 1.0843983 | 1.22684 | 2.0086727 | 1.235367 | 1.038313 | 1.004716 | 1.651996 | 1.79463 | 2.022773 | 1.8772777 | 1.119829 | 1.58355 | 1.021552 |
|  | 0.796933 | 1.0825203 | 1.220532 | 2.0244443 | 1.149871 | 1.0292 | 0.995749 | 1.798694 | 1.882756 | 2.0116032 | 1.9000951 | 1.101699 | 1.743736 | 1.023892 |
|  | 0.788338 | 1.0939787 | 1.177189 | 2.0364003 | 1.210567 | 1.037618 | 0.990973 | 1.877019 | 1.789194 | 2.014883 | 2.0300854 | 1.08766 | 1.648541 | 1.056429 |
|  | 0.798602 | 1.0808513 | 1.217993 | 2.0395163 | 1.136041 | 1.030592 | 1.00348 | 1.717759 | 1.784398 | 2.0472376 | 1.9015799 | 1.101069 | 1.611913 | 1.022325 |
|  | 0.798111 |  | 1.225919 |  | 1.280664 | 1.032074 | 0.982545 | 1.767735 |  | 2.0185205 |  | 1.091315 | 1.578744 | 1.016297 |
|  | 0.799111 |  | 1.224775 |  | 1.112261 | 1.021791 | 0.992242 | 1.670493 |  | 2.0185308 |  | 1.0946 | 1.391004 | 1.016297 |
|  | 0.786409 |  | 1.21746 |  | 1.267941 | 1.041932 | 0.991771 | 1.835839 |  | 2.0417014 |  | 1.141936 | 1.584838 | 1.01975 |
|  | 0.778323 |  | 1.14506 |  | 1.214538 | 1.023969 | 0.990954 | 1.848282 |  | 1.9673312 |  | 1.105422 | 2.006343 | 1.015293 |
|  | 0.783348 |  | 1.133152 |  | 1.335603 | 1.027902 | 0.992042 | 1.761189 |  | 2.026438 |  | 1.133963 | 1.820668 | 1.099354 |
|  |  |  | 1.131376 |  |  | 0.999332 |  | 1.858134 |  | 2.19089 |  |  | 1.627134 |  |
|  |  |  | 1.135568 |  |  | 1.021861 |  | 1.80463 |  | 2.04739 |  |  | 1.752769 |  |
|  |  |  | 1.159232 |  |  | 1.029482 |  | 1.747778 |  | 2.192134 |  |  | 1.904536 |  |
|  |  |  | 1.192126 |  |  | 1.026935 |  | 1.742224 |  | 2.127509 |  |  | 1.827962 |  |
|  |  |  | 1.135591 |  |  | 1.029476 |  | 1.844197 |  | 2.138892 |  |  | 1.788349 |  |
|  |  |  | 1.172775 |  |  | 1.016152 |  | 1.857504 |  | 2.177243 |  |  | 1.851202 |  |
|  |  |  | 1.338019 |  |  | 1.021938 |  |  |  | 2.126525 |  |  | 1.866673 |  |
|  |  |  |  |  |  | 1.056791 |  |  |  | 2.073156 |  |  | 1.872951 |  |
|  |  |  |  |  |  | 1.029805 |  |  |  |  |  |  | 1.695855 |  |
|  |  |  |  |  |  | 1.004805 |  |  |  |  |  |  |  |  |
|  |  |  |  |  |  | 0.966978 |  |  |  |  |  |  |  |  |
|  |  |  |  |  |  | 1.007999 |  |  |  |  |  |  |  |  |
|  |  |  |  |  |  | 1.006509 |  |  |  |  |  |  |  |  |
|  |  |  |  |  |  | 1.237301 |  |  |  |  |  |  |  |  |
|  |  |  |  |  |  | 1.015917 |  |  |  |  |  |  |  |  |
|  |  |  |  |  |  | 0.998102 |  |  |  |  |  |  |  |  |
|  |  |  |  |  |  | 1.01846 |  |  |  |  |  |  |  |  |
|  |  |  |  |  |  | 0.993069 |  |  |  |  |  |  |  |  |

**Supplemental Table 4.** Data used to calculate relative fitness coefficients.

| strain | growth conditions | dataset | average slope across replicates | relative fitness coefficient (w) |
| --- | --- | --- | --- | --- |
| Diploid | ypd | preselected monosome | 0.15325525 | 1 |
| chr1_mono1 | ypd | preselected monosome | 0.12382518 | 0.80796697 |
| chr1_mono2 | ypd | preselected monosome | 0.1229021 | 0.80194382 |
| chr1_mono3 | ypd | preselected monosome | 0.12976573 | 0.84672943 |
| chr2_mono1 | ypd | preselected monosome | 0.07163986 | 0.46745452 |
| chr2_mono2 | ypd | preselected monosome | 0.0479965 | 0.31318014 |
| chr2_mono3 | ypd | preselected monosome | 0.0275979 | 0.18007801 |
| chr3_mono1 | ypd | preselected monosome | 0.0951958 | 0.62115849 |
| chr3_mono2 | ypd | preselected monosome | 0.10685315 | 0.69722342 |
| chr3_mono3 | ypd | preselected monosome | 0.08958042 | 0.58451779 |
| chr9_mono1 | ypd | preselected monosome | 0.11943706 | 0.77933422 |
| chr9_mono2 | ypd | preselected monosome | 0.11726573 | 0.76516615 |
| chr9_mono3 | ypd | preselected monosome | 0.11322028 | 0.73876934 |
| chr14_mono1 | ypd | preselected monosome | 0.10458042 | 0.68239372 |
| chr14_mono2 | ypd | preselected monosome | 0.11020979 | 0.71912571 |
| chr14_mono3 | ypd | preselected monosome | 0.10097203 | 0.65884875 |
| chr16_mono1 | ypd | preselected monosome | 0.10924476 | 0.71282883 |
| chr16_mono2 | ypd | preselected monosome | 0.10456643 | 0.68230243 |
| chr16_mono3 | ypd | preselected monosome | 0.14008741 | 0.91407903 |
| Diploid | SD_FOA | induced monosome | 0.17776224 | 1 |
| cen1 | SD_FOA | induced monosome | 0.10507552 | 0.59110146 |
| cen3 | SD_FOA | induced monosome | 0.09698601 | 0.54559399 |
| cen4 | SD_FOA | induced monosome | 0.0166479 | 0.09365262 |
| cen5 | SD_FOA | induced monosome | 0.08880525 | 0.49957319 |
| cen9 | SD_FOA | induced monosome | 0.09082448 | 0.51093236 |
| cen10 | SD_FOA | induced monosome | 0.01125315 | 0.0633045 |
| cen11 | SD_FOA | induced monosome | 0.07437168 | 0.41837727 |
| cen12 | SD_FOA | induced monosome | 0.05091504 | 0.28642213 |
| cen13 | SD_FOA | induced monosome | 0.04152273 | 0.23358577 |
| cen15 | SD_FOA | induced monosome | 0.05432518 | 0.30560585 |

**Supplemental Table 5.**
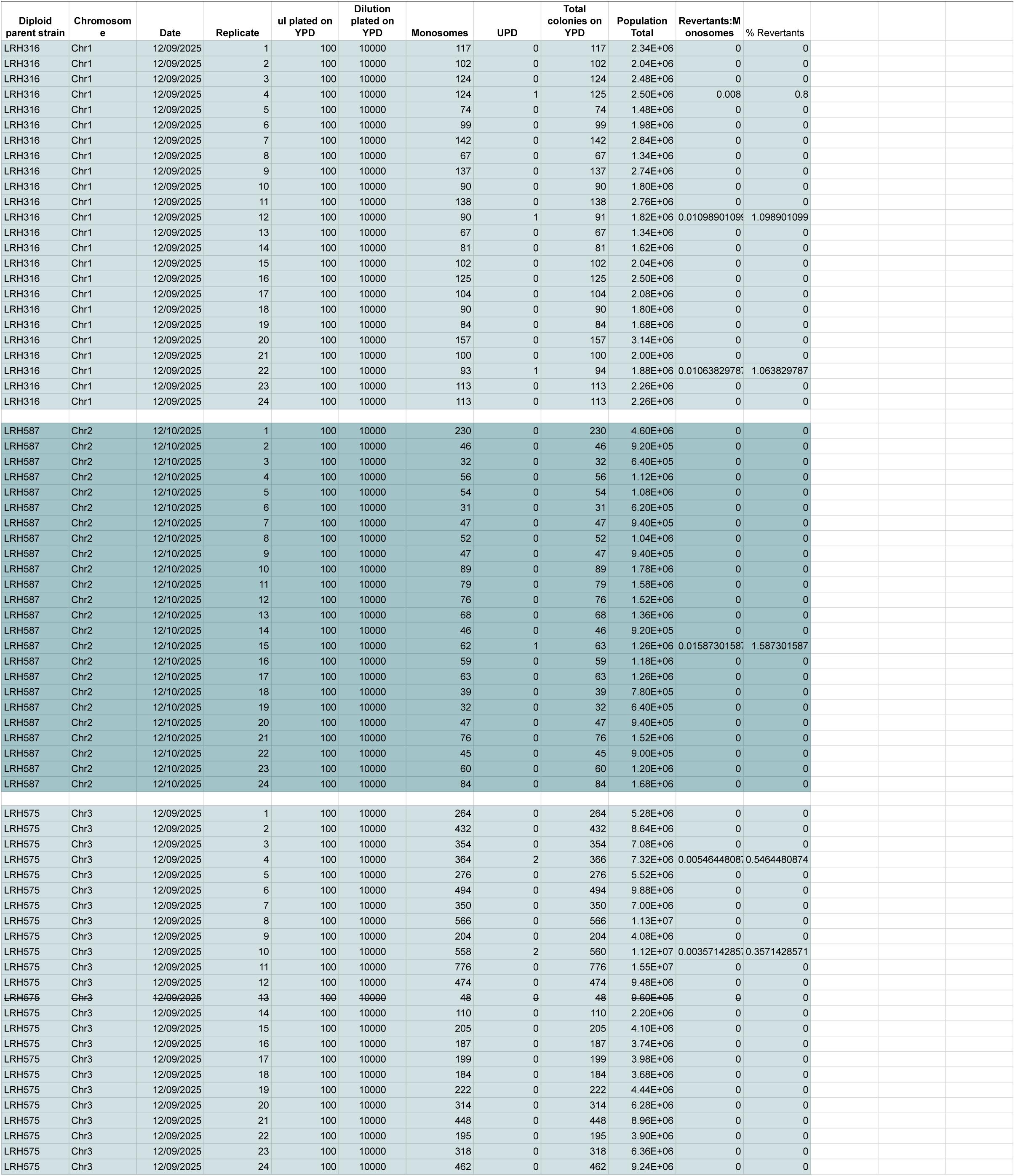
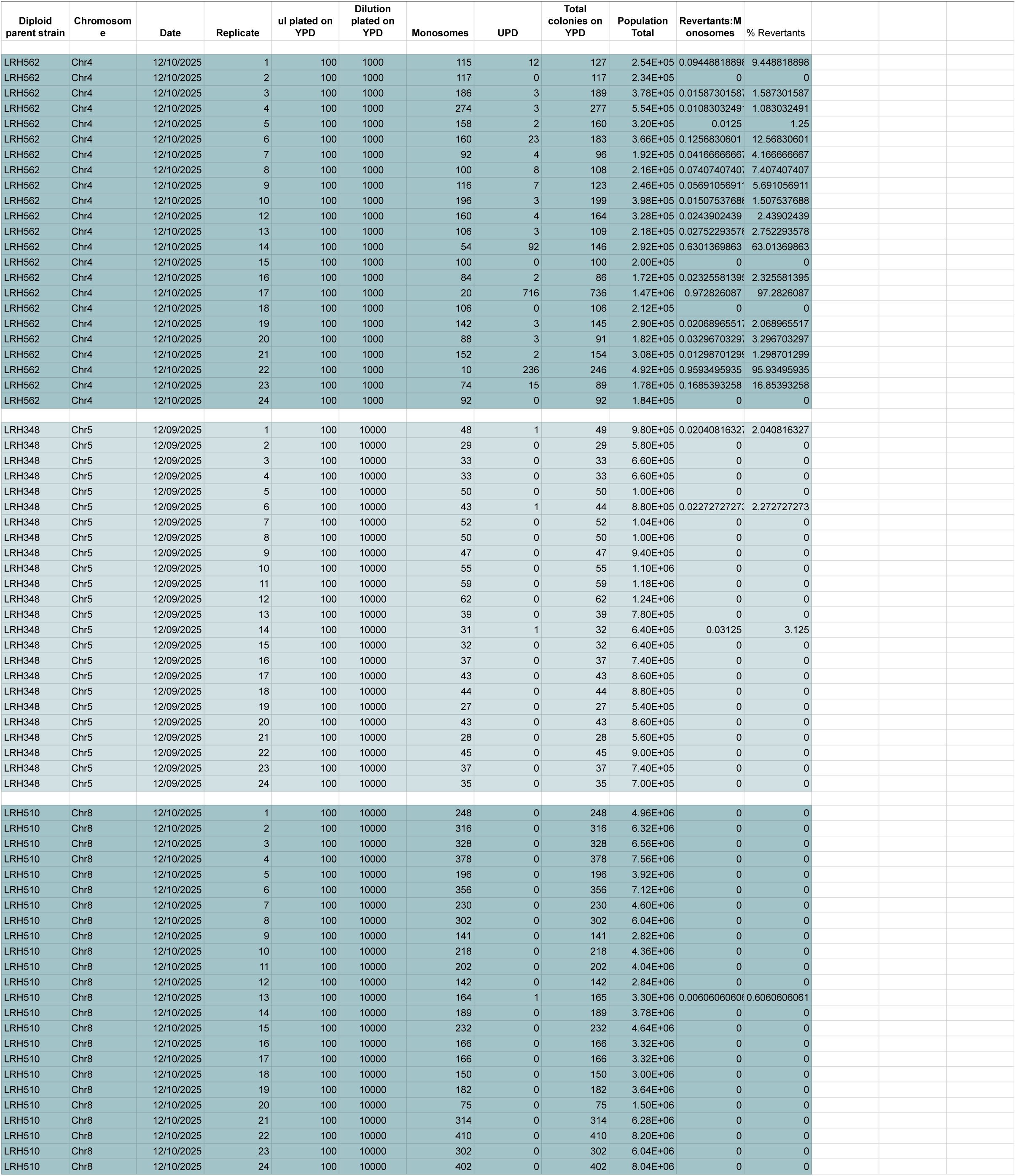
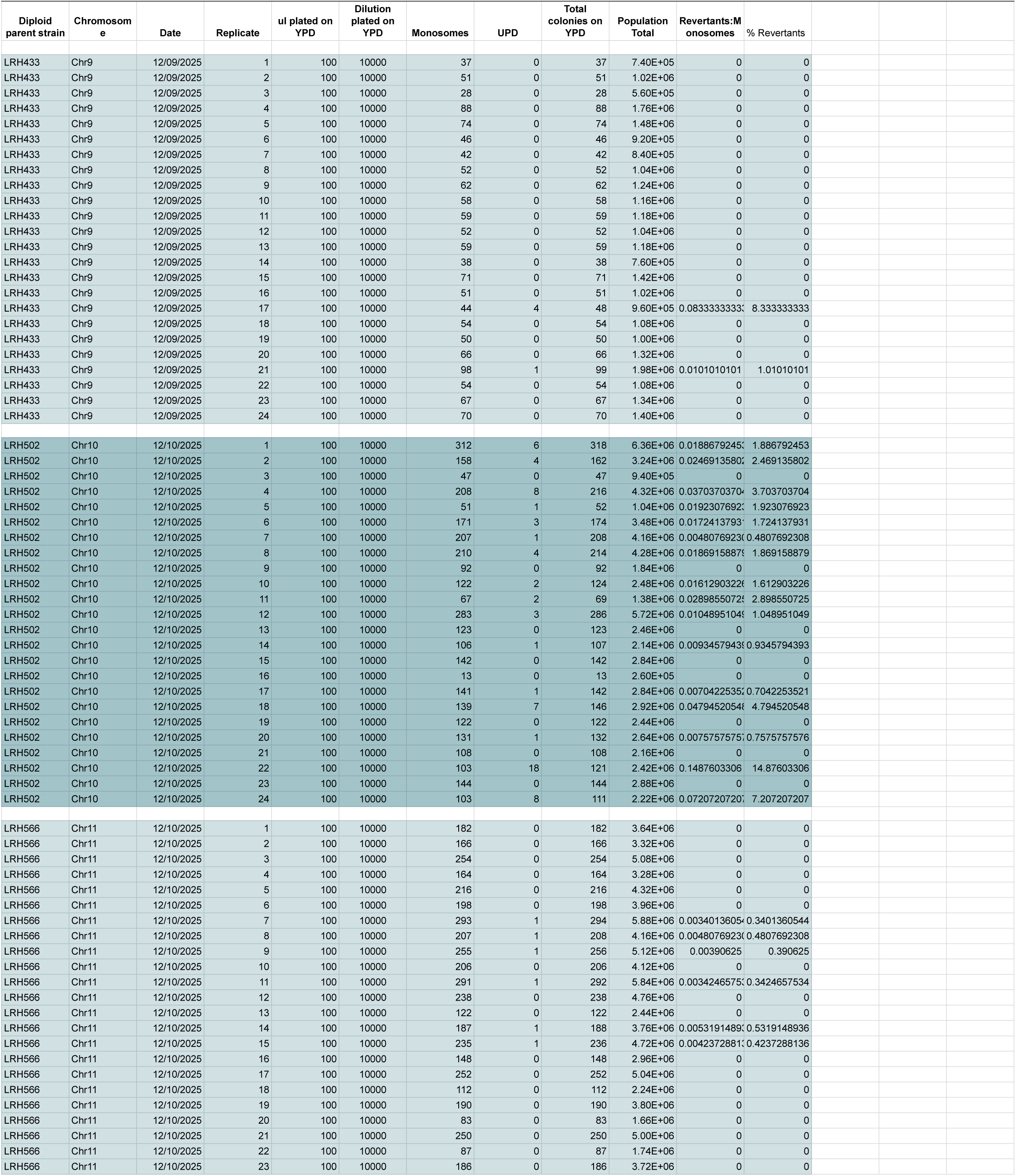
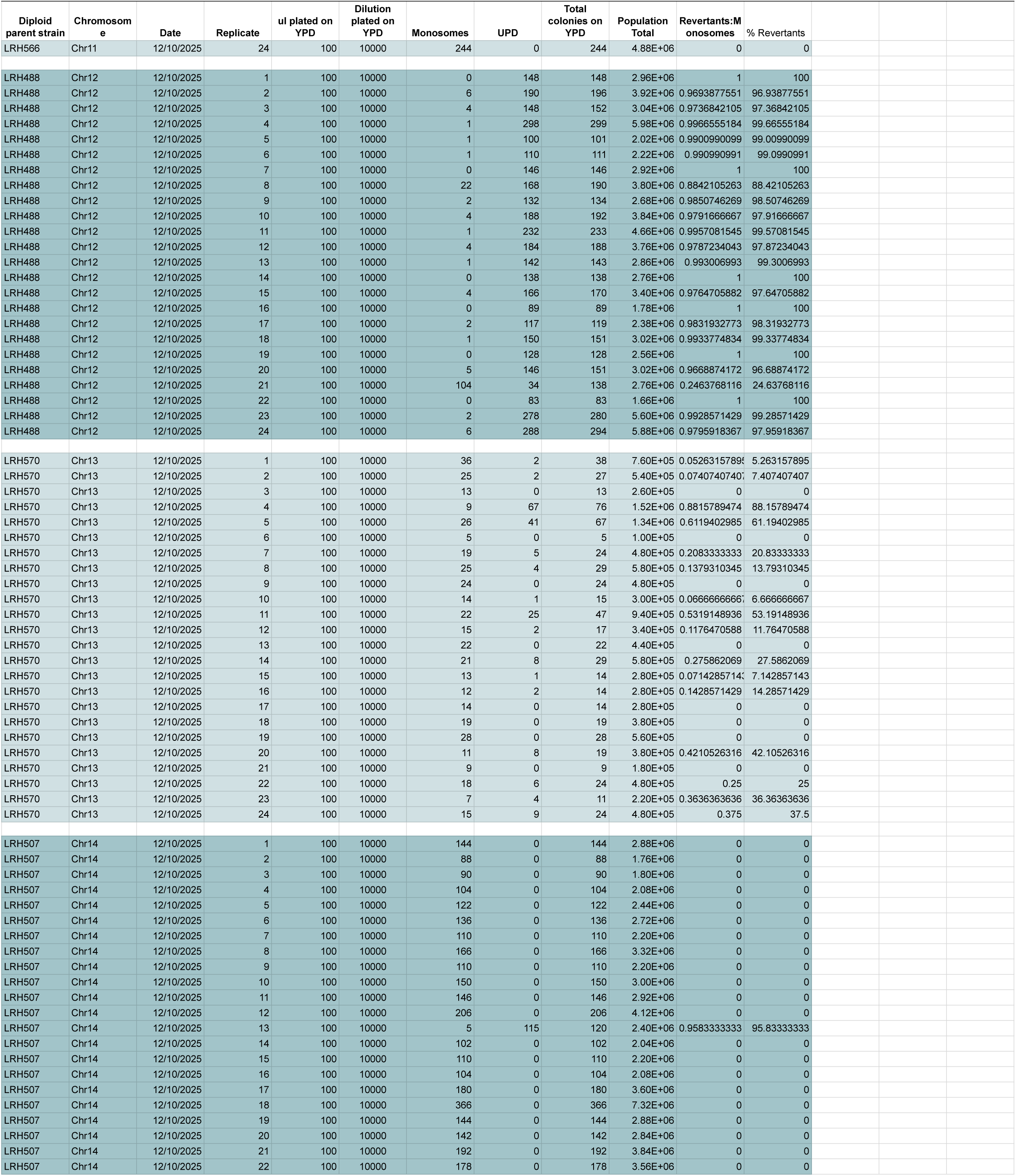
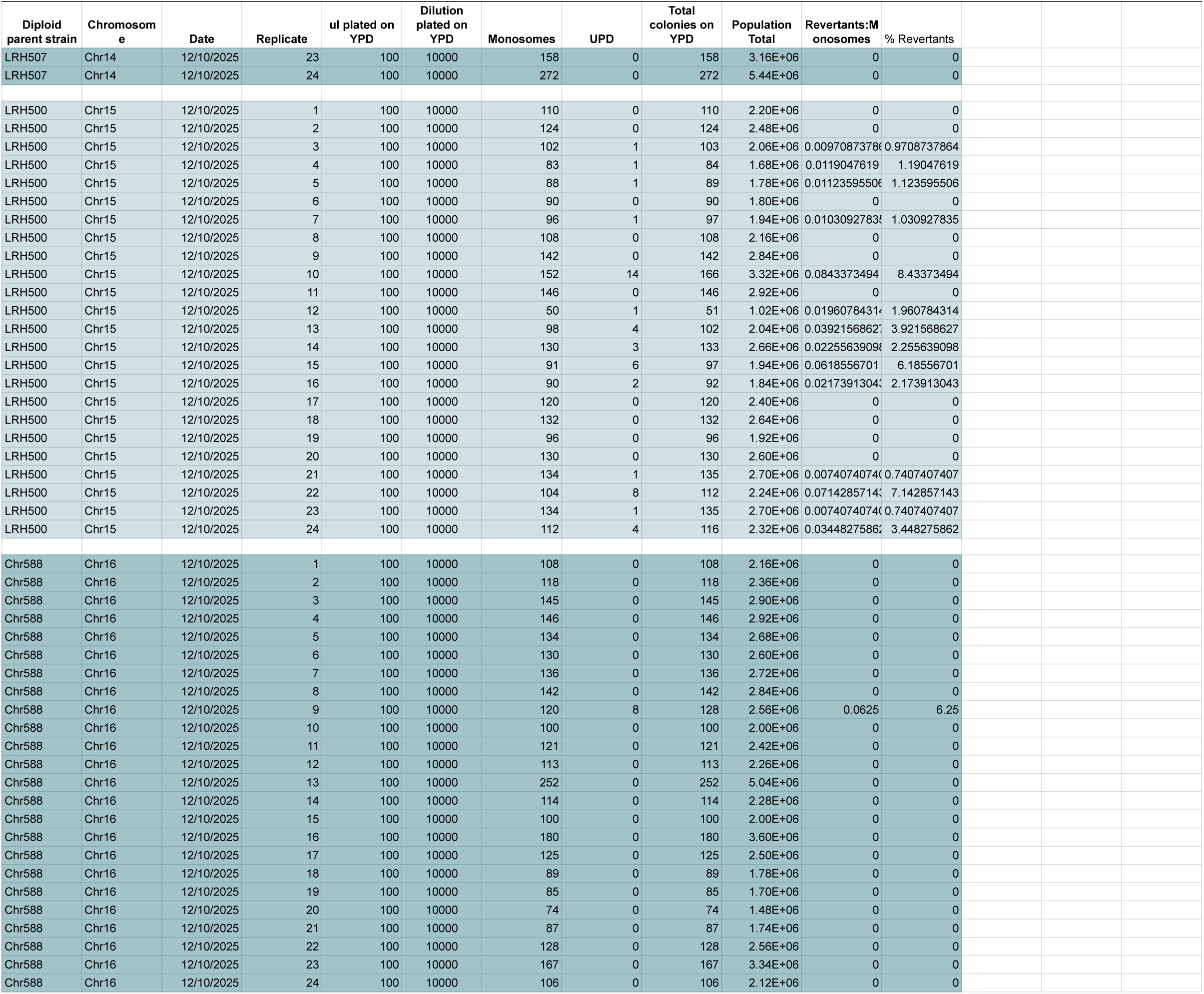
Monosome replating test count data.

| Diploid parent strain | Chromosome | Date | Replicate | ul plated on YPD | Dilution plated on YPD | Monosomes | UPD | Total colonies on YPD | Population Total | Revertants:Monosomes | % Revertants |
| --- | --- | --- | --- | --- | --- | --- | --- | --- | --- | --- | --- |
| LRH316 | Chr1 | 12/09/2025 | 1 | 100 | 10000 | 117 | 0 | 117 | 2.34E+06 | 0 | 0 |
| LRH316 | Chr1 | 12/09/2025 | 2 | 100 | 10000 | 102 | 0 | 102 | 2.04E+06 | 0 | 0 |
| LRH316 | Chr1 | 12/09/2025 | 3 | 100 | 10000 | 124 | 0 | 124 | 2.48E+06 | 0 | 0 |
| LRH316 | Chr1 | 12/09/2025 | 4 | 100 | 10000 | 124 | 1 | 125 | 2.50E+06 | 0.008 | 0.8 |
| LRH316 | Chr1 | 12/09/2025 | 5 | 100 | 10000 | 74 | 0 | 74 | 1.48E+06 | 0 | 0 |
| LRH316 | Chr1 | 12/09/2025 | 6 | 100 | 10000 | 99 | 0 | 99 | 1.98E+06 | 0 | 0 |
| LRH316 | Chr1 | 12/09/2025 | 7 | 100 | 10000 | 142 | 0 | 142 | 2.84E+06 | 0 | 0 |
| LRH316 | Chr1 | 12/09/2025 | 8 | 100 | 10000 | 67 | 0 | 67 | 1.34E+06 | 0 | 0 |
| LRH316 | Chr1 | 12/09/2025 | 9 | 100 | 10000 | 137 | 0 | 137 | 2.74E+06 | 0 | 0 |
| LRH316 | Chr1 | 12/09/2025 | 10 | 100 | 10000 | 90 | 0 | 90 | 1.80E+06 | 0 | 0 |
| LRH316 | Chr1 | 12/09/2025 | 11 | 100 | 10000 | 138 | 0 | 138 | 2.76E+06 | 0 | 0 |
| LRH316 | Chr1 | 12/09/2025 | 12 | 100 | 10000 | 90 | 1 | 91 | 1.82E+06 | 0.0109890109 | 1.098901099 |
| LRH316 | Chr1 | 12/09/2025 | 13 | 100 | 10000 | 67 | 0 | 67 | 1.34E+06 | 0 | 0 |
| LRH316 | Chr1 | 12/09/2025 | 14 | 100 | 10000 | 81 | 0 | 81 | 1.62E+06 | 0 | 0 |
| LRH316 | Chr1 | 12/09/2025 | 15 | 100 | 10000 | 102 | 0 | 102 | 2.04E+06 | 0 | 0 |
| LRH316 | Chr1 | 12/09/2025 | 16 | 100 | 10000 | 125 | 0 | 125 | 2.50E+06 | 0 | 0 |
| LRH316 | Chr1 | 12/09/2025 | 17 | 100 | 10000 | 104 | 0 | 104 | 2.08E+06 | 0 | 0 |
| LRH316 | Chr1 | 12/09/2025 | 18 | 100 | 10000 | 90 | 0 | 90 | 1.80E+06 | 0 | 0 |
| LRH316 | Chr1 | 12/09/2025 | 19 | 100 | 10000 | 84 | 0 | 84 | 1.68E+06 | 0 | 0 |
| LRH316 | Chr1 | 12/09/2025 | 20 | 100 | 10000 | 157 | 0 | 157 | 3.14E+06 | 0 | 0 |
| LRH316 | Chr1 | 12/09/2025 | 21 | 100 | 10000 | 100 | 0 | 100 | 2.00E+06 | 0 | 0 |
| LRH316 | Chr1 | 12/09/2025 | 22 | 100 | 10000 | 93 | 1 | 94 | 1.88E+06 | 0.01063829787 | 1.063829787 |
| LRH316 | Chr1 | 12/09/2025 | 23 | 100 | 10000 | 113 | 0 | 113 | 2.26E+06 | 0 | 0 |
| LRH316 | Chr1 | 12/09/2025 | 24 | 100 | 10000 | 113 | 0 | 113 | 2.26E+06 | 0 | 0 |
| LRH587 | Chr2 | 12/10/2025 | 1 | 100 | 10000 | 230 | 0 | 230 | 4.60E+06 | 0 | 0 |
| LRH587 | Chr2 | 12/10/2025 | 2 | 100 | 10000 | 46 | 0 | 46 | 9.20E+05 | 0 | 0 |
| LRH587 | Chr2 | 12/10/2025 | 3 | 100 | 10000 | 32 | 0 | 32 | 6.40E+05 | 0 | 0 |
| LRH587 | Chr2 | 12/10/2025 | 4 | 100 | 10000 | 56 | 0 | 56 | 1.12E+06 | 0 | 0 |
| LRH587 | Chr2 | 12/10/2025 | 5 | 100 | 10000 | 54 | 0 | 54 | 1.08E+06 | 0 | 0 |
| LRH587 | Chr2 | 12/10/2025 | 6 | 100 | 10000 | 31 | 0 | 31 | 6.20E+05 | 0 | 0 |
| LRH587 | Chr2 | 12/10/2025 | 7 | 100 | 10000 | 47 | 0 | 47 | 9.40E+05 | 0 | 0 |
| LRH587 | Chr2 | 12/10/2025 | 8 | 100 | 10000 | 52 | 0 | 52 | 1.04E+06 | 0 | 0 |
| LRH587 | Chr2 | 12/10/2025 | 9 | 100 | 10000 | 47 | 0 | 47 | 9.40E+05 | 0 | 0 |
| LRH587 | Chr2 | 12/10/2025 | 10 | 100 | 10000 | 89 | 0 | 89 | 1.78E+06 | 0 | 0 |
| LRH587 | Chr2 | 12/10/2025 | 11 | 100 | 10000 | 79 | 0 | 79 | 1.58E+06 | 0 | 0 |
| LRH587 | Chr2 | 12/10/2025 | 12 | 100 | 10000 | 76 | 0 | 76 | 1.52E+06 | 0 | 0 |
| LRH587 | Chr2 | 12/10/2025 | 13 | 100 | 10000 | 68 | 0 | 68 | 1.36E+06 | 0 | 0 |
| LRH587 | Chr2 | 12/10/2025 | 14 | 100 | 10000 | 46 | 0 | 46 | 9.20E+05 | 0 | 0 |
| LRH587 | Chr2 | 12/10/2025 | 15 | 100 | 10000 | 62 | 1 | 63 | 1.26E+06 | 0.01587301587 | 1.587301587 |
| LRH587 | Chr2 | 12/10/2025 | 16 | 100 | 10000 | 59 | 0 | 59 | 1.18E+06 | 0 | 0 |
| LRH587 | Chr2 | 12/10/2025 | 17 | 100 | 10000 | 63 | 0 | 63 | 1.26E+06 | 0 | 0 |
| LRH587 | Chr2 | 12/10/2025 | 18 | 100 | 10000 | 39 | 0 | 39 | 7.80E+05 | 0 | 0 |
| LRH587 | Chr2 | 12/10/2025 | 19 | 100 | 10000 | 32 | 0 | 32 | 6.40E+05 | 0 | 0 |
| LRH587 | Chr2 | 12/10/2025 | 20 | 100 | 10000 | 47 | 0 | 47 | 9.40E+05 | 0 | 0 |
| LRH587 | Chr2 | 12/10/2025 | 21 | 100 | 10000 | 76 | 0 | 76 | 1.52E+06 | 0 | 0 |
| LRH587 | Chr2 | 12/10/2025 | 22 | 100 | 10000 | 45 | 0 | 45 | 9.00E+05 | 0 | 0 |
| LRH587 | Chr2 | 12/10/2025 | 23 | 100 | 10000 | 60 | 0 | 60 | 1.20E+06 | 0 | 0 |
| LRH587 | Chr2 | 12/10/2025 | 24 | 100 | 10000 | 84 | 0 | 84 | 1.68E+06 | 0 | 0 |
| LRH575 | Chr3 | 12/09/2025 | 1 | 100 | 10000 | 264 | 0 | 264 | 5.28E+06 | 0 | 0 |
| LRH575 | Chr3 | 12/09/2025 | 2 | 100 | 10000 | 432 | 0 | 432 | 8.64E+06 | 0 | 0 |
| LRH575 | Chr3 | 12/09/2025 | 3 | 100 | 10000 | 354 | 0 | 354 | 7.08E+06 | 0 | 0 |
| LRH575 | Chr3 | 12/09/2025 | 4 | 100 | 10000 | 364 | 2 | 366 | 7.32E+06 | 0.00546448087 | 0.5464480874 |
| LRH575 | Chr3 | 12/09/2025 | 5 | 100 | 10000 | 276 | 0 | 276 | 5.52E+06 | 0 | 0 |
| LRH575 | Chr3 | 12/09/2025 | 6 | 100 | 10000 | 494 | 0 | 494 | 9.88E+06 | 0 | 0 |
| LRH575 | Chr3 | 12/09/2025 | 7 | 100 | 10000 | 350 | 0 | 350 | 7.00E+06 | 0 | 0 |
| LRH575 | Chr3 | 12/09/2025 | 8 | 100 | 10000 | 566 | 0 | 566 | 1.13E+07 | 0 | 0 |
| LRH575 | Chr3 | 12/09/2025 | 9 | 100 | 10000 | 204 | 0 | 204 | 4.08E+06 | 0 | 0 |
| LRH575 | Chr3 | 12/09/2025 | 10 | 100 | 10000 | 558 | 2 | 560 | 1.12E+07 | 0.00357142857 | 0.3571428571 |
| LRH575 | Chr3 | 12/09/2025 | 11 | 100 | 10000 | 776 | 0 | 776 | 1.55E+07 | 0 | 0 |
| LRH575 | Chr3 | 12/09/2025 | 12 | 100 | 10000 | 474 | 0 | 474 | 9.48E+06 | 0 | 0 |
| LRH575 | Chr3 | 12/09/2025 | 13 | 100 | 10000 | 48 | 0 | 48 | 9.60E+05 | 0 | 0 |
| LRH575 | Chr3 | 12/09/2025 | 14 | 100 | 10000 | 110 | 0 | 110 | 2.20E+06 | 0 | 0 |
| LRH575 | Chr3 | 12/09/2025 | 15 | 100 | 10000 | 205 | 0 | 205 | 4.10E+06 | 0 | 0 |
| LRH575 | Chr3 | 12/09/2025 | 16 | 100 | 10000 | 187 | 0 | 187 | 3.74E+06 | 0 | 0 |
| LRH575 | Chr3 | 12/09/2025 | 17 | 100 | 10000 | 199 | 0 | 199 | 3.98E+06 | 0 | 0 |
| LRH575 | Chr3 | 12/09/2025 | 18 | 100 | 10000 | 184 | 0 | 184 | 3.68E+06 | 0 | 0 |
| LRH575 | Chr3 | 12/09/2025 | 19 | 100 | 10000 | 222 | 0 | 222 | 4.44E+06 | 0 | 0 |
| LRH575 | Chr3 | 12/09/2025 | 20 | 100 | 10000 | 314 | 0 | 314 | 6.28E+06 | 0 | 0 |
| LRH575 | Chr3 | 12/09/2025 | 21 | 100 | 10000 | 448 | 0 | 448 | 8.96E+06 | 0 | 0 |
| LRH575 | Chr3 | 12/09/2025 | 22 | 100 | 10000 | 195 | 0 | 195 | 3.90E+06 | 0 | 0 |
| LRH575 | Chr3 | 12/09/2025 | 23 | 100 | 10000 | 318 | 0 | 318 | 6.36E+06 | 0 | 0 |
| LRH575 | Chr3 | 12/09/2025 | 24 | 100 | 10000 | 462 | 0 | 462 | 9.24E+06 | 0 | 0 |
| LRH562 | Chr4 | 12/10/2025 | 1 | 100 | 1000 | 115 | 12 | 127 | 2.54E+05 | 0.0944881889 | 9.44881889 |
| LRH562 | Chr4 | 12/10/2025 | 2 | 100 | 1000 | 117 | 0 | 117 | 2.34E+05 | 0 | 0 |
| LRH562 | Chr4 | 12/10/2025 | 3 | 100 | 1000 | 186 | 3 | 189 | 3.78E+05 | 0.0158730158 | 1.587301587 |
| LRH562 | Chr4 | 12/10/2025 | 4 | 100 | 1000 | 274 | 3 | 277 | 5.54E+05 | 0.0108303249 | 1.083032491 |
| LRH562 | Chr4 | 12/10/2025 | 5 | 100 | 1000 | 158 | 2 | 160 | 3.20E+05 | 0.0125 | 1.25 |
| LRH562 | Chr4 | 12/10/2025 | 6 | 100 | 1000 | 160 | 23 | 183 | 3.66E+05 | 0.1256830601 | 12.56830601 |
| LRH562 | Chr4 | 12/10/2025 | 7 | 100 | 1000 | 92 | 4 | 96 | 1.92E+05 | 0.0416666667 | 4.16666667 |
| LRH562 | Chr4 | 12/10/2025 | 8 | 100 | 1000 | 100 | 8 | 108 | 2.16E+05 | 0.0740740740 | 7.407407407 |
| LRH562 | Chr4 | 12/10/2025 | 9 | 100 | 1000 | 116 | 7 | 123 | 2.46E+05 | 0.0569105691 | 5.691056911 |
| LRH562 | Chr4 | 12/10/2025 | 10 | 100 | 1000 | 196 | 3 | 199 | 3.98E+05 | 0.0150753768 | 1.507537688 |
| LRH562 | Chr4 | 12/10/2025 | 12 | 100 | 1000 | 160 | 4 | 164 | 3.28E+05 | 0.0243902439 | 2.43902439 |
| LRH562 | Chr4 | 12/10/2025 | 13 | 100 | 1000 | 106 | 3 | 109 | 2.18E+05 | 0.0275229357 | 2.752293578 |
| LRH562 | Chr4 | 12/10/2025 | 14 | 100 | 1000 | 54 | 92 | 146 | 2.92E+05 | 0.6301369863 | 63.01369863 |
| LRH562 | Chr4 | 12/10/2025 | 15 | 100 | 1000 | 100 | 0 | 100 | 2.00E+05 | 0 | 0 |
| LRH562 | Chr4 | 12/10/2025 | 16 | 100 | 1000 | 84 | 2 | 86 | 1.72E+05 | 0.0232558139 | 2.325581395 |
| LRH562 | Chr4 | 12/10/2025 | 17 | 100 | 1000 | 20 | 716 | 736 | 1.47E+06 | 0.972826087 | 97.2826087 |
| LRH562 | Chr4 | 12/10/2025 | 18 | 100 | 1000 | 106 | 0 | 106 | 2.12E+05 | 0 | 0 |
| LRH562 | Chr4 | 12/10/2025 | 19 | 100 | 1000 | 142 | 3 | 145 | 2.90E+05 | 0.0206896551 | 2.068965517 |
| LRH562 | Chr4 | 12/10/2025 | 20 | 100 | 1000 | 88 | 3 | 91 | 1.82E+05 | 0.0329670329 | 3.296703297 |
| LRH562 | Chr4 | 12/10/2025 | 21 | 100 | 1000 | 152 | 2 | 154 | 3.08E+05 | 0.0129870129 | 1.298701299 |
| LRH562 | Chr4 | 12/10/2025 | 22 | 100 | 1000 | 10 | 236 | 246 | 4.92E+05 | 0.9593495935 | 95.93495935 |
| LRH562 | Chr4 | 12/10/2025 | 23 | 100 | 1000 | 74 | 15 | 89 | 1.78E+05 | 0.1685393258 | 16.85393258 |
| LRH562 | Chr4 | 12/10/2025 | 24 | 100 | 1000 | 92 | 0 | 92 | 1.84E+05 | 0 | 0 |
| LRH348 | Chr5 | 12/09/2025 | 1 | 100 | 10000 | 48 | 1 | 49 | 9.80E+05 | 0.0204081632 | 2.040816327 |
| LRH348 | Chr5 | 12/09/2025 | 2 | 100 | 10000 | 29 | 0 | 29 | 5.80E+05 | 0 | 0 |
| LRH348 | Chr5 | 12/09/2025 | 3 | 100 | 10000 | 33 | 0 | 33 | 6.60E+05 | 0 | 0 |
| LRH348 | Chr5 | 12/09/2025 | 4 | 100 | 10000 | 33 | 0 | 33 | 6.60E+05 | 0 | 0 |
| LRH348 | Chr5 | 12/09/2025 | 5 | 100 | 10000 | 50 | 0 | 50 | 1.00E+06 | 0 | 0 |
| LRH348 | Chr5 | 12/09/2025 | 6 | 100 | 10000 | 43 | 1 | 44 | 8.80E+05 | 0.0227272727 | 2.272727273 |
| LRH348 | Chr5 | 12/09/2025 | 7 | 100 | 10000 | 52 | 0 | 52 | 1.04E+06 | 0 | 0 |
| LRH348 | Chr5 | 12/09/2025 | 8 | 100 | 10000 | 50 | 0 | 50 | 1.00E+06 | 0 | 0 |
| LRH348 | Chr5 | 12/09/2025 | 9 | 100 | 10000 | 47 | 0 | 47 | 9.40E+05 | 0 | 0 |
| LRH348 | Chr5 | 12/09/2025 | 10 | 100 | 10000 | 55 | 0 | 55 | 1.10E+06 | 0 | 0 |
| LRH348 | Chr5 | 12/09/2025 | 11 | 100 | 10000 | 59 | 0 | 59 | 1.18E+06 | 0 | 0 |
| LRH348 | Chr5 | 12/09/2025 | 12 | 100 | 10000 | 62 | 0 | 62 | 1.24E+06 | 0 | 0 |
| LRH348 | Chr5 | 12/09/2025 | 13 | 100 | 10000 | 39 | 0 | 39 | 7.80E+05 | 0 | 0 |
| LRH348 | Chr5 | 12/09/2025 | 14 | 100 | 10000 | 31 | 1 | 32 | 6.40E+05 | 0.03125 | 3.125 |
| LRH348 | Chr5 | 12/09/2025 | 15 | 100 | 10000 | 32 | 0 | 32 | 6.40E+05 | 0 | 0 |
| LRH348 | Chr5 | 12/09/2025 | 16 | 100 | 10000 | 37 | 0 | 37 | 7.40E+05 | 0 | 0 |
| LRH348 | Chr5 | 12/09/2025 | 17 | 100 | 10000 | 43 | 0 | 43 | 8.60E+05 | 0 | 0 |
| LRH348 | Chr5 | 12/09/2025 | 18 | 100 | 10000 | 44 | 0 | 44 | 8.80E+05 | 0 | 0 |
| LRH348 | Chr5 | 12/09/2025 | 19 | 100 | 10000 | 27 | 0 | 27 | 5.40E+05 | 0 | 0 |
| LRH348 | Chr5 | 12/09/2025 | 20 | 100 | 10000 | 43 | 0 | 43 | 8.60E+05 | 0 | 0 |
| LRH348 | Chr5 | 12/09/2025 | 21 | 100 | 10000 | 28 | 0 | 28 | 5.60E+05 | 0 | 0 |
| LRH348 | Chr5 | 12/09/2025 | 22 | 100 | 10000 | 45 | 0 | 45 | 9.00E+05 | 0 | 0 |
| LRH348 | Chr5 | 12/09/2025 | 23 | 100 | 10000 | 37 | 0 | 37 | 7.40E+05 | 0 | 0 |
| LRH348 | Chr5 | 12/09/2025 | 24 | 100 | 10000 | 35 | 0 | 35 | 7.00E+05 | 0 | 0 |
| LRH510 | Chr8 | 12/10/2025 | 1 | 100 | 10000 | 248 | 0 | 248 | 4.96E+06 | 0 | 0 |
| LRH510 | Chr8 | 12/10/2025 | 2 | 100 | 10000 | 316 | 0 | 316 | 6.32E+06 | 0 | 0 |
| LRH510 | Chr8 | 12/10/2025 | 3 | 100 | 10000 | 328 | 0 | 328 | 6.56E+06 | 0 | 0 |
| LRH510 | Chr8 | 12/10/2025 | 4 | 100 | 10000 | 378 | 0 | 378 | 7.56E+06 | 0 | 0 |
| LRH510 | Chr8 | 12/10/2025 | 5 | 100 | 10000 | 196 | 0 | 196 | 3.92E+06 | 0 | 0 |
| LRH510 | Chr8 | 12/10/2025 | 6 | 100 | 10000 | 356 | 0 | 356 | 7.12E+06 | 0 | 0 |
| LRH510 | Chr8 | 12/10/2025 | 7 | 100 | 10000 | 230 | 0 | 230 | 4.60E+06 | 0 | 0 |
| LRH510 | Chr8 | 12/10/2025 | 8 | 100 | 10000 | 302 | 0 | 302 | 6.04E+06 | 0 | 0 |
| LRH510 | Chr8 | 12/10/2025 | 9 | 100 | 10000 | 141 | 0 | 141 | 2.82E+06 | 0 | 0 |
| LRH510 | Chr8 | 12/10/2025 | 10 | 100 | 10000 | 218 | 0 | 218 | 4.36E+06 | 0 | 0 |
| LRH510 | Chr8 | 12/10/2025 | 11 | 100 | 10000 | 202 | 0 | 202 | 4.04E+06 | 0 | 0 |
| LRH510 | Chr8 | 12/10/2025 | 12 | 100 | 10000 | 142 | 0 | 142 | 2.84E+06 | 0 | 0 |
| LRH510 | Chr8 | 12/10/2025 | 13 | 100 | 10000 | 164 | 1 | 165 | 3.30E+06 | 0.0060606060 | 0.6060606061 |
| LRH510 | Chr8 | 12/10/2025 | 14 | 100 | 10000 | 189 | 0 | 189 | 3.78E+06 | 0 | 0 |
| LRH510 | Chr8 | 12/10/2025 | 15 | 100 | 10000 | 232 | 0 | 232 | 4.64E+06 | 0 | 0 |
| LRH510 | Chr8 | 12/10/2025 | 16 | 100 | 10000 | 166 | 0 | 166 | 3.32E+06 | 0 | 0 |
| LRH510 | Chr8 | 12/10/2025 | 17 | 100 | 10000 | 166 | 0 | 166 | 3.32E+06 | 0 | 0 |
| LRH510 | Chr8 | 12/10/2025 | 18 | 100 | 10000 | 150 | 0 | 150 | 3.00E+06 | 0 | 0 |
| LRH510 | Chr8 | 12/10/2025 | 19 | 100 | 10000 | 182 | 0 | 182 | 3.64E+06 | 0 | 0 |
| LRH510 | Chr8 | 12/10/2025 | 20 | 100 | 10000 | 75 | 0 | 75 | 1.50E+06 | 0 | 0 |
| LRH510 | Chr8 | 12/10/2025 | 21 | 100 | 10000 | 314 | 0 | 314 | 6.28E+06 | 0 | 0 |
| LRH510 | Chr8 | 12/10/2025 | 22 | 100 | 10000 | 410 | 0 | 410 | 8.20E+06 | 0 | 0 |
| LRH510 | Chr8 | 12/10/2025 | 23 | 100 | 10000 | 302 | 0 | 302 | 6.04E+06 | 0 | 0 |
| LRH510 | Chr8 | 12/10/2025 | 24 | 100 | 10000 | 402 | 0 | 402 | 8.04E+06 | 0 | 0 |
| LRH433 | Chr9 | 12/09/2025 | 1 | 100 | 10000 | 37 | 0 | 37 | 7.40E+05 | 0 | 0 |
| LRH433 | Chr9 | 12/09/2025 | 2 | 100 | 10000 | 51 | 0 | 51 | 1.02E+06 | 0 | 0 |
| LRH433 | Chr9 | 12/09/2025 | 3 | 100 | 10000 | 28 | 0 | 28 | 5.60E+05 | 0 | 0 |
| LRH433 | Chr9 | 12/09/2025 | 4 | 100 | 10000 | 88 | 0 | 88 | 1.76E+06 | 0 | 0 |
| LRH433 | Chr9 | 12/09/2025 | 5 | 100 | 10000 | 74 | 0 | 74 | 1.48E+06 | 0 | 0 |
| LRH433 | Chr9 | 12/09/2025 | 6 | 100 | 10000 | 46 | 0 | 46 | 9.20E+05 | 0 | 0 |
| LRH433 | Chr9 | 12/09/2025 | 7 | 100 | 10000 | 42 | 0 | 42 | 8.40E+05 | 0 | 0 |
| LRH433 | Chr9 | 12/09/2025 | 8 | 100 | 10000 | 52 | 0 | 52 | 1.04E+06 | 0 | 0 |
| LRH433 | Chr9 | 12/09/2025 | 9 | 100 | 10000 | 62 | 0 | 62 | 1.24E+06 | 0 | 0 |
| LRH433 | Chr9 | 12/09/2025 | 10 | 100 | 10000 | 58 | 0 | 58 | 1.16E+06 | 0 | 0 |
| LRH433 | Chr9 | 12/09/2025 | 11 | 100 | 10000 | 59 | 0 | 59 | 1.18E+06 | 0 | 0 |
| LRH433 | Chr9 | 12/09/2025 | 12 | 100 | 10000 | 52 | 0 | 52 | 1.04E+06 | 0 | 0 |
| LRH433 | Chr9 | 12/09/2025 | 13 | 100 | 10000 | 59 | 0 | 59 | 1.18E+06 | 0 | 0 |
| LRH433 | Chr9 | 12/09/2025 | 14 | 100 | 10000 | 38 | 0 | 38 | 7.60E+05 | 0 | 0 |
| LRH433 | Chr9 | 12/09/2025 | 15 | 100 | 10000 | 71 | 0 | 71 | 1.42E+06 | 0 | 0 |
| LRH433 | Chr9 | 12/09/2025 | 16 | 100 | 10000 | 51 | 0 | 51 | 1.02E+06 | 0 | 0 |
| LRH433 | Chr9 | 12/09/2025 | 17 | 100 | 10000 | 44 | 4 | 48 | 9.60E+05 | 0.0833333333333333 | 8.333333333 |
| LRH433 | Chr9 | 12/09/2025 | 18 | 100 | 10000 | 54 | 0 | 54 | 1.08E+06 | 0 | 0 |
| LRH433 | Chr9 | 12/09/2025 | 19 | 100 | 10000 | 50 | 0 | 50 | 1.00E+06 | 0 | 0 |
| LRH433 | Chr9 | 12/09/2025 | 20 | 100 | 10000 | 66 | 0 | 66 | 1.32E+06 | 0 | 0 |
| LRH433 | Chr9 | 12/09/2025 | 21 | 100 | 10000 | 98 | 1 | 99 | 1.98E+06 | 0.0101010101 | 1.01010101 |
| LRH433 | Chr9 | 12/09/2025 | 22 | 100 | 10000 | 54 | 0 | 54 | 1.08E+06 | 0 | 0 |
| LRH433 | Chr9 | 12/09/2025 | 23 | 100 | 10000 | 67 | 0 | 67 | 1.34E+06 | 0 | 0 |
| LRH433 | Chr9 | 12/09/2025 | 24 | 100 | 10000 | 70 | 0 | 70 | 1.40E+06 | 0 | 0 |
| LRH502 | Chr10 | 12/10/2025 | 1 | 100 | 10000 | 312 | 6 | 318 | 6.36E+06 | 0.01886792453 | 1.886792453 |
| LRH502 | Chr10 | 12/10/2025 | 2 | 100 | 10000 | 158 | 4 | 162 | 3.24E+06 | 0.02469135802 | 2.469135802 |
| LRH502 | Chr10 | 12/10/2025 | 3 | 100 | 10000 | 47 | 0 | 47 | 9.40E+05 | 0 | 0 |
| LRH502 | Chr10 | 12/10/2025 | 4 | 100 | 10000 | 208 | 8 | 216 | 4.32E+06 | 0.0370370370370370 | 3.703703704 |
| LRH502 | Chr10 | 12/10/2025 | 5 | 100 | 10000 | 51 | 1 | 52 | 1.04E+06 | 0.0192307692307692 | 1.923076923 |
| LRH502 | Chr10 | 12/10/2025 | 6 | 100 | 10000 | 171 | 3 | 174 | 3.48E+06 | 0.0172413793103448 | 1.724137931 |
| LRH502 | Chr10 | 12/10/2025 | 7 | 100 | 10000 | 207 | 1 | 208 | 4.16E+06 | 0.004807692307692308 | 0.4807692308 |
| LRH502 | Chr10 | 12/10/2025 | 8 | 100 | 10000 | 210 | 4 | 214 | 4.28E+06 | 0.0186915887943972 | 1.869158879 |
| LRH502 | Chr10 | 12/10/2025 | 9 | 100 | 10000 | 92 | 0 | 92 | 1.84E+06 | 0 | 0 |
| LRH502 | Chr10 | 12/10/2025 | 10 | 100 | 10000 | 122 | 2 | 124 | 2.48E+06 | 0.0161290322580645 | 1.612903226 |
| LRH502 | Chr10 | 12/10/2025 | 11 | 100 | 10000 | 67 | 2 | 69 | 1.38E+06 | 0.02898550725190496 | 2.898550725 |
| LRH502 | Chr10 | 12/10/2025 | 12 | 100 | 10000 | 283 | 3 | 286 | 5.72E+06 | 0.0104895104545455 | 1.048951049 |
| LRH502 | Chr10 | 12/10/2025 | 13 | 100 | 10000 | 123 | 0 | 123 | 2.46E+06 | 0 | 0 |
| LRH502 | Chr10 | 12/10/2025 | 14 | 100 | 10000 | 106 | 1 | 107 | 2.14E+06 | 0.0093457943972528 | 0.9345794393 |
| LRH502 | Chr10 | 12/10/2025 | 15 | 100 | 10000 | 142 | 0 | 142 | 2.84E+06 | 0 | 0 |
| LRH502 | Chr10 | 12/10/2025 | 16 | 100 | 10000 | 13 | 0 | 13 | 2.60E+05 | 0 | 0 |
| LRH502 | Chr10 | 12/10/2025 | 17 | 100 | 10000 | 141 | 1 | 142 | 2.84E+06 | 0.00704225352112676 | 0.7042253521 |
| LRH502 | Chr10 | 12/10/2025 | 18 | 100 | 10000 | 139 | 7 | 146 | 2.92E+06 | 0.0479452054863014 | 4.794520548 |
| LRH502 | Chr10 | 12/10/2025 | 19 | 100 | 10000 | 122 | 0 | 122 | 2.44E+06 | 0 | 0 |
| LRH502 | Chr10 | 12/10/2025 | 20 | 100 | 10000 | 131 | 1 | 132 | 2.64E+06 | 0.0075757575757576 | 0.7575757576 |
| LRH502 | Chr10 | 12/10/2025 | 21 | 100 | 10000 | 108 | 0 | 108 | 2.16E+06 | 0 | 0 |
| LRH502 | Chr10 | 12/10/2025 | 22 | 100 | 10000 | 103 | 18 | 121 | 2.42E+06 | 0.1487603306528926 | 14.87603306 |
| LRH502 | Chr10 | 12/10/2025 | 23 | 100 | 10000 | 144 | 0 | 144 | 2.88E+06 | 0 | 0 |
| LRH502 | Chr10 | 12/10/2025 | 24 | 100 | 10000 | 103 | 8 | 111 | 2.22E+06 | 0.0720720720720721 | 7.207207207 |
| LRH566 | Chr11 | 12/10/2025 | 1 | 100 | 10000 | 182 | 0 | 182 | 3.64E+06 | 0 | 0 |
| LRH566 | Chr11 | 12/10/2025 | 2 | 100 | 10000 | 166 | 0 | 166 | 3.32E+06 | 0 | 0 |
| LRH566 | Chr11 | 12/10/2025 | 3 | 100 | 10000 | 254 | 0 | 254 | 5.08E+06 | 0 | 0 |
| LRH566 | Chr11 | 12/10/2025 | 4 | 100 | 10000 | 164 | 0 | 164 | 3.28E+06 | 0 | 0 |
| LRH566 | Chr11 | 12/10/2025 | 5 | 100 | 10000 | 216 | 0 | 216 | 4.32E+06 | 0 | 0 |
| LRH566 | Chr11 | 12/10/2025 | 6 | 100 | 10000 | 198 | 0 | 198 | 3.96E+06 | 0 | 0 |
| LRH566 | Chr11 | 12/10/2025 | 7 | 100 | 10000 | 293 | 1 | 294 | 5.88E+06 | 0.0034013605442177 | 0.3401360544 |
| LRH566 | Chr11 | 12/10/2025 | 8 | 100 | 10000 | 207 | 1 | 208 | 4.16E+06 | 0.004807692307692308 | 0.4807692308 |
| LRH566 | Chr11 | 12/10/2025 | 9 | 100 | 10000 | 255 | 1 | 256 | 5.12E+06 | 0.00390625 | 0.390625 |
| LRH566 | Chr11 | 12/10/2025 | 10 | 100 | 10000 | 206 | 0 | 206 | 4.12E+06 | 0 | 0 |
| LRH566 | Chr11 | 12/10/2025 | 11 | 100 | 10000 | 291 | 1 | 292 | 5.84E+06 | 0.0034246575342466 | 0.3424657534 |
| LRH566 | Chr11 | 12/10/2025 | 12 | 100 | 10000 | 238 | 0 | 238 | 4.76E+06 | 0 | 0 |
| LRH566 | Chr11 | 12/10/2025 | 13 | 100 | 10000 | 122 | 0 | 122 | 2.44E+06 | 0 | 0 |
| LRH566 | Chr11 | 12/10/2025 | 14 | 100 | 10000 | 187 | 1 | 188 | 3.76E+06 | 0.0053191489361702 | 0.5319148936 |
| LRH566 | Chr11 | 12/10/2025 | 15 | 100 | 10000 | 235 | 1 | 236 | 4.72E+06 | 0.0042372881156779 | 0.4237288136 |
| LRH566 | Chr11 | 12/10/2025 | 16 | 100 | 10000 | 148 | 0 | 148 | 2.96E+06 | 0 | 0 |
| LRH566 | Chr11 | 12/10/2025 | 17 | 100 | 10000 | 252 | 0 | 252 | 5.04E+06 | 0 | 0 |
| LRH566 | Chr11 | 12/10/2025 | 18 | 100 | 10000 | 112 | 0 | 112 | 2.24E+06 | 0 | 0 |
| LRH566 | Chr11 | 12/10/2025 | 19 | 100 | 10000 | 190 | 0 | 190 | 3.80E+06 | 0 | 0 |
| LRH566 | Chr11 | 12/10/2025 | 20 | 100 | 10000 | 83 | 0 | 83 | 1.66E+06 | 0 | 0 |
| LRH566 | Chr11 | 12/10/2025 | 21 | 100 | 10000 | 250 | 0 | 250 | 5.00E+06 | 0 | 0 |
| LRH566 | Chr11 | 12/10/2025 | 22 | 100 | 10000 | 87 | 0 | 87 | 1.74E+06 | 0 | 0 |
| LRH566 | Chr11 | 12/10/2025 | 23 | 100 | 10000 | 186 | 0 | 186 | 3.72E+06 | 0 | 0 |
| LRH566 | Chr11 | 12/10/2025 | 24 | 100 | 10000 | 244 | 0 | 244 | 4.88E+06 | 0 | 0 |
| LRH488 | Chr12 | 12/10/2025 | 1 | 100 | 10000 | 0 | 148 | 148 | 2.96E+06 | 1 | 100 |
| LRH488 | Chr12 | 12/10/2025 | 2 | 100 | 10000 | 6 | 190 | 196 | 3.92E+06 | 0.9693877551 | 96.93877551 |
| LRH488 | Chr12 | 12/10/2025 | 3 | 100 | 10000 | 4 | 148 | 152 | 3.04E+06 | 0.9736842105 | 97.36842105 |
| LRH488 | Chr12 | 12/10/2025 | 4 | 100 | 10000 | 1 | 298 | 299 | 5.98E+06 | 0.9966555184 | 99.66555184 |
| LRH488 | Chr12 | 12/10/2025 | 5 | 100 | 10000 | 1 | 100 | 101 | 2.02E+06 | 0.9900990099 | 99.00990099 |
| LRH488 | Chr12 | 12/10/2025 | 6 | 100 | 10000 | 1 | 110 | 111 | 2.22E+06 | 0.990990991 | 99.0990991 |
| LRH488 | Chr12 | 12/10/2025 | 7 | 100 | 10000 | 0 | 146 | 146 | 2.92E+06 | 1 | 100 |
| LRH488 | Chr12 | 12/10/2025 | 8 | 100 | 10000 | 22 | 168 | 190 | 3.80E+06 | 0.8842105263 | 88.42105263 |
| LRH488 | Chr12 | 12/10/2025 | 9 | 100 | 10000 | 2 | 132 | 134 | 2.68E+06 | 0.9850746269 | 98.50746269 |
| LRH488 | Chr12 | 12/10/2025 | 10 | 100 | 10000 | 4 | 188 | 192 | 3.84E+06 | 0.9791666667 | 97.91666667 |
| LRH488 | Chr12 | 12/10/2025 | 11 | 100 | 10000 | 1 | 232 | 233 | 4.66E+06 | 0.9957081545 | 99.57081545 |
| LRH488 | Chr12 | 12/10/2025 | 12 | 100 | 10000 | 4 | 184 | 188 | 3.76E+06 | 0.9787234043 | 97.87234043 |
| LRH488 | Chr12 | 12/10/2025 | 13 | 100 | 10000 | 1 | 142 | 143 | 2.86E+06 | 0.993006993 | 99.3006993 |
| LRH488 | Chr12 | 12/10/2025 | 14 | 100 | 10000 | 0 | 138 | 138 | 2.76E+06 | 1 | 100 |
| LRH488 | Chr12 | 12/10/2025 | 15 | 100 | 10000 | 4 | 166 | 170 | 3.40E+06 | 0.9764705882 | 97.64705882 |
| LRH488 | Chr12 | 12/10/2025 | 16 | 100 | 10000 | 0 | 89 | 89 | 1.78E+06 | 1 | 100 |
| LRH488 | Chr12 | 12/10/2025 | 17 | 100 | 10000 | 2 | 117 | 119 | 2.38E+06 | 0.9831932773 | 98.31932773 |
| LRH488 | Chr12 | 12/10/2025 | 18 | 100 | 10000 | 1 | 150 | 151 | 3.02E+06 | 0.9933774834 | 99.33774834 |
| LRH488 | Chr12 | 12/10/2025 | 19 | 100 | 10000 | 0 | 128 | 128 | 2.56E+06 | 1 | 100 |
| LRH488 | Chr12 | 12/10/2025 | 20 | 100 | 10000 | 5 | 146 | 151 | 3.02E+06 | 0.9668874172 | 96.68874172 |
| LRH488 | Chr12 | 12/10/2025 | 21 | 100 | 10000 | 104 | 34 | 138 | 2.76E+06 | 0.2463768116 | 24.63768116 |
| LRH488 | Chr12 | 12/10/2025 | 22 | 100 | 10000 | 0 | 83 | 83 | 1.66E+06 | 1 | 100 |
| LRH488 | Chr12 | 12/10/2025 | 23 | 100 | 10000 | 2 | 278 | 280 | 5.60E+06 | 0.9928571429 | 99.28571429 |
| LRH488 | Chr12 | 12/10/2025 | 24 | 100 | 10000 | 6 | 288 | 294 | 5.88E+06 | 0.9795918367 | 97.95918367 |
| LRH570 | Chr13 | 12/10/2025 | 1 | 100 | 10000 | 36 | 2 | 38 | 7.60E+05 | 0.0526315789 | 5.26315789 |
| LRH570 | Chr13 | 12/10/2025 | 2 | 100 | 10000 | 25 | 2 | 27 | 5.40E+05 | 0.0740740740 | 7.40740740 |
| LRH570 | Chr13 | 12/10/2025 | 3 | 100 | 10000 | 13 | 0 | 13 | 2.60E+05 | 0 | 0 |
| LRH570 | Chr13 | 12/10/2025 | 4 | 100 | 10000 | 9 | 67 | 76 | 1.52E+06 | 0.8815789474 | 88.15789474 |
| LRH570 | Chr13 | 12/10/2025 | 5 | 100 | 10000 | 26 | 41 | 67 | 1.34E+06 | 0.6119402985 | 61.19402985 |
| LRH570 | Chr13 | 12/10/2025 | 6 | 100 | 10000 | 5 | 0 | 5 | 1.00E+05 | 0 | 0 |
| LRH570 | Chr13 | 12/10/2025 | 7 | 100 | 10000 | 19 | 5 | 24 | 4.80E+05 | 0.2083333333 | 20.83333333 |
| LRH570 | Chr13 | 12/10/2025 | 8 | 100 | 10000 | 25 | 4 | 29 | 5.80E+05 | 0.1379310345 | 13.79310345 |
| LRH570 | Chr13 | 12/10/2025 | 9 | 100 | 10000 | 24 | 0 | 24 | 4.80E+05 | 0 | 0 |
| LRH570 | Chr13 | 12/10/2025 | 10 | 100 | 10000 | 14 | 1 | 15 | 3.00E+05 | 0.0666666667 | 6.66666667 |
| LRH570 | Chr13 | 12/10/2025 | 11 | 100 | 10000 | 22 | 25 | 47 | 9.40E+05 | 0.5319148936 | 53.19148936 |
| LRH570 | Chr13 | 12/10/2025 | 12 | 100 | 10000 | 15 | 2 | 17 | 3.40E+05 | 0.1176470588 | 11.76470588 |
| LRH570 | Chr13 | 12/10/2025 | 13 | 100 | 10000 | 22 | 0 | 22 | 4.40E+05 | 0 | 0 |
| LRH570 | Chr13 | 12/10/2025 | 14 | 100 | 10000 | 21 | 8 | 29 | 5.80E+05 | 0.275862069 | 27.5862069 |
| LRH570 | Chr13 | 12/10/2025 | 15 | 100 | 10000 | 13 | 1 | 14 | 2.80E+05 | 0.0714285714 | 7.14285714 |
| LRH570 | Chr13 | 12/10/2025 | 16 | 100 | 10000 | 12 | 2 | 14 | 2.80E+05 | 0.1428571429 | 14.28571429 |
| LRH570 | Chr13 | 12/10/2025 | 17 | 100 | 10000 | 14 | 0 | 14 | 2.80E+05 | 0 | 0 |
| LRH570 | Chr13 | 12/10/2025 | 18 | 100 | 10000 | 19 | 0 | 19 | 3.80E+05 | 0 | 0 |
| LRH570 | Chr13 | 12/10/2025 | 19 | 100 | 10000 | 28 | 0 | 28 | 5.60E+05 | 0 | 0 |
| LRH570 | Chr13 | 12/10/2025 | 20 | 100 | 10000 | 11 | 8 | 19 | 3.80E+05 | 0.4210526316 | 42.10526316 |
| LRH570 | Chr13 | 12/10/2025 | 21 | 100 | 10000 | 9 | 0 | 9 | 1.80E+05 | 0 | 0 |
| LRH570 | Chr13 | 12/10/2025 | 22 | 100 | 10000 | 18 | 6 | 24 | 4.80E+05 | 0.25 | 25 |
| LRH570 | Chr13 | 12/10/2025 | 23 | 100 | 10000 | 7 | 4 | 11 | 2.20E+05 | 0.3636363636 | 36.36363636 |
| LRH570 | Chr13 | 12/10/2025 | 24 | 100 | 10000 | 15 | 9 | 24 | 4.80E+05 | 0.375 | 37.5 |
| LRH507 | Chr14 | 12/10/2025 | 1 | 100 | 10000 | 144 | 0 | 144 | 2.88E+06 | 0 | 0 |
| LRH507 | Chr14 | 12/10/2025 | 2 | 100 | 10000 | 88 | 0 | 88 | 1.76E+06 | 0 | 0 |
| LRH507 | Chr14 | 12/10/2025 | 3 | 100 | 10000 | 90 | 0 | 90 | 1.80E+06 | 0 | 0 |
| LRH507 | Chr14 | 12/10/2025 | 4 | 100 | 10000 | 104 | 0 | 104 | 2.08E+06 | 0 | 0 |
| LRH507 | Chr14 | 12/10/2025 | 5 | 100 | 10000 | 122 | 0 | 122 | 2.44E+06 | 0 | 0 |
| LRH507 | Chr14 | 12/10/2025 | 6 | 100 | 10000 | 136 | 0 | 136 | 2.72E+06 | 0 | 0 |
| LRH507 | Chr14 | 12/10/2025 | 7 | 100 | 10000 | 110 | 0 | 110 | 2.20E+06 | 0 | 0 |
| LRH507 | Chr14 | 12/10/2025 | 8 | 100 | 10000 | 166 | 0 | 166 | 3.32E+06 | 0 | 0 |
| LRH507 | Chr14 | 12/10/2025 | 9 | 100 | 10000 | 110 | 0 | 110 | 2.20E+06 | 0 | 0 |
| LRH507 | Chr14 | 12/10/2025 | 10 | 100 | 10000 | 150 | 0 | 150 | 3.00E+06 | 0 | 0 |
| LRH507 | Chr14 | 12/10/2025 | 11 | 100 | 10000 | 146 | 0 | 146 | 2.92E+06 | 0 | 0 |
| LRH507 | Chr14 | 12/10/2025 | 12 | 100 | 10000 | 206 | 0 | 206 | 4.12E+06 | 0 | 0 |
| LRH507 | Chr14 | 12/10/2025 | 13 | 100 | 10000 | 5 | 115 | 120 | 2.40E+06 | 0.9583333333 | 95.83333333 |
| LRH507 | Chr14 | 12/10/2025 | 14 | 100 | 10000 | 102 | 0 | 102 | 2.04E+06 | 0 | 0 |
| LRH507 | Chr14 | 12/10/2025 | 15 | 100 | 10000 | 110 | 0 | 110 | 2.20E+06 | 0 | 0 |
| LRH507 | Chr14 | 12/10/2025 | 16 | 100 | 10000 | 104 | 0 | 104 | 2.08E+06 | 0 | 0 |
| LRH507 | Chr14 | 12/10/2025 | 17 | 100 | 10000 | 180 | 0 | 180 | 3.60E+06 | 0 | 0 |
| LRH507 | Chr14 | 12/10/2025 | 18 | 100 | 10000 | 366 | 0 | 366 | 7.32E+06 | 0 | 0 |
| LRH507 | Chr14 | 12/10/2025 | 19 | 100 | 10000 | 144 | 0 | 144 | 2.88E+06 | 0 | 0 |
| LRH507 | Chr14 | 12/10/2025 | 20 | 100 | 10000 | 142 | 0 | 142 | 2.84E+06 | 0 | 0 |
| LRH507 | Chr14 | 12/10/2025 | 21 | 100 | 10000 | 192 | 0 | 192 | 3.84E+06 | 0 | 0 |
| LRH507 | Chr14 | 12/10/2025 | 22 | 100 | 10000 | 178 | 0 | 178 | 3.56E+06 | 0 | 0 |

| Supplemental Table 5. Monosome replating test count data |  |  |  |  |  |  |  |  |  |  |  |
| --- | --- | --- | --- | --- | --- | --- | --- | --- | --- | --- | --- |
| Diploid parent strain | Chromosome | Date | Replicate | ul plated on YPD | Dilution plated on YPD | Monosomes | UPD | Total colonies on YPD | Population Total | Revertants:Monosomes | % Revertants |
| LRH507 | Chr14 | 12/10/2025 | 23 | 100 | 10000 | 158 | 0 | 158 | 3.16E+06 | 0 | 0 |
| LRH507 | Chr14 | 12/10/2025 | 24 | 100 | 10000 | 272 | 0 | 272 | 5.44E+06 | 0 | 0 |
| LRH500 | Chr15 | 12/10/2025 | 1 | 100 | 10000 | 110 | 0 | 110 | 2.20E+06 | 0 | 0 |
| LRH500 | Chr15 | 12/10/2025 | 2 | 100 | 10000 | 124 | 0 | 124 | 2.48E+06 | 0 | 0 |
| LRH500 | Chr15 | 12/10/2025 | 3 | 100 | 10000 | 102 | 1 | 103 | 2.06E+06 | 0.00970873786 | 0.9708737864 |
| LRH500 | Chr15 | 12/10/2025 | 4 | 100 | 10000 | 83 | 1 | 84 | 1.68E+06 | 0.0119047619 | 1.19047619 |
| LRH500 | Chr15 | 12/10/2025 | 5 | 100 | 10000 | 88 | 1 | 89 | 1.78E+06 | 0.01123595506 | 1.123595506 |
| LRH500 | Chr15 | 12/10/2025 | 6 | 100 | 10000 | 90 | 0 | 90 | 1.80E+06 | 0 | 0 |
| LRH500 | Chr15 | 12/10/2025 | 7 | 100 | 10000 | 96 | 1 | 97 | 1.94E+06 | 0.01030927835 | 1.030927835 |
| LRH500 | Chr15 | 12/10/2025 | 8 | 100 | 10000 | 108 | 0 | 108 | 2.16E+06 | 0 | 0 |
| LRH500 | Chr15 | 12/10/2025 | 9 | 100 | 10000 | 142 | 0 | 142 | 2.84E+06 | 0 | 0 |
| LRH500 | Chr15 | 12/10/2025 | 10 | 100 | 10000 | 152 | 14 | 166 | 3.32E+06 | 0.0843373494 | 8.43373494 |
| LRH500 | Chr15 | 12/10/2025 | 11 | 100 | 10000 | 146 | 0 | 146 | 2.92E+06 | 0 | 0 |
| LRH500 | Chr15 | 12/10/2025 | 12 | 100 | 10000 | 50 | 1 | 51 | 1.02E+06 | 0.01960784314 | 1.960784314 |
| LRH500 | Chr15 | 12/10/2025 | 13 | 100 | 10000 | 98 | 4 | 102 | 2.04E+06 | 0.03921568627 | 3.921568627 |
| LRH500 | Chr15 | 12/10/2025 | 14 | 100 | 10000 | 130 | 3 | 133 | 2.66E+06 | 0.0225639098 | 2.255639098 |
| LRH500 | Chr15 | 12/10/2025 | 15 | 100 | 10000 | 91 | 6 | 97 | 1.94E+06 | 0.0618556701 | 6.18556701 |
| LRH500 | Chr15 | 12/10/2025 | 16 | 100 | 10000 | 90 | 2 | 92 | 1.84E+06 | 0.02173913043 | 2.173913043 |
| LRH500 | Chr15 | 12/10/2025 | 17 | 100 | 10000 | 120 | 0 | 120 | 2.40E+06 | 0 | 0 |
| LRH500 | Chr15 | 12/10/2025 | 18 | 100 | 10000 | 132 | 0 | 132 | 2.64E+06 | 0 | 0 |
| LRH500 | Chr15 | 12/10/2025 | 19 | 100 | 10000 | 96 | 0 | 96 | 1.92E+06 | 0 | 0 |
| LRH500 | Chr15 | 12/10/2025 | 20 | 100 | 10000 | 130 | 0 | 130 | 2.60E+06 | 0 | 0 |
| LRH500 | Chr15 | 12/10/2025 | 21 | 100 | 10000 | 134 | 1 | 135 | 2.70E+06 | 0.007407407407 | 0.7407407407 |
| LRH500 | Chr15 | 12/10/2025 | 22 | 100 | 10000 | 104 | 8 | 112 | 2.24E+06 | 0.07142857143 | 7.142857143 |
| LRH500 | Chr15 | 12/10/2025 | 23 | 100 | 10000 | 134 | 1 | 135 | 2.70E+06 | 0.007407407407 | 0.7407407407 |
| LRH500 | Chr15 | 12/10/2025 | 24 | 100 | 10000 | 112 | 4 | 116 | 2.32E+06 | 0.03448275862 | 3.448275862 |
| Chr588 | Chr16 | 12/10/2025 | 1 | 100 | 10000 | 108 | 0 | 108 | 2.16E+06 | 0 | 0 |
| Chr588 | Chr16 | 12/10/2025 | 2 | 100 | 10000 | 118 | 0 | 118 | 2.36E+06 | 0 | 0 |
| Chr588 | Chr16 | 12/10/2025 | 3 | 100 | 10000 | 145 | 0 | 145 | 2.90E+06 | 0 | 0 |
| Chr588 | Chr16 | 12/10/2025 | 4 | 100 | 10000 | 146 | 0 | 146 | 2.92E+06 | 0 | 0 |
| Chr588 | Chr16 | 12/10/2025 | 5 | 100 | 10000 | 134 | 0 | 134 | 2.68E+06 | 0 | 0 |
| Chr588 | Chr16 | 12/10/2025 | 6 | 100 | 10000 | 130 | 0 | 130 | 2.60E+06 | 0 | 0 |
| Chr588 | Chr16 | 12/10/2025 | 7 | 100 | 10000 | 136 | 0 | 136 | 2.72E+06 | 0 | 0 |
| Chr588 | Chr16 | 12/10/2025 | 8 | 100 | 10000 | 142 | 0 | 142 | 2.84E+06 | 0 | 0 |
| Chr588 | Chr16 | 12/10/2025 | 9 | 100 | 10000 | 120 | 8 | 128 | 2.56E+06 | 0.0625 | 6.25 |
| Chr588 | Chr16 | 12/10/2025 | 10 | 100 | 10000 | 100 | 0 | 100 | 2.00E+06 | 0 | 0 |
| Chr588 | Chr16 | 12/10/2025 | 11 | 100 | 10000 | 121 | 0 | 121 | 2.42E+06 | 0 | 0 |
| Chr588 | Chr16 | 12/10/2025 | 12 | 100 | 10000 | 113 | 0 | 113 | 2.26E+06 | 0 | 0 |
| Chr588 | Chr16 | 12/10/2025 | 13 | 100 | 10000 | 252 | 0 | 252 | 5.04E+06 | 0 | 0 |
| Chr588 | Chr16 | 12/10/2025 | 14 | 100 | 10000 | 114 | 0 | 114 | 2.28E+06 | 0 | 0 |
| Chr588 | Chr16 | 12/10/2025 | 15 | 100 | 10000 | 100 | 0 | 100 | 2.00E+06 | 0 | 0 |
| Chr588 | Chr16 | 12/10/2025 | 16 | 100 | 10000 | 180 | 0 | 180 | 3.60E+06 | 0 | 0 |
| Chr588 | Chr16 | 12/10/2025 | 17 | 100 | 10000 | 125 | 0 | 125 | 2.50E+06 | 0 | 0 |
| Chr588 | Chr16 | 12/10/2025 | 18 | 100 | 10000 | 89 | 0 | 89 | 1.78E+06 | 0 | 0 |
| Chr588 | Chr16 | 12/10/2025 | 19 | 100 | 10000 | 85 | 0 | 85 | 1.70E+06 | 0 | 0 |
| Chr588 | Chr16 | 12/10/2025 | 20 | 100 | 10000 | 74 | 0 | 74 | 1.48E+06 | 0 | 0 |
| Chr588 | Chr16 | 12/10/2025 | 21 | 100 | 10000 | 87 | 0 | 87 | 1.74E+06 | 0 | 0 |
| Chr588 | Chr16 | 12/10/2025 | 22 | 100 | 10000 | 128 | 0 | 128 | 2.56E+06 | 0 | 0 |
| Chr588 | Chr16 | 12/10/2025 | 23 | 100 | 10000 | 167 | 0 | 167 | 3.34E+06 | 0 | 0 |
| Chr588 | Chr16 | 12/10/2025 | 24 | 100 | 10000 | 106 | 0 | 106 | 2.12E+06 | 0 | 0 |

**Supplemental Table 6.**
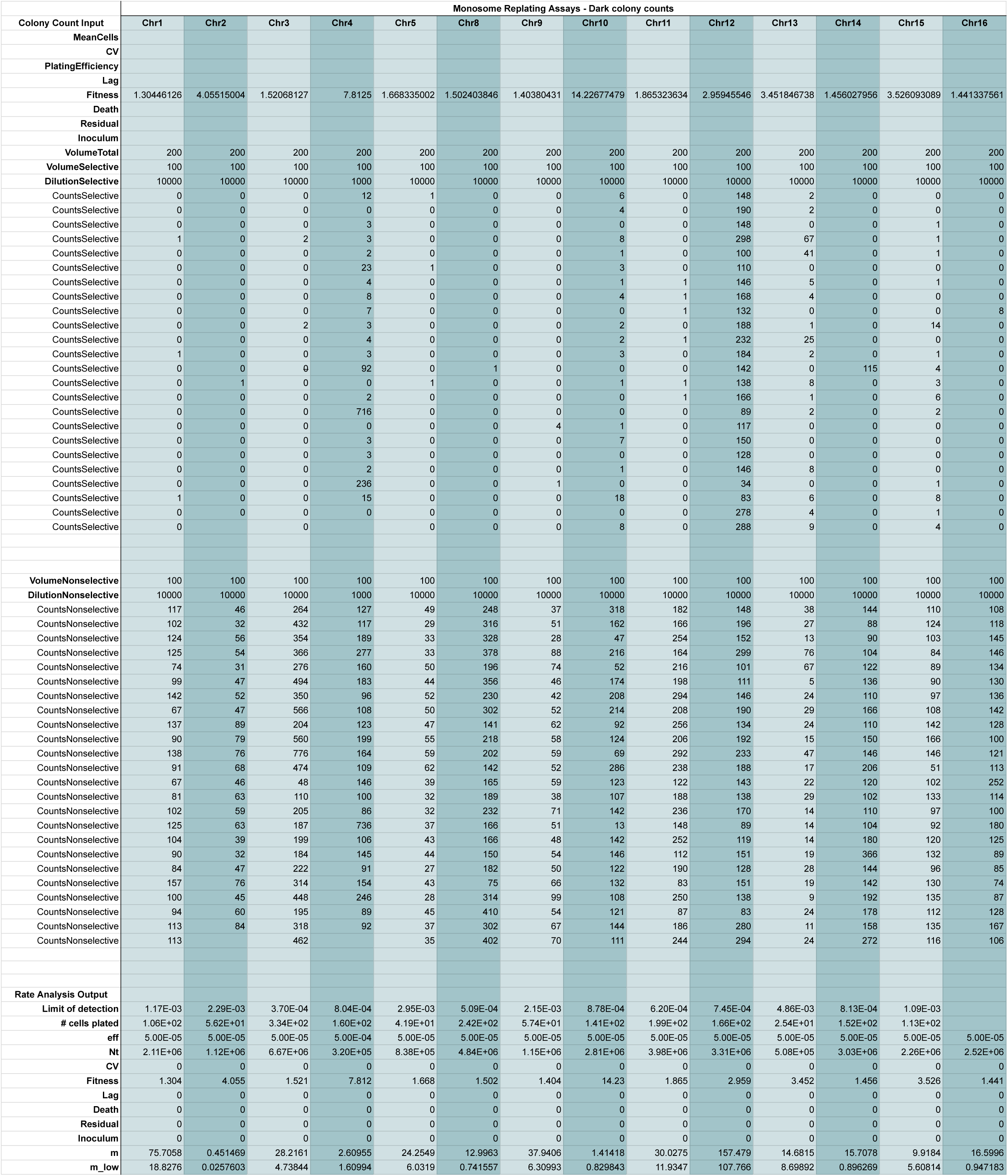
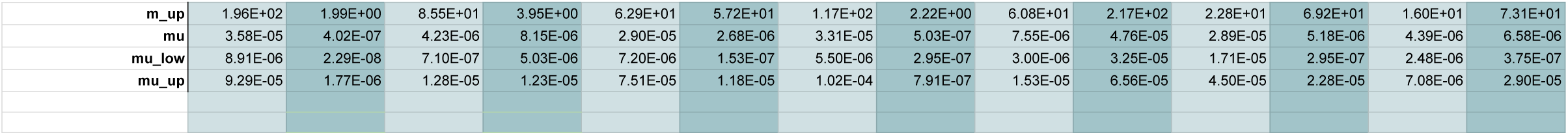
Monosome replating test rate analysis.

**Supplemental Table 7.**
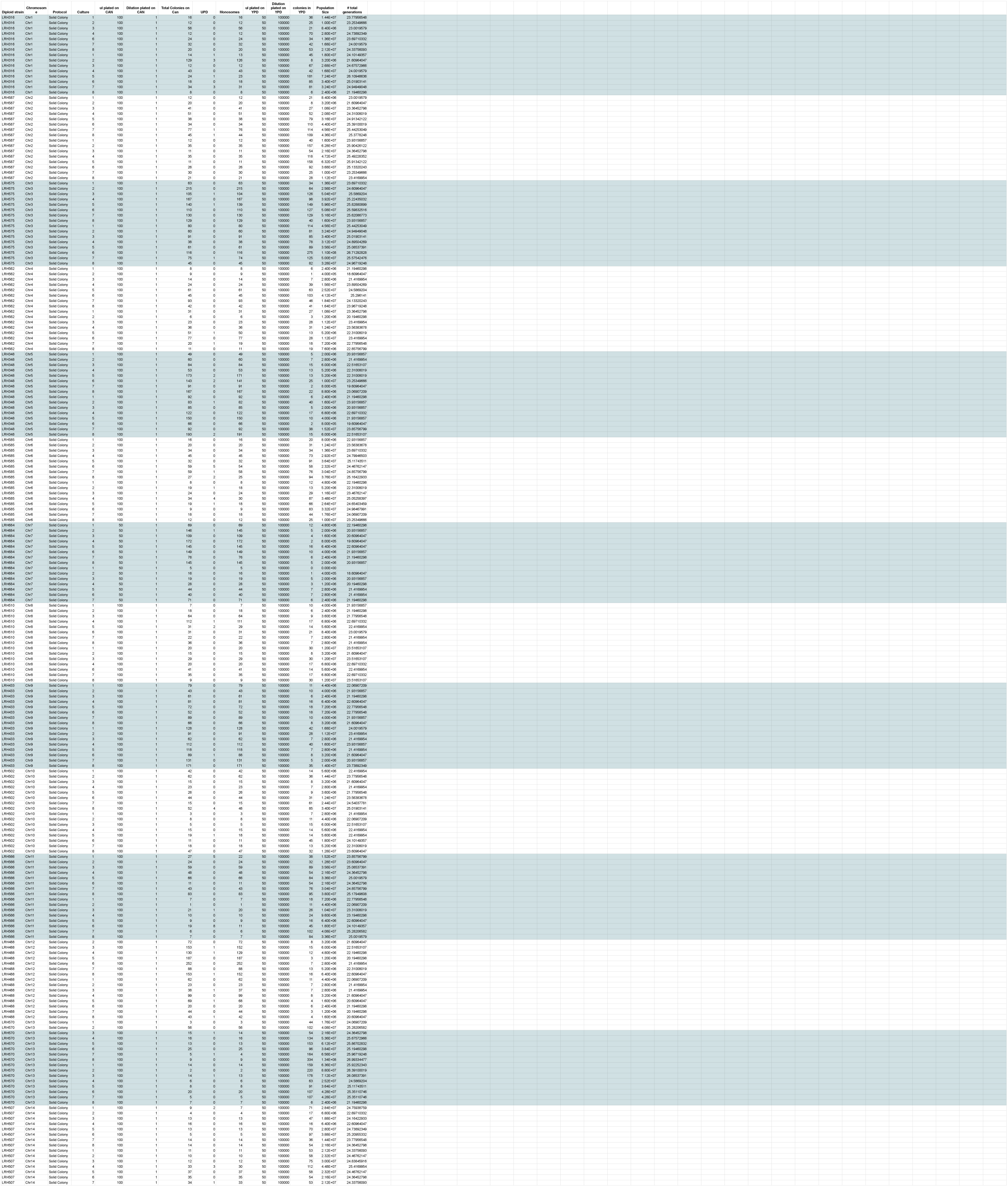
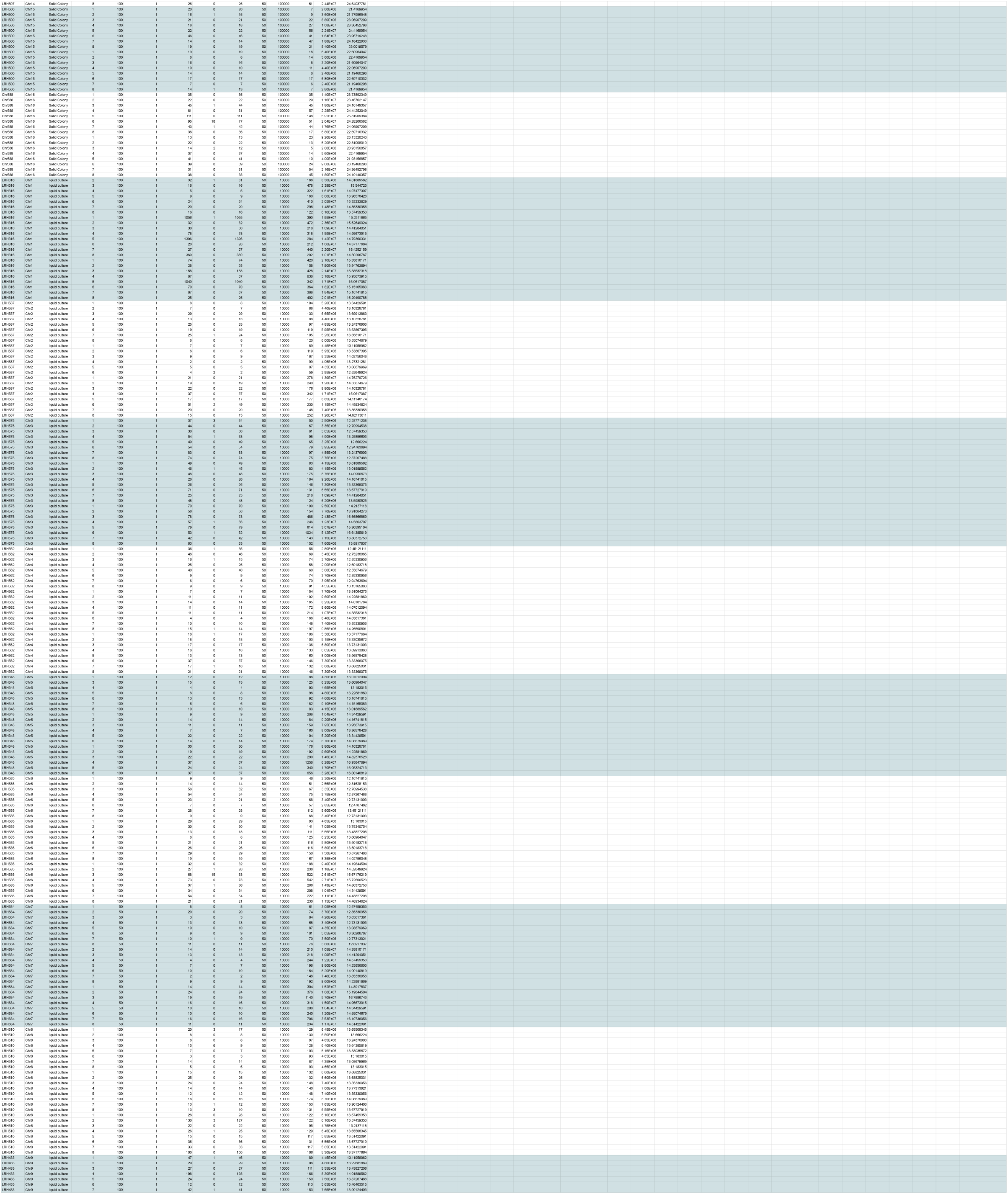
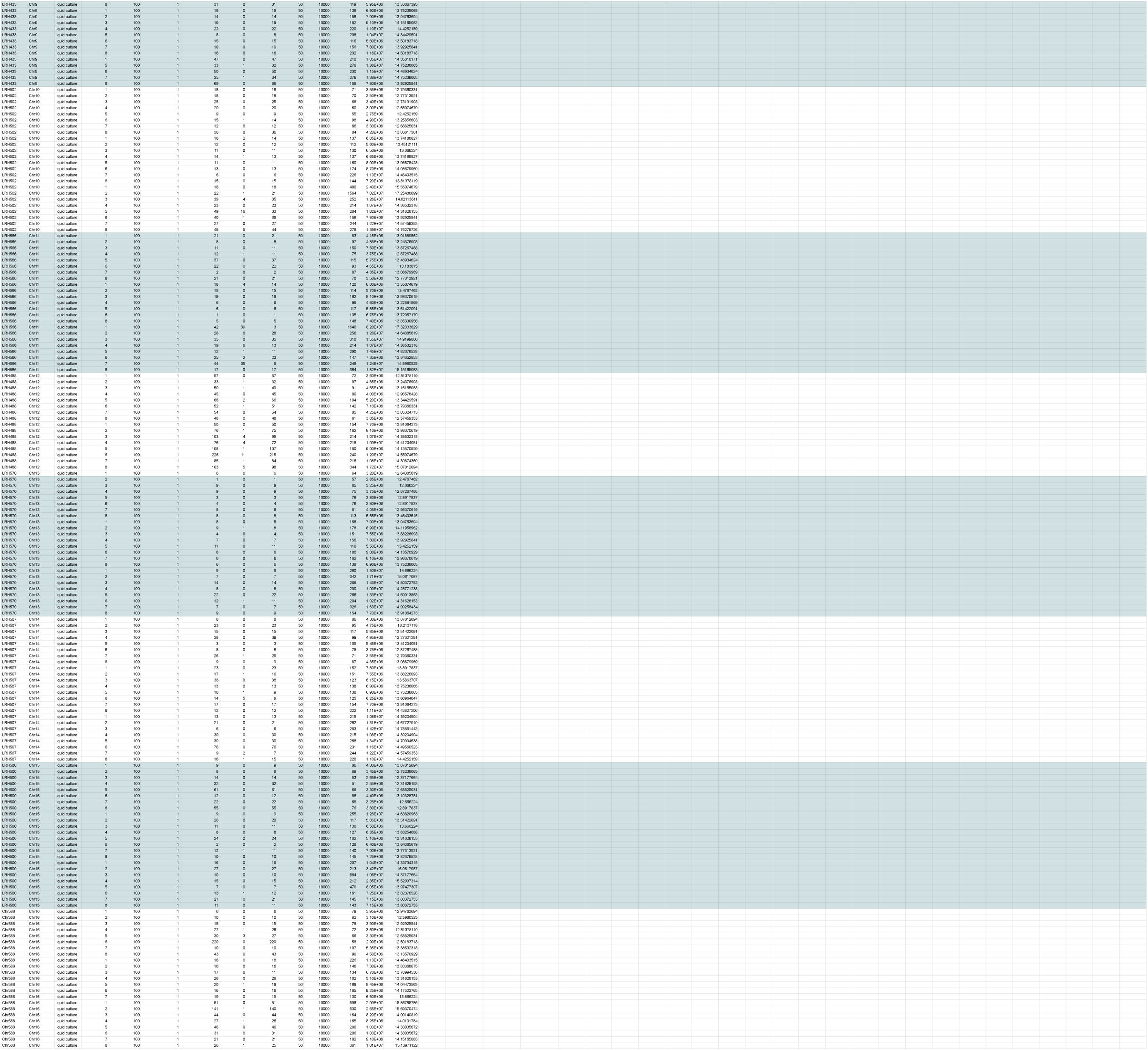
Canavanine fluctuation test colony count data.

**Supplemental Table 8.**
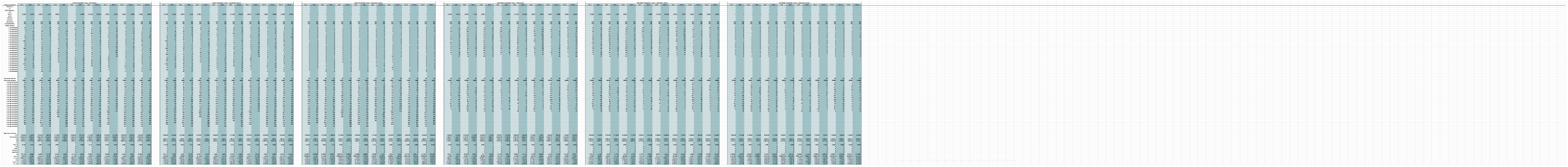
Canavanine fluctuation test rate analysis.

**Supplemental Table 9.**
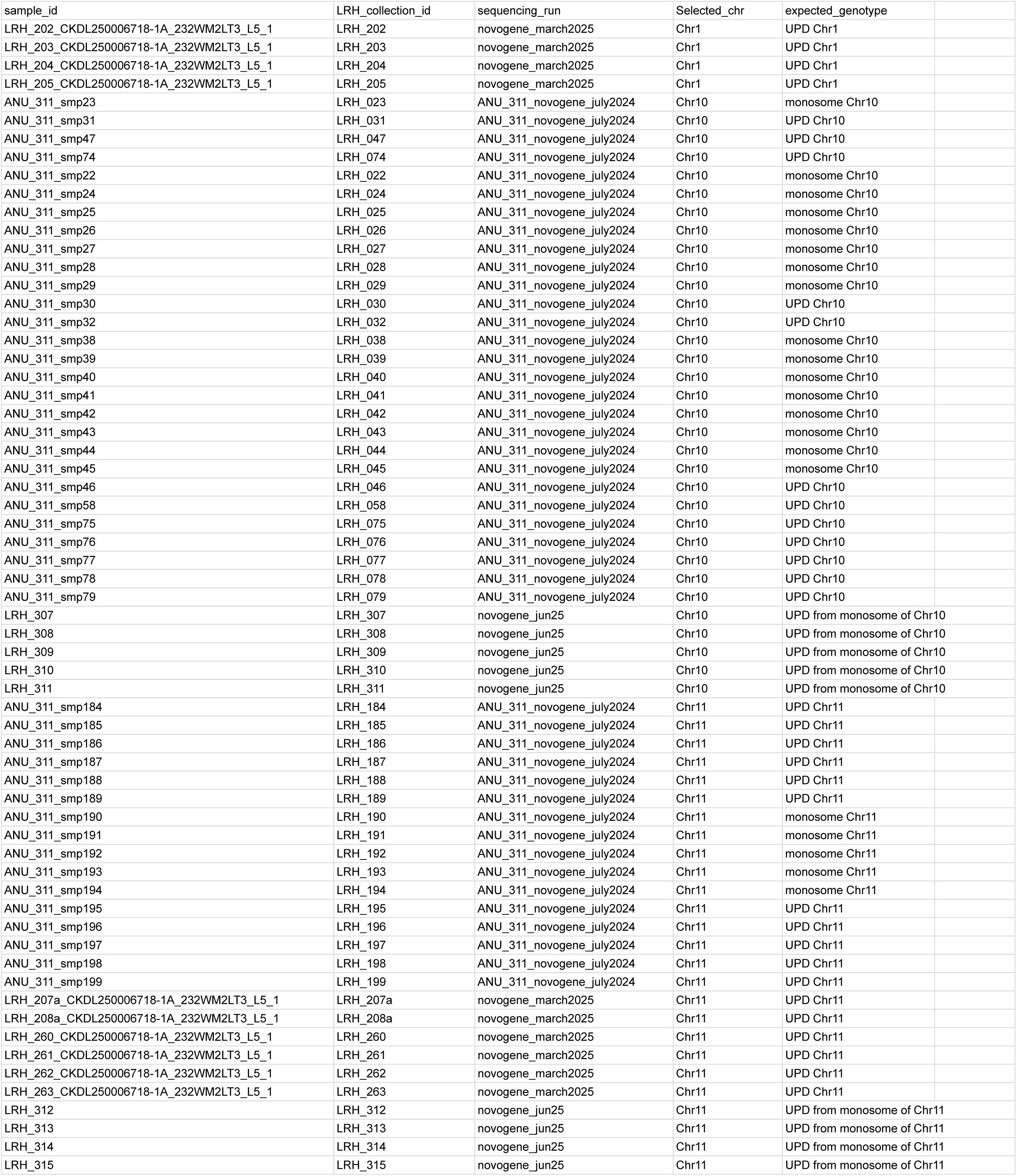
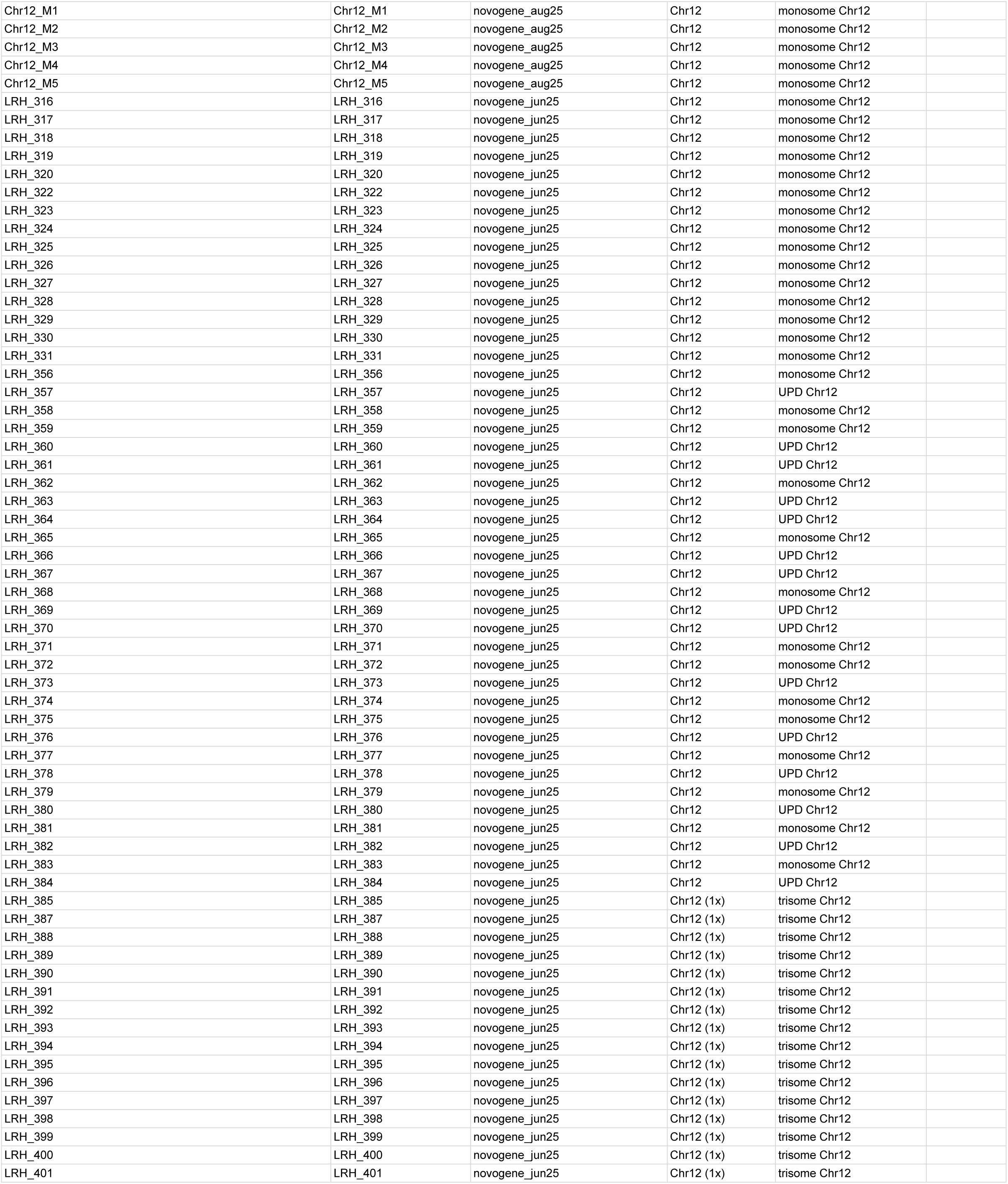
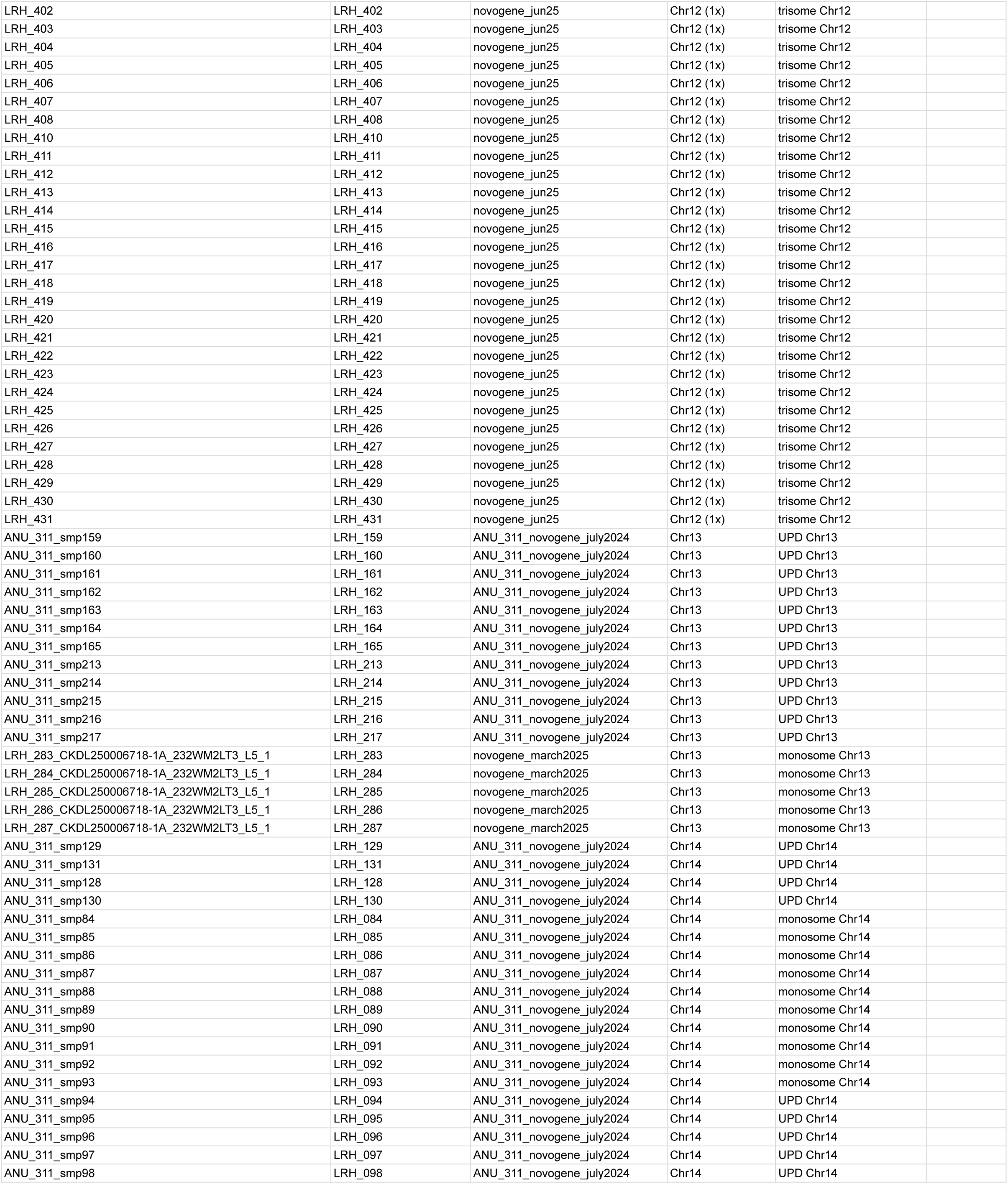
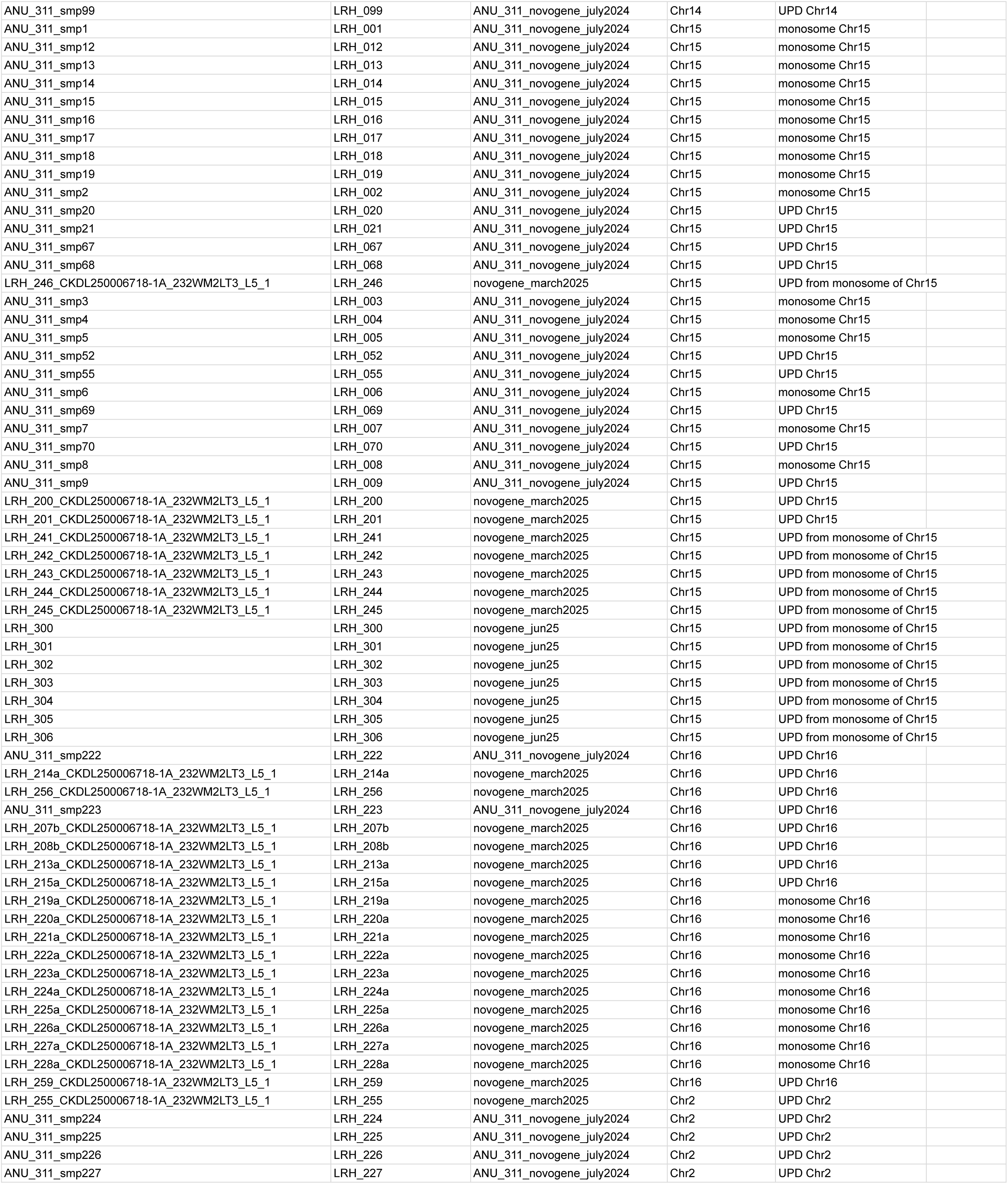
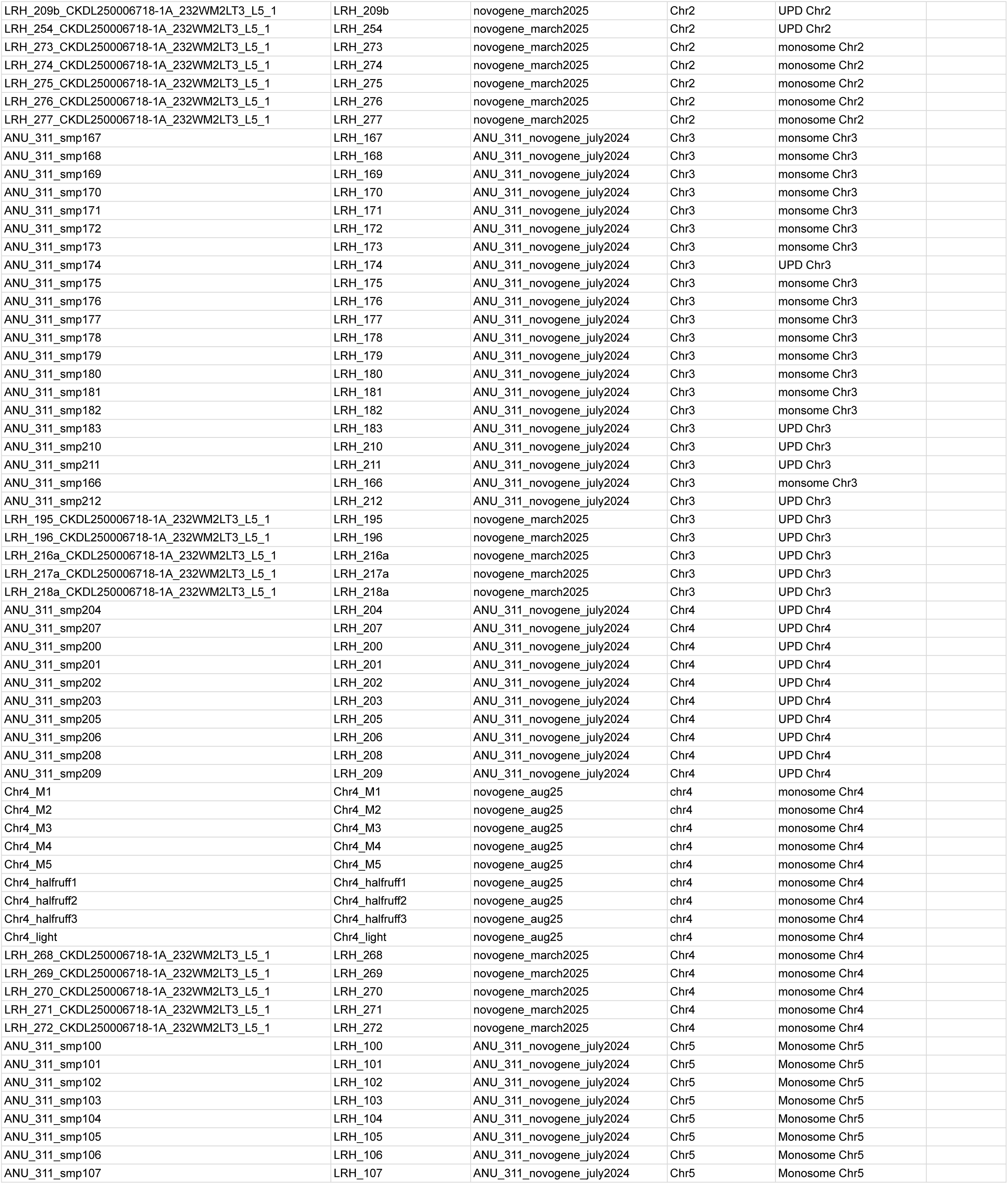
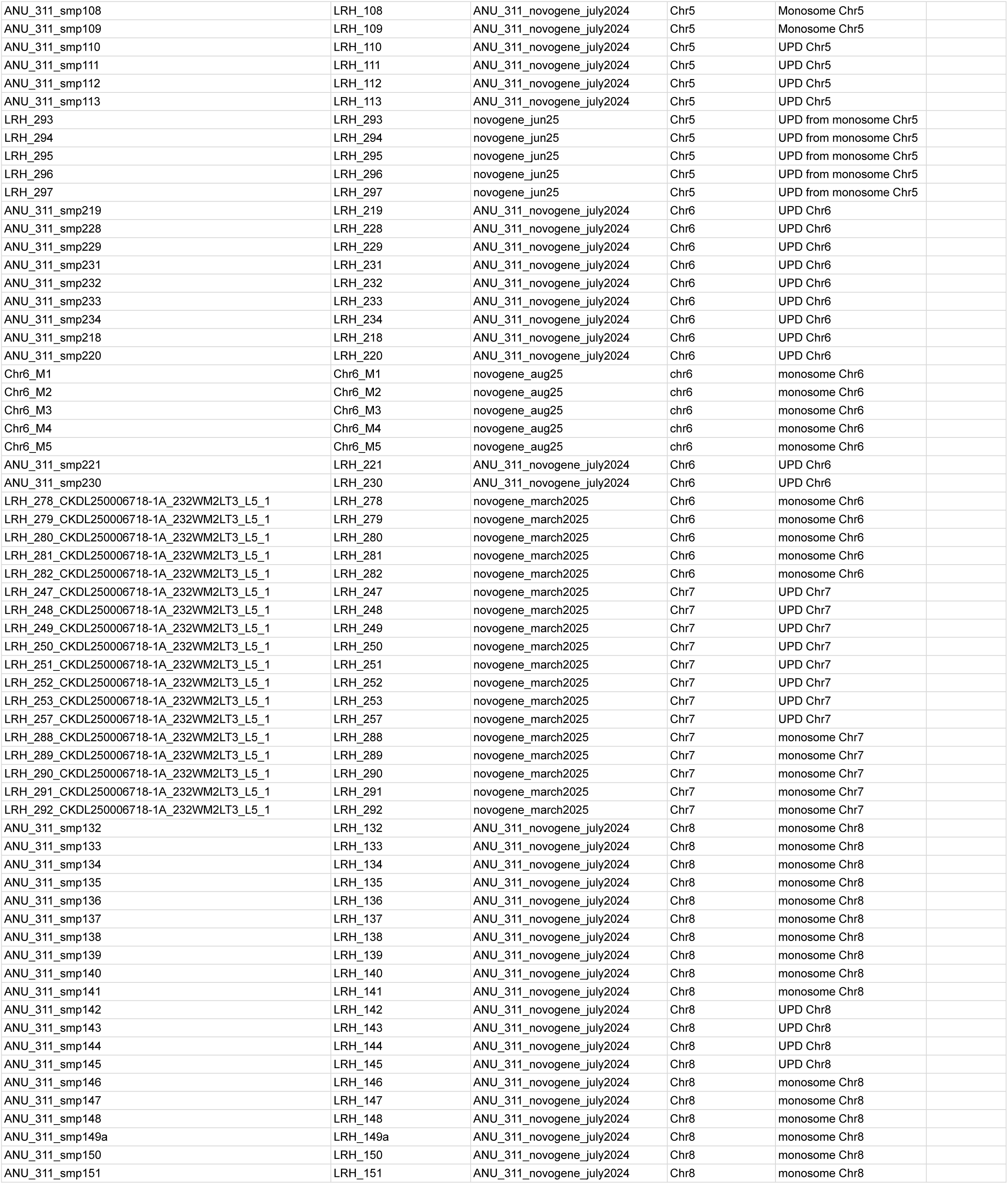
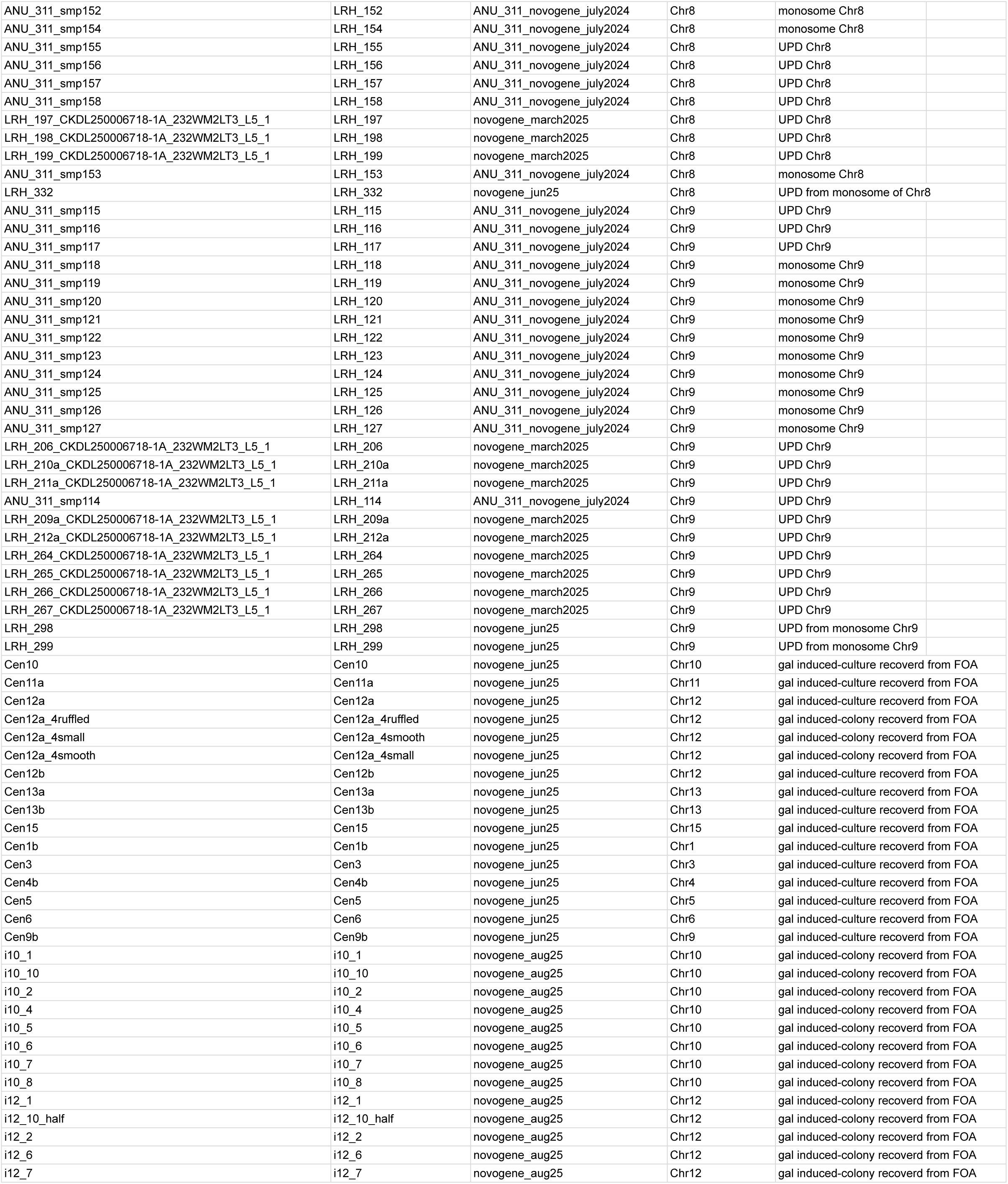
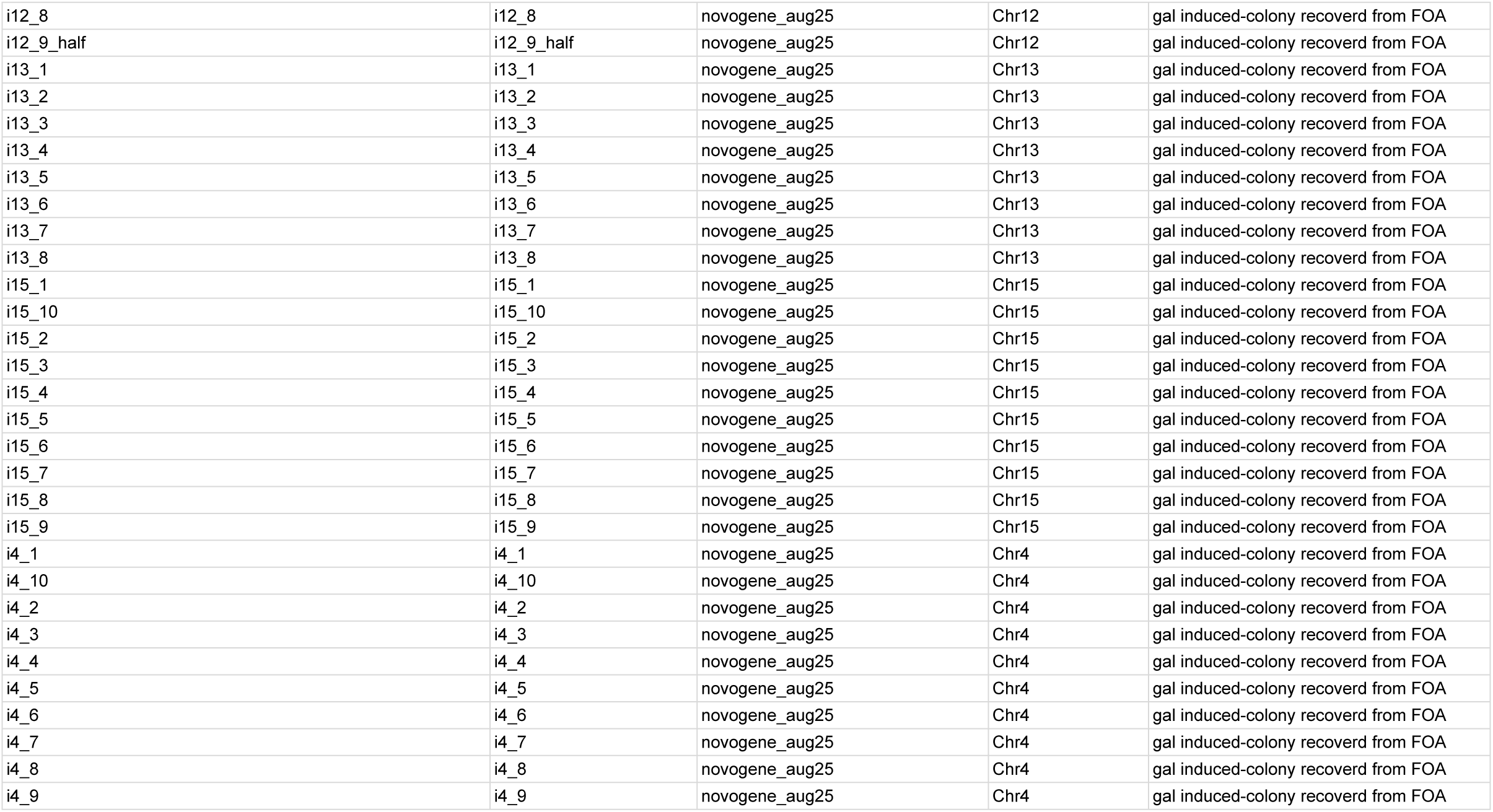
WGS sample information.

